# Persistent lytic bacteriophage infection as a novel strategy for exploitation of nutrient-limited host bacteria

**DOI:** 10.1101/2022.10.26.513911

**Authors:** Jack Dorling, Naima Nhiri, Andrés Corral-Lugo, Eric Jacquet, Paulo Tavares

## Abstract

Wild bacteria, from the open ocean to the gut, experience persistent nutrient limitation. This fundamentally affects bacterial physiology and metabolism and has profound impacts on their infection by bacterial viruses (bacteriophages). For virulent bacteriophages, which cannot enter a lysogenic state, this poses a problem for environmental persistence. Here we demonstrate that virulent bacteriophage SPP1 productively infects nutrient-limited stationary phase cultures of the Gram-positive bacterium *Bacillus subtilis*. Slow production and release of low numbers of infective viral particles resulted from a prolonged infection of the host population. Extensive culture lysis was greatly delayed, releasing additional viral particles and promoting fresh infections of bacterial survivors. Induced overproduction of cell surface bacteriophage receptor YueB, compensating for its scarcity in stationary phase, expedited infection dynamics under nutrient-limiting conditions, but did not change overall infection productivity. The temporal program of SPP1 gene expression differed from exponential phase, consistent with a prolonged, persistent mode of infection. Reduced expression of genes coding viral structural proteins correlated with the low yield of infectious particles. Importantly, exogenous influx of the carbon source maltose enhanced viral particle production. Our results uncover a novel adaptive strategy of a lytic phage for productive infection of nutrient-limited bacterial populations through persistent, exhaustive infection.

## Introduction

In contrast to the highly-controlled, near ideal conditions under which most laboratory-based bacteriophage (phage)-bacteria infection studies are conducted, the growth and proliferation of wild bacteria is primarily restricted by limited or fluctuating carbon and energy supplies [1,2]. Availability of other nutrients [3,4] and physico-chemical environmental factors also limit bacterial growth [2,5]. These conditions are observed not only in aquatic and soil ecosystems but also in environments such as the mammalian gut where intense competition limits nutrient availability [6]. Consequently many environmental bacteria experience severe nutrient limitation and occupy a physiological state of slow or arrested metabolism known as ‘starvation-survival’ [1,7]. Growth rates in both aquatic and soil ecosystems are accordingly low [8–10] but bacterial densities may reach 10^5^-10^6^ colony forming units (CFU) ml^-1^ [5] or 10^9^ CFU g^-1^ [11], in each respective environment, demonstrating the adaptation of bacteria to such oligotrophic conditions.

Phages depend upon their hosts’ metabolism for multiplication [12,13]. However, global phage abundance is estimated to be higher than 10^30^ [14], even with considerable estimated environmental phage particle decay rates [15,16]. Thus, as bacteria have adapted to starvation-survival lifestyles, phages have conceivably adapted to exploit starving hosts. Such adaptive strategies remain largely unstudied in spite of their critical importance for phage persistence in natural ecosystems.

Phages exhibit varied infection strategies [13,15-19], ranging from lytic to lysogenic cycles. In lysogenic bacteria, chromosomally-integrated or episomal temperate phage genomes are maintained and propagated within the bacterial population [16]. Entry into lysogenic states and concomitant silencing of most phage gene expression is largely determined by host metabolic state upon infection [16,20]. However, strictly lytic (i.e. unable to establish a lysogenic state) phages persist in soil and aquatic environments. This raises the fundamental questions of if and how lytic phages are able to overcome the challenges of infecting starving hosts. Indeed, it is often implicitly assumed that such phages require exponentially growing bacteria in order to stage productive infections.

To address this dearth of information, we investigated whether a strictly lytic phage could productively exploit a non-growing, nutrient limited host. To do so, we employed the model Gram-positive soil bacterium *Bacillus subtilis*, which displays numerous adaptations to nutrient limiting conditions [21,22] and its lytic siphovirus Subtilis Phage Pavia 1 (SPP1) [23]. SPP1 infection of *B. subtilis* starts with reversible adsorption to cell wall teichoic acids, irreversible adsorption to the protein YueB [24], and transfer of naked phage DNA into the bacterial cytoplasm [25,26]. These processes depend on the presence of Ca^2+^ and host membrane potential [27,28]. Under optimised laboratory conditions phage DNA is delivered to the cell within the first 3min [28], followed by early gene expression and genome replication [29,30]. Late gene expression begins after 10-12min and virion assembly ensues [31]. After ∼30-60min host cells lyse, liberating ∼200 viral particles per cell [32]. Previous work suggested that SPP1 may only be able to infect growing bacteria [33].

Here we report the discovery of a low productivity stationary phase (SP) infection strategy of *B. subtilis* by SPP1. The greatly prolonged infection of most of the bacterial population, the low yield of infectious particles (or virions) and gene expression patterns differ significantly from infection of exponentially growing bacteria. Such a persistent infection mode of nutrient-limited *B. subtilis* bacteria represents a resiliency strategy to promote sustainability of lytic phage populations under conditions that do not sustain host bacterial growth.

## Materials and Methods

### Bacteria and phage strains

Bacterial and phage strains are in **Table S1**. *B. subtilis* 168 strain 1A1 [34] (*Bs*^*wt*^) was the wild-type strain. Strain YB886 [35] was used for phage amplification and titration. *B. subtilis* strain *Bs*^*Pspac::yueB*^ over-expresses the SPP1 receptor YueB in the presence of IPTG. Either wild-type SPP1 (SPP1^*wt*^) or SPP1 expressing an mNeonGreen reporter (SPP1^*mNeonGreen*^) were used for infection experiments.

*Bs*^*Pspac::yueB*^ and SPP1^*mNeonGreen*^ were constructed as described in **Supplementary Material and Methods**. Primers used are in **Table S2**.

### Bacteria growth and phage infection

Bacteria were cultured in LB medium at 37°C unless otherwise stated. Experiments were either performed in flasks or in 96-well plates in a Tecan plate reader. Bacterial CFU were enumerated by serial dilution and plating. CFU originating from spores were enumerated by the same method after heating the sample to 80°C for 20 minutes. Phage infections were made using 2×10^9^PFU ml^-1^ with the addition of 10mM CaCl_2_. Phages were enumerated by serial dilution and titration in semi-solid agar. For free phage populations only sample supernatant was titrated [33]. For total phage populations the entire sample was mixed with chloroform, left on ice for >2h and titrated. SPP1 irreversible adsorption (IA) was measured as described in [24].

### Microscopy

For microscopy, bacteria were placed on a water-agarose pad and imaged using an Axio Observer Z1 microscope with a 63x oil immersion lens (Zeiss; Marly Le Roi, France). DAPI and GFP filter sets were used to image TMA-DPH and mNeonGreen/AlexaFluor 488 fluorophores, respectively. Post-capture image processing was carried out using Fiji [36]. MicrobeJ [37] was used to extract morphological and fluorescence data from microscopy images. After background fluorescence correction infected cells were identified as those exhibiting mNeonGreen fluorescence intensity greater than +4·SD above the mean fluorescence of non-infected cells. This was verified by manual counting (**Supplementary figure 1**).

### RNA and protein methods

For RNA and protein isolation, samples were mixed with sample stop buffer containing 100mM NaN_3_, supernatant thoroughly removed, pellets frozen in liquid nitrogen and stored at -80°C. RNA was extracted with a ‘NucleoSpin RNA Mini’ RNA purification kit (Macherey-Nagel). Samples were purified twice with an extra DNase digestion step between purifications. RNA was quantified using a Nanodrop One spectrophotometer (ThermoFisher Scientific) and RNA integrity verified using an Agilent 2100 bioanalyzer (Agilent Technologies). 200ng total RNA was then used to produce cDNA. Quantitative real-time PCR (qRT-PCR) was performed on a QuantStudio 12K Flex Real-Time PCR System (Life Technologies) with a SYBR green detection protocol. Data normalisation was performed with five *B. subtilis* reference genes. Relative gene expression ratios were determined using the ΔΔCt method with a Ct limit of 33 cycles.

Protein was extracted as described previously [38]. Approximate protein concentrations were verified by intensity of Coomassie staining of gels and Western-blot detection of gp11 was carried out with a rabbit polyclonal α-gp11 antibody [39].

### Quantitative and statistical data analyses

Analyses were performed using R version 3.6.3 [40]. Growth and lysis rates were derived using the smooth.spline and predict packages [40]. All statistical analyses involving more than 2 independent groups were performed using linear modelling. The package lm was used to fit general linear models (LM’s) to normally distributed, constant variance (or transformed) data, and glm to fit generalised linear models (GLM’s) to untransformed data requiring different error structures or link functions. Piece-wise regression analyses breakpoints were performed using the segmented R package [41]. Multiple comparisons were made using either multcomp [42] or emmeans [43] packages. Model predictions for visual representation were generated using the predict package. Two-tailed t-tests or Wilcoxon rank sum tests were performed using appropriate R packages [40]. Pearsons correlation coefficients were calculated using the cor package [40].

Further details of Materials and Methods are provided in **Supplementary Material and Methods**.

## Results

### Assessment of onset nutrient limitation during the *B subtilis* growth cycle

Growth and morphological characteristics of *Bs*^*wt*^ were studied during its growth cycle in LB medium to determine the timing of growth arrest and nutrient limitation. Cell size was used as a proxy for nutrient limitation [44–46]. Detectable growth halted from ∼5h onward (**Supplementary figure 2a, inset**), clearly delimiting periods of growth and non-growth. Transition phase (TP) was designated as the period between 4-6h, preceded by exponential phase (EP) and followed by SP. This determined time-points to study phage infection during different host growth phases. We observed a decline in SP OD_600_ between ∼9 and 30h (**Supplementary Fig. 2a**), though viable cell counts indicated no significant cell death throughout this period (**Supplementary Fig. 2b**). This was accompanied by a reduction of cell length with culture age (**Supplementary figure 2c**) while cell width varied little resulting in a concomitant reduction of surface area and volume. By 10h post-inoculation ≥97.1% of cells had reached ‘minimal’ cell dimensions as demonstrated by their similarity to those at 30h (**Supplementary figure 2d; Supplementary Data Analysis**). Cell length reduction thus explained the reduction of culture OD_600_ during SP without loss of cell viability. As bacteria adopt a smaller size under nutrient poor conditions [44,45], we concluded that nutrient limitation increased with time, and that by 10h post-inoculation cells could be considered non-growing and nutrient-limited.

### SPP1 infection during different *B. subtilis* growth cycle stages

The zone of clearance within SPP1 phage plaques does not grow detectably after reaching a defined size, though lysis halo formation is observed (**Supplementary figure 3**). We hypothesised that this results from an inability of SPP1 to infect bacteria in SP. We thus decided to examine the efficiency of SPP1 infection at selected time-points during the *B subtilis* growth cycle in liquid medium (**Fig. 1a**). Infection efficiency, assessed by culture lysis (**Fig. 1b**) and free PFU production (**Fig. 1c**), gradually decreased with bacterial growth rate phase. Indeed, EP and early transition phase (eTP) infected cultures lost OD_600_ more rapidly over time than non-infected cultures, while late transition phase (lTP) or SP infected cultures did not (**Fig. 1b; Table S3)**. Only EP and eTP infections resulted in net PFU production (**Fig. 1c**). These initial experiments were limited to a 2h window post-infection (p.i.). After 2h of infection in SP, little lysis had occurred but the number of free infectious phages had clearly decreased below the input level (**Fig. 1c**). This led us to the hypothesis that phage adsorption and inactivation had occurred and that infection may occur but more slowly than under nutrient-replete conditions. Therefore, we infected SP cells as previously and followed infection for 18h. During this longer p.i. period slow lysis of the infected culture occurred (**Fig. 1d**) and by 18h p.i. a ∼5-fold net increase in free infectious phage particles relative to the input level was observed (**Fig. 1e**).

**Fig. 1.**
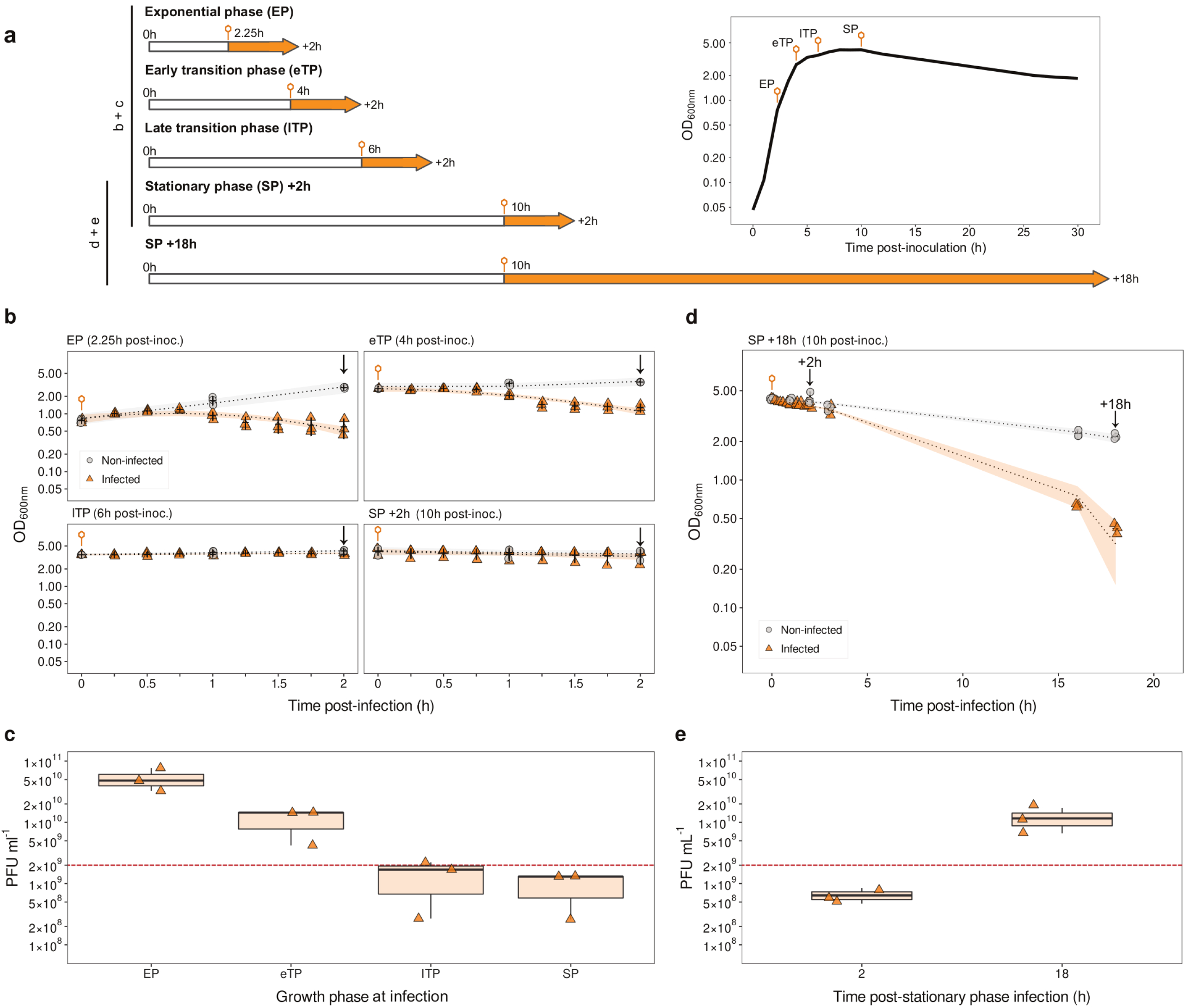
SPP1 infection of *B. subtilis* during exponential, transition and stationary phase. **a** Infection experiment setup. White bars represent growth of uninfected cultures up until infection (orange phage symbols, time-point adjacent) during the bacterial growth phase indicated on the left, as defined in **Supplementary figure 2**. Following infection, both one non-infected culture and one infected culture were then followed for the duration shown on the right of each arrow. The timing of each infection during the growth of *B, subtilis* is marked on the growth curve displayed to the right of the panel. **b** OD_600_ curves of *Bs*^*wt*^ grown in flasks in LB medium at 37°C and infected with an input of 2×10^9^ pfu ml^-1^ of SPP1^*wt*^ (orange triangles) at the time-points indicated in **a**. Infection was monitored for 2 h. Control growth curves of non-infected bacteria are in grey. **c** Titre of free phage PFU at the end of each infection in **b**. The dashed line indicates the input of PFU used for infection. **d** OD_600_ curve of *Bs*^*wt*^ infected in SP for 18h as represented schematically in **a. e** Titre of free phage PFU after 2 and 18h p.i. in SP. Data in **b**-**e** are from three independent biological replicates. Linear regression models fit to the data for statistical analysis in **b** and **d** are represented graphically by dotted lines (model fit) and shaded areas (95% confidence intervals). The mean (horizontal lines) and standard deviation (vertical bars) are displayed in **b** and **d**. The median, upper and lower quartiles (boxes) and the limits of the data (whiskers) of the data are given in **c** and **e**. Additional data and analyses related to these experiments are provided in **Table S3** and **Supplementary Data Analysis**. Colour version of figure available online.

### Dynamics of SPP1 infection of *B subtilis* during stationary-phase

We then continuously monitored, over a 30h period, the infection of *Bs*^*wt*^ by SPP1^*wt*^ during different growth phases using a plate-reader-based setup. Lysis was evident in all infected cultures (orange triangles in **Fig. 2a**) relative to non-infected controls (grey circles in **Fig. 2a**). Infection duration increased with culture age (**Fig. 2b**). Cultures infected early during TP displayed multiphasic lysis behavior while late in TP and SP a single phase of slow lysis preceded a single more rapid lysis phase (**Fig. 2a**).

**Fig. 2.**
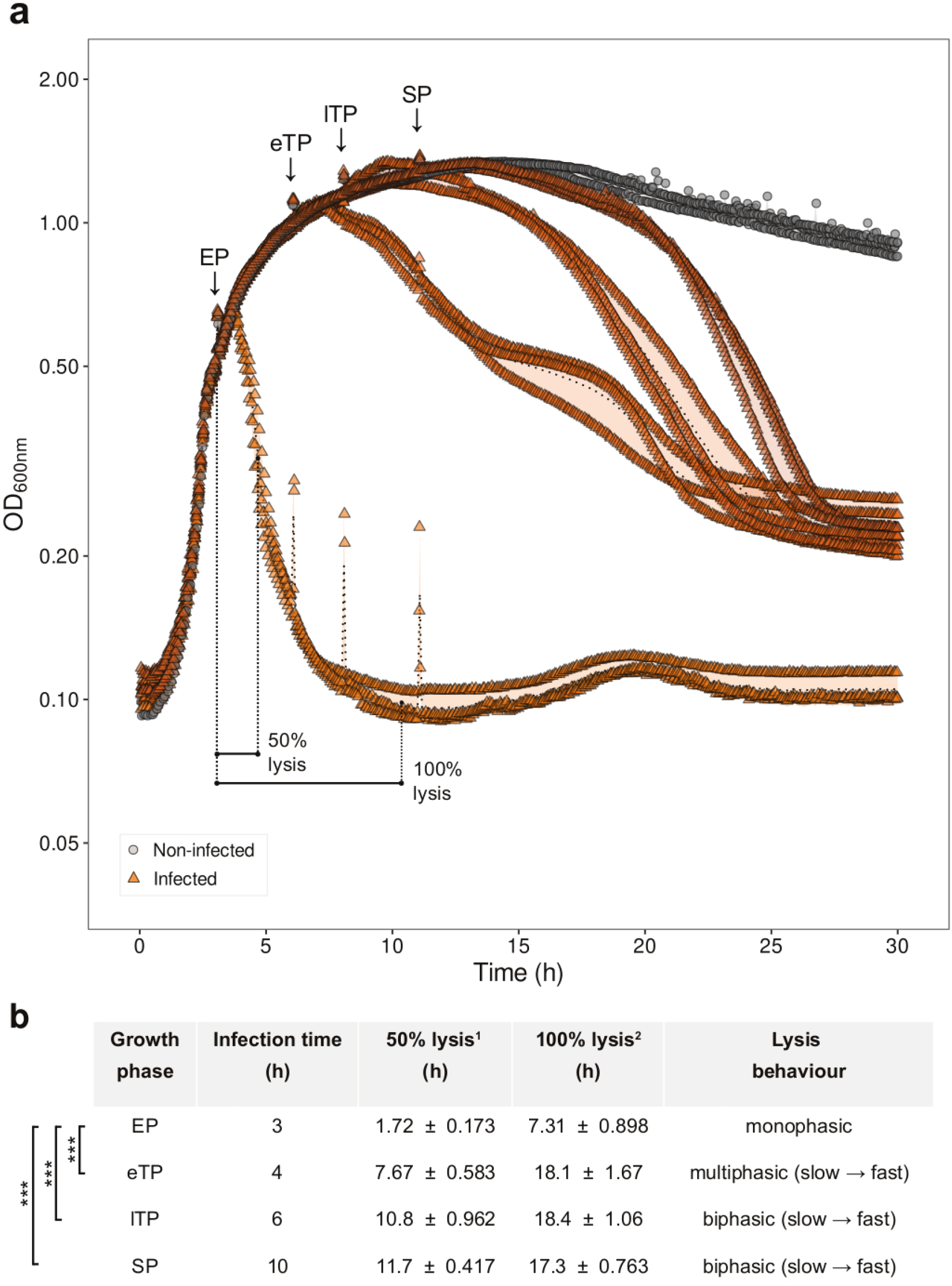
Long-term observation of growth and lysis of *B. subtilis* infected during exponential, transition and stationary phase. **a** OD_600_ curves of *Bs*^*wt*^ cultures grown in a 96-well plate and monitored continuously in a plate reader. Bacteria were infected with 2×10^9^ pfu ml^-1^ of SPP1^*wt*^ at different phases of the *B. subtilis* growth cycle (orange triangles). Infection times differ slightly from experiments performed in flasks (see **Fig. 1**) to take into account the lower growth rate of cultures in 96-well plates. Sharp peaks in OD_600_ are due to settling of cells when plates were removed for infection with SPP1. Control growth curves of non-infected bacteria are in grey. An example of the times taken to reach 50% of the maximum OD_600_ (50% lysis) and for lysis to stop (100% lysis) in EP infection is displayed (see **Supplementary Material and Methods**). **b** Lysis behavior features extracted from the data in **a**. Tukey’s honest significant difference contrasts (THC’s) are displayed to the left of the table and are identical for both data columns. Statistical significance levels; *** p < 0.001. Non-significant differences are not shown. Data are from three independent biological replicates. Means (dotted line in **a**) and standard deviations (shaded areas in **a**) are displayed in both panels. Data and analyses related to these experiments are presented in **Table S3** and **Supplementary Data Analysis**. Colour version of figure available online.

We then used a flask-based setup to simultaneously monitor OD_600_, viable cell counts, phage yield and infection prevalence at the single-cell level, using the reporter phage SPP1^*mNeonGreen*^. As expected, slow, but statistically significant lysis was evident in the first 10h of infection (**Fig. 3a**; analysis of covariance; F_1,12_^time:infection_status^ = 71.9, p < 0.001) and was accompanied by a significant reduction in CFU recovered from infected cultures following phage-induced lysis (green triangles in **Fig. 3b**). In contrast, no differences were found between infected and non-infected cultures in the proportion of CFU originating from spores (diamonds and squares in **Fig. 3b**, respectively; ordinate on the right), which are resistant to SPP1 infection. Sporulation frequency increased over time in all cultures reaching ∼2.5-2.8% by 40h post-inoculation. This indicated that SPP1 infection of the bacterial population did not appreciably affect the developmental decision of *B. subtilis* to sporulate under nutrient limitation [47]. Immediately after infection, free phage numbers dropped 3.4 ± 0.3-fold relative to the input due to their engagement in infection. After a period of ∼2h, phage counts increased at a relatively steady rate from 2 to 10h p.i. in both free and total phage populations (**Fig. 3c, Table S3**). This coincided with steady culture lysis in the first 10h p.i. (**Fig. 3a**). Between 10h and 30h p.i. further phage multiplication had occurred with a convergence in the number of phages found in free and total populations during this period (**Fig. 3c**, Wilcoxon rank sum test^30h p.i.^; W = 6, p = 0.686). This indicated release of most intracellular PFU into the extracellular environment and coincided with extensive culture lysis.

**Fig. 3.**
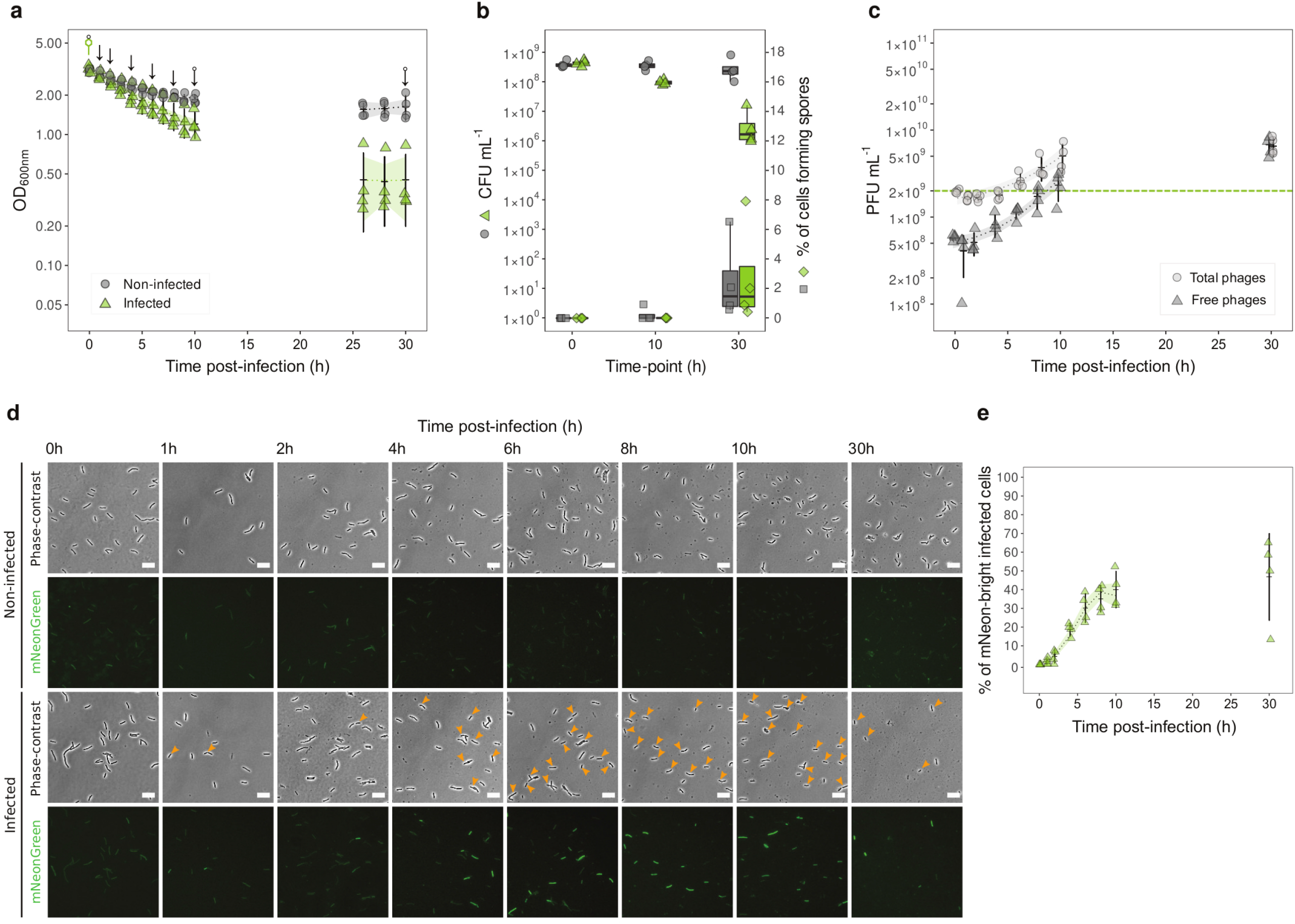
SPP1 infection dynamics during prolonged stationary phase infection of *B. subtilis*. *Bs*^*wt*^ bacteria grown in flasks with LB medium at 37°C were infected with 2×10^9^ pfu ml^-1^ of SPP1^*mNeonGreen*^ at 10h post-inoculation (green phage symbol in **a**). Bacterial density (OD_600_ in **a** and CFU ml^-1^ in **b**), total and free phage (PFU ml^-1^) (**c**), and the number of visibly infected bacteria (**d, e**) were monitored until 30h p.i., in parallel. The number and percentage of CFU originating from spores was also quantified (grey squares for non-infected culture and green diamonds for infected culture in **b**; ordinate on the right). Individual infected cells were counted at each time-point by microscopy (orange arrowheads in **d**, identifying bright cells in the mNeonGreen channel) and the percentage of infected cells in the infected culture determined (**e**). The white scale bars represent 10μm. Sample sizes were as follows; 0h; n = 510, 1h; n = 630, 2h; n = 816, 4h; n = 658, 6h; n = 967, 8h; n= 780, 10h; n = 764, 30h; n = 241. Data are from three independent biological replicates. Linear regression models fit to the data for statistical analysis in **a, c** and **e** are represented graphically by dotted lines (model fit) and shaded areas (95% confidence intervals). The mean (horizontal lines) and standard deviation (vertical bars) are displayed in **a, c** and **e**. The median, upper and lower quartiles (boxes) and the limits of the data (whiskers) of the data are given in **b**. Data and analyses related to these experiments are detailed in **Table S3** and **Supplementary Data Analysis**. Colour version of figure available online.

In parallel, we imaged infected cultures to examine infection propagation within the bacterial population. To do so we used a phage engineered to express the fluorescent reporter mNeonGreen under the control of the SPP1 *P*E2 early promoter. Control experiments conducted in EP, where infection prevalence is known to be ∼100%, showed that SPP1^*mNeonGreen*^ robustly reported infection in only ∼72% of the bacterial population at 1h p.i. (**Supplementary Fig. S4**). This is due to the time required for mNeonGreen folding and fluorophore maturation, leading to some underestimation of the number of infected cells at defined time points. This caveat in mind, we documented a very slow initial increase in the proportion of visibly infected, mNeonGreen-bright, cells in SP (**Fig. 3d,e**) before a more rapid increase until observations were halted at 10h p.i. (**Fig. 3d,e**). Interestingly, despite the major release of intracellular phage particles between 10-30h and the lysis of 98.5 ± 2.4% cells in the culture, 46.7 ± 2.3% of the remaining cells were visibly infected at 30h p.i. (**Fig. 3e**).

### Irreversible adsorption’s (IA) role in determining stationary-phase infection dynamics

The essential step determining phage infection initiation is irreversible adsorption (IA) to the host cell. SPP1 irreversible binding to the bacterial receptor YueB [48] and the resulting inactivation of input phages is considered a proxy for the number of phages that engage in host infection [49]. The 3.4 ± 0.3-fold reduction in the titre of free infectious phages observed upon mixing of SPP1 with SP cells showed that SPP1 adsorbs relatively rapidly to nutrient deprived bacteria (**Fig. 3c**).

Availability of YueB at the bacterial surface is critical for SPP1 IA [48]. In order to investigate if this could be a limiting factor for SPP1 infection during SP we used immunofluorescence microscopy to quantify the proportion of cells exposing detectable amounts of receptor and, where present, cell surface YueB abundance at different stages of the *B. subtilis* growth cycle (**Supplementary figure 5a,b**). This was performed using *Bs*^*wt*^ and an isogenic strain overproducing YueB under the control of an IPTG-inducible promoter (*Bs*^*Pspac::yueB*^) (**Supplementary figure 5b, c**). The number of cell surface YueB foci decreased through TP (4-6h) into SP (10h) in both strains, though significantly more rapidly in *Bs*^*wt*^ (**Supplementary figure 5c**, analysis of covariance; χ^2 time_point:strain^ = 804, df = 3, p < 0.001). In parallel, the proportion of cells without detectable YueB foci increased sharply to 30.2% as *Bs*^*wt*^ entered into eTP, remaining high in lTP (53%) and SP (46.5%) (**Supplementary figure 5c**). This limitation was overcome in *Bs*^*Pspac::yueB*^ where surface exposed receptor was present on all cells (**Supplementary figure 5c**).

We then measured IA efficiency of SPP1^*wt*^ to both strains throughout the growth cycle. The two strains showed identical growth properties (**Supplementary figure 5a**) but drastically different profiles of phage SPP1 IA over the course of the growth cycle (**Fig. 4a**). During EP *Bs*^*wt*^ and *Bs*^*Pspac::yueB*^ had an IA of 99.2 ± 0.5 and 99.9 ± 0.007% of all added phages at 10min post-phage addition, respectively (**Fig. 4a**). From 4h post-inoculation onward (eTP through SP growth phases), *Bs*^*wt*^ bacteria supported IA of only ∼85-88% of all added phages while *Bs*^*Pspac::yueB*^ retained a ∼99.9% IA throughout its growth cycle (**Fig. 4a**).

**Fig. 4.**
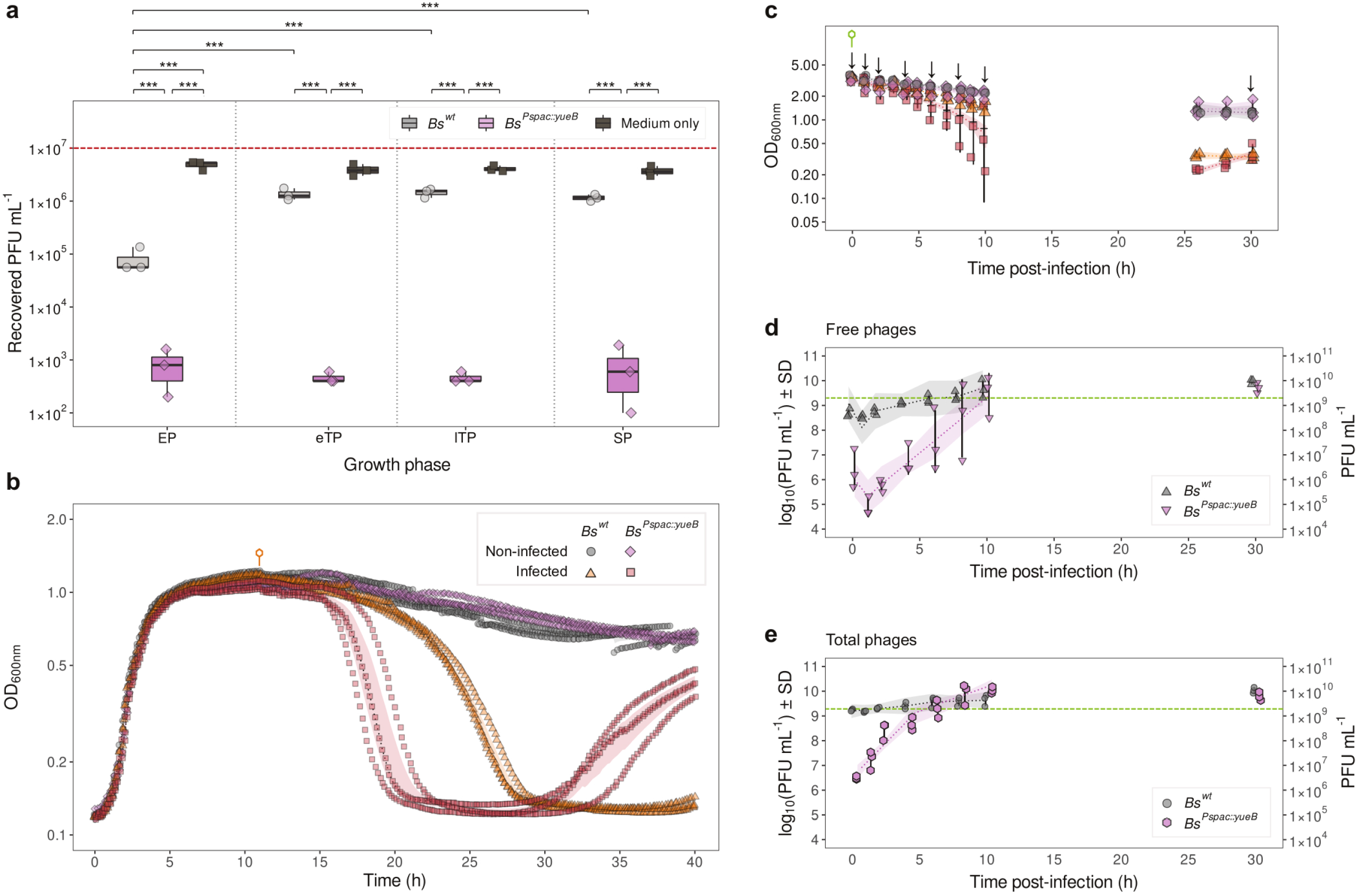
The impact of irreversible adsorption efficiency on stationary phase infection dynamics. **a** Irreversible adsorption (IA) of SPP1^*wt*^ at different phases of *Bs*^*wt*^ (grey circles) and *Bs*^*Pspac::yueB*^ (magenta diamonds) growth. IA was quantified as described [24] using bacterial growth medium (dark grey squares) as control for residual phage particle loss during manipulation. THC’s are displayed above the plot. Sampling points during bacterial growth are presented in **Supplementary figure 5a. b** Growth and lysis of bacteria infected with 2×10^9^ PFU ml^-1^ SPP1^*wt*^ (orange phage symbol) during SP in a 96-well plate monitored by OD_600_ measurements in a plate reader. Data points of non-infected or infected *Bs*^*wt*^ and *Bs*^*Pspac::yueB*^ are represented by the symbols shown in the inset. **c** Growth and lysis of bacteria infected during SP with 2×10^9^ PFU ml^-1^ SPP1^*mNeonGreen*^ (green phage symbol, for comparison with SPP1^*mNeonGreen*^ infections in **Fig. 3a**) in flasks was monitored by OD_600_ measurement. Symbols are as in **b**. Vertical arrows identify sampling time points for free and total PFU titration. **d,e** Titration of free and total infectious SPP1 particles, respectively, sampled at the time points shown in **c**. The dashed green line shows the input phages added for infection. Data in all panels are from three independent biological replicates. The median, upper and lower quartiles (boxes) and the limits of the data (whiskers) of the data are given in panel **a**. The mean (dotted lines) and standard deviation (shaded areas) are displayed in panel **b**. The mean (horizontal lines) and standard deviation (vertical bars) are displayed in panels **c**-**e**. Piece-wise linear regression models were fit to OD_600_ and PFU data for analysis (dotted lines (model fits) and shaded areas (confidence intervals) in **c**-**e**). Data and statistical analyses related to these experiments are provided in **Table S3** and **Supplementary Data Analysis**.

Significantly higher IA to *Bs*^*Pspac::yueB*^ than to *Bs*^*wt*^ prompted study of IA’s impact on the dynamics of SPP1 infection in SP. *Bs*^*Pspac::yueB*^ lysed more rapidly than *Bs*^*wt*^ during SP infection, albeit exhibiting some variability among experiments (**Fig. 4b,c**) that was reflected in the numbers of free PFU observed (**Fig. 4d**). In spite of this variability, the *Bs*^*Pspac::yueB*^ total phage yield was remarkably reproducible (**Fig. 4e**). The rate of phage production in Bs^*Pspac::yueB*^ cultures was faster than during infection of *Bs*^*wt*^ but, interestingly, the final phage yield at 30h p.i. was very similar (**Fig. 4d, e**).

### SPP1 gene expression during stationary phase infection of *B. subtilis*

The radically different dynamics and low yield of SP SPP1 infection prompted us to compare phage gene expression during SP infection with that of highly efficient EP infection. Using qRT-PCR we quantified mRNA expression of early genes *35* (recT-like recombinase [50]) and *46* (putative host take-over gene), of the late genes *6* and *11* (SPP1 procapsid assembly proteins), and of the late lysis genes *24*.*1* and *26* (**Fig. 5a**) [51,52]. qRT-PCR data were calibrated firstly to uninfected early EP samples to clearly assess patterns of early and late gene expression, and secondly to 8min p.i. EP samples for comparative gene expression analysis across different EP and SP infection time-points (**Fig. 5b**).

**Fig. 5.**
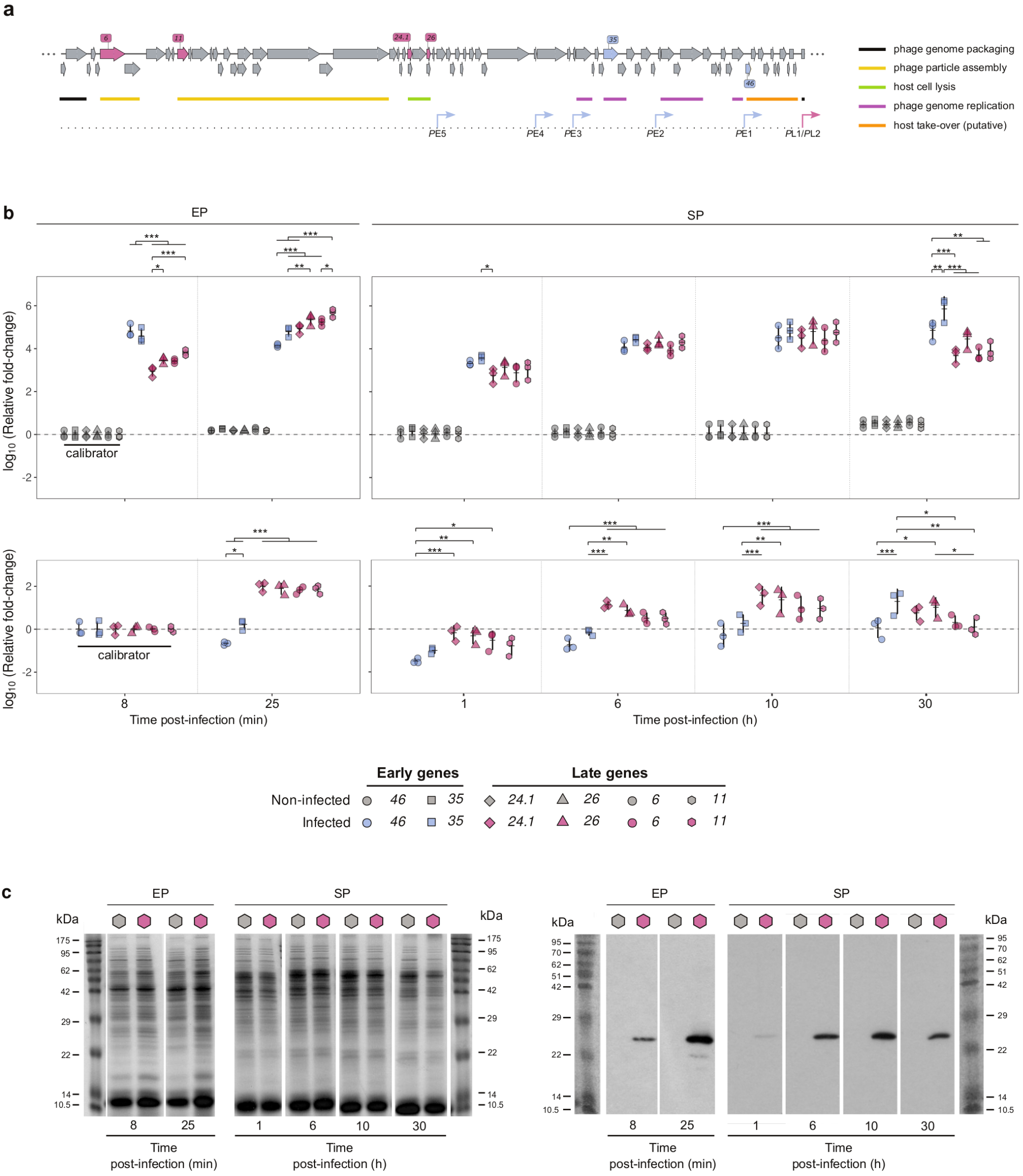
SPP1 gene expression during exponential and stationary phase infection. **a** Bacteriophage SPP1 genetic map. Early (light blue) and late genes (salmon) investigated in this study are highlighted. Functional classes of genes involved in different steps of SPP1 infection are identified by coloured bars annotated in the legend on the right. Early promoters (*P*E1 to *P*E5; blue) and the late promoters *P*L1/*P*L2 (salmon) are labeled below. Note that *P*L1/*P*L2 control the transcription of the late DNA packaging genes and gene *6* displayed on the left in the linear representation of the SPP1 genome. **b** Quantification of phage gene expression by qRT-PCR from *Bs*^*wt*^ during EP and SP phases infected with SPP1^*mNeonGreen*^ (**Supplementary figure 6a,d**). Samples of EP cultures were taken at 8 and 25min post-infection while SP cultures were sampled at 1, 6, 10 and 30h p.i.. Data were calibrated (‘calibrator’) to the gene expression of non-infected EP bacteria (*top*) and to that of infected EP bacteria at 8 min p.i. (*bottom*). The expression of early genes *35* and *46* (blue symbols), and of late genes *6, 11, 24*.*1* and *26* (magenta symbols) was quantified as described in detail in **Supplementary Materials and Methods**. THC’s are displayed above the plots. Statistical significance levels: * *p* < 0.05, ** *p* < 0.01, *** *p* < 0.001. **c** Production of procapsid scaffolding protein gp11 in EP and SP-infected bacteria. Samples were taken from the same cultures and time-points as RNA used for qRT-PCR in **b**. Total protein was stained with Coomassie blue in SDS-PAGE gels (left) and gp11 was detected specifically in western blots with anti-gp11 antibodies (right). Gel loading was normalised using an identical amount of total protein. Data in **b** are from three independent biological replicates. Data in **c** are representative of three independent Coomassie-stained gels and Western blots. The mean (horizontal lines) and standard deviation (vertical bars) are displayed in **b**. Data and analyses related to these experiments are detailed in **Table S3** and **Supplementary Data Analysis**. Colour version of figure available online.

During EP infection (**Supplementary figure 6a-c**) early gene expression was clearly dominant at 8min p.i. while stronger expression of late genes was evident at 25min p.i. (**Fig. 5b**, *left panels*). Between these two time points the early gene *46* was downregulated while early gene *35* was approximately equivalently transcribed (**Fig. 5b**).

During infection of SP cultures (**Supplementary figure 6d-f**) the transcriptional profile of SPP1 was very different (**Fig. 5b**, *right panels*). Phage gene expression increased more slowly than during EP infections as highlighted by calibration to infected early EP samples (**Fig. 5b**, *bottom*). The SPP1 expression temporal program in SP was far more homogeneous, displaying reduced late gene expression relative to early gene expression (**Fig. 5b**, *top*). At 1h p.i., expression of early genes was slightly higher than late genes while at 6 and 10h no differences in expression of the examined genes were observed (**Fig. 5b**, *top right*). Expression of early genes *35* and *46* in infected SP cells took 6 and 10 h, respectively, to reach approximately the same expression observed at 8 min p.i. of EP cells **(Fig. 5b**, *bottom*). Moreover, expression of late genes encoding SPP1 structural and lysis proteins at these time points p.i. was significantly lower than at 25min p.i. during EP **(Fig. 5b**, *top* and *bottom*; **Table S4**). Interestingly, the ∼24.5- and ∼11-fold lower expression of genes essential for viral particle assembly at 6 and 10 h p.i. in SP bacteria (**Table S4**), respectively, correlates particularly well with the reduction in yield of viable SPP1 particles relative to EP infection (**Supplementary figure 6c,f**). These numbers provide a good proxy of differences between infected cells at EP and SP as a large majority of cells are infected in both populations at the time points compared, as denoted by their lysis behavior (**Supplementary figure 6a,d,e**).

Intriguingly, the pattern of SP gene expression at 30h p.i. resembles that of early EP, though exhibiting higher expression of gene *35*. This occurs following extensive lysis of the infected culture (**Supplementary figure 6d-e**). Many (40.9 ± 14%) of the surviving cells at this time were detectably infected as revealed by fluorescence imaging of the SPP1^*mNeonGreen*^ reporter gene (**Supplementary figure 6g**). This may indicate new infections of a small fraction of previously uninfected cells.

Reduced transcription of late genes required for assembly of viral particles is a distinctive feature of SP infected cells and provides an explanation for the low yield of PFU. We thus monitored the production of gp11, a scaffolding protein present in hundreds of copies per SPP1 procapsid [53]. Strong expression of its encoding gene *11* in late EP (**Fig. 5b**, *left*) correlated with strong production of gp11 after 25min of EP infection (**Fig. 5c**). In contrast, after 1h of SP infection gp11 was present in very low quantities, and significantly less than at 8min EP p.i., demonstrating the slow pace of SP infection at the molecular level. SP gp11 levels initially increased, remaining roughly constant after 6h p.i., though in clearly lower quantities than during EP (**Fig. 5c**).

### The impact of nutrient influx on the outcome of stationary phase infection

Our results show that the duration and productivity of the infectious cycle of SPP1 are radically altered upon infection of nutrient-limited bacteria, notwithstanding the large number of bacteria infected. We posited that this persistent infection state of the host population would enable SPP1 to multiply rapidly when new nutrients become available, stimulating bacterial metabolism. To test this hypothesis we grew *Bs*^*wt*^ in a plate reader and infected cells at SP before supplementation with maltose (15mM) at 0, 1, 4 and 8h p.i. (**Fig. 6a**). Maltose is an energy-rich carbon source that causes only very weak carbon catabolite repression in *B. subtilis* [54].

**Fig. 6.**
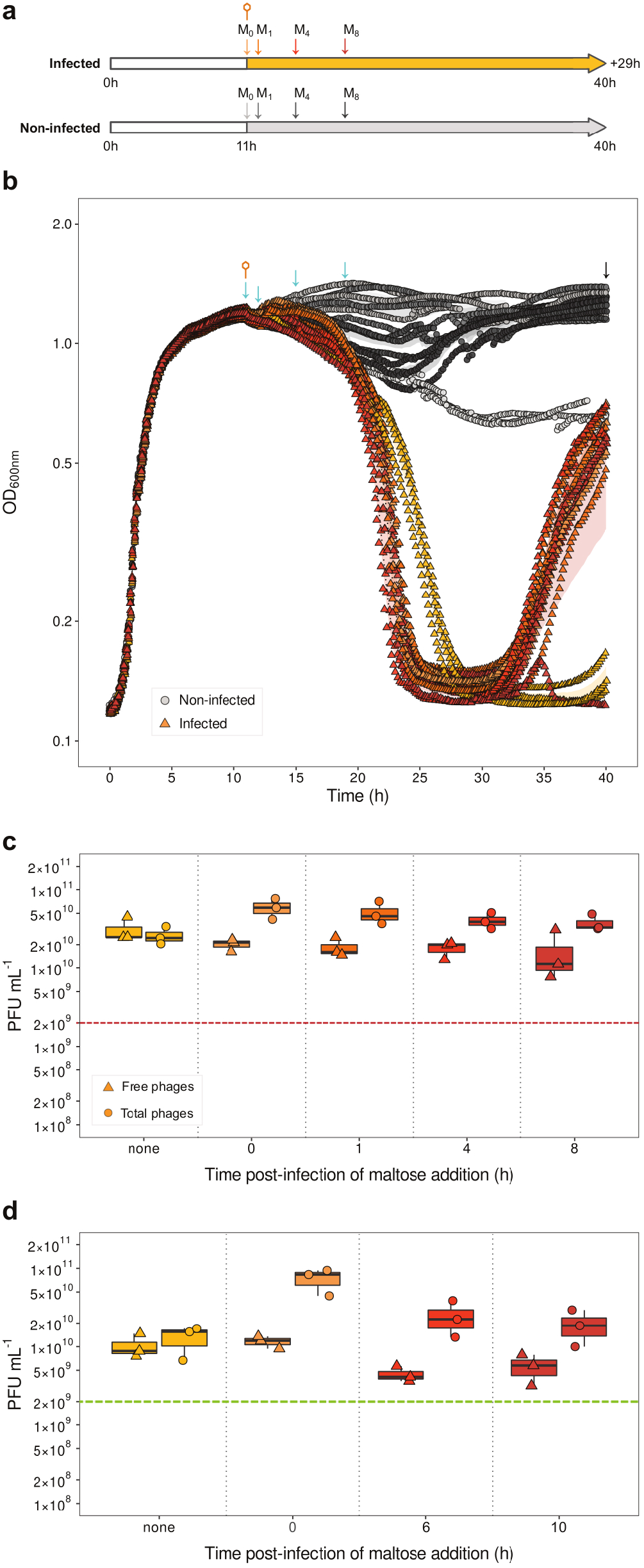
Influence of nutrient supplementation on stationary phase infection of *B. subtilis* by SPP1. **a** Experimental setup used to assess the effect of adding the carbon source maltose to SP bacteria non-infected (grey) and infected with SPP1 (orange). Phage infection is denoted by the orange phage symbol and times of maltose addition by the letter ‘M’. **b** OD_600nm_ curves of *Bs*^*wt*^ cultures grown in a 96-well plate and monitored continuously in a plate reader. Bacteria were infected with 2×10^9^ pfu ml^-1^ of SPP1^*wt*^ at 11h post-inoculation (SP) (orange triangles) and supplemented, in some cases, with 15mM maltose (triangles are coloured in variations of orange as the vertical arrows in **a**, dependent on the time of addition of maltose). Growth curves of non-infected bacteria supplemented with maltose (variations of dark grey according to the vertical arrows greyscale in **a**) or not supplemented (light grey) are also displayed. The orange phage symbol denotes the time of infection, blue arrows denote times of maltose addition and the black arrow the time of titration of PFU. **c** Titre of total (circles) and free phages (triangles) at 29h p.i. from the experiments in **b**. Color code is as in **a** and **b**. A dashed line displays the phage input for infection. **d** Titre of total and free phages at 30h p.i. of *Bs*^*wt*^ grown and infected at SP in flasks as in **Fig. 3**. The infection and maltose addition time-points were adapted to flask infection conditions. Data display is as in **c**. Data are from three independent biological replicates. The mean (dotted lines) and standard deviation (shaded areas) are displayed in panel **b**. The median, upper and lower quartiles (boxes) and the limits of the data (whiskers) of the data are given in panels **c,d**. Data and analyses related to these experiments are detailed in **Table S3** and **Supplementary Data Analysis**. Colour version of figure available online.

Maltose addition abrogated the reduction of OD_600_ observed during late SP (dark grey curves in **Fig. 6b**). Supplementation with maltose of infected bacteria in SP led to extensive, synchronised, lysis of infected cultures reaching 50% of the maximal OD_600_ at ∼10.5h p.i, irrespective of the time of its addition (**Fig. 6b**). Lysis occurred later and less abruptly in absence of exogenous maltose (50 % of the maximal OD_600_ after ∼12h p.i.).

Approximately 6.5 h after lysis, cultures supplemented with maltose resumed exponential growth, likely nourished by maltose and the pool of nutrients released from previously lysed cells (**Fig. 6b**). These cells were infected by SPP1 as demonstrated by the significantly higher number of total phages relative to free phages in the culture at the end of the experiment (**Fig. 6c**). This effect was amplified in experiments performed in flasks (**Fig. 6d**). In contrast, infected cultures that were not supplemented with maltose lysed later, regrew only very poorly within the experimental time-frame (**Fig. 6b**), and most phage particles were free in the culture at the end of the experiment (**Fig. 6c**).

## Discussion

We report the discovery that lytic bacteriophage SPP1 stages persistent, productive infections in planktonic cultures of nutrient-limited *B subtilis*. In this persistent infection mode, the infection cycle spans >10h. Lysis initiates slowly, followed by a rapid mass lysis phase. It is unlikely that lysis of bacteria from without [55] occurs because SPP1 particles do not kill *B. subtilis* upon contact and non-infected cells are insensitive to the SPP1 lysin gp25 released from lysed bacteria in liquid culture [56]. A vast majority of cells in SP are therefore infected, with lysis resulting from complete infection cycles. The pace and overall productivity of this persistent SP infection mode contrasts starkly with that of rapid, highly productive EP infections (**Figs. 1,2**). We conclude that SPP1 is equipped for fast and efficient multiplication in metabolically active host cells, but also for engaging a persistent infection mode for its environmental maintenance under adverse conditions. This adaptive strategy offers an explanation for how lytic phages may maintain their abundance in natural ecosystems such as the soil where bacteria, like *B subtilis*, exist primarily in nutrient-limited stationary phase [1,2]. Under such conditions, a pool of virions would be renewed by persistent infection while infrequent blooms of exponentially growing host bacteria would be efficiently exploited for major phage amplification.

Discovery of this persistent infection mode prompted investigation of how landmark steps of the viral infectious cycle were affected. Phage adsorption to host cells, the first step of infection, is impacted by scarcity of the SPP1 receptor YueB at the bacterial cell surface [24,25]. The effect of reduced SP cell surface YueB abundance was assessed experimentally in a strain overproducing YueB that maintains very high levels of SPP1 irreversible adsorption through the complete bacterial growth cycle (**Fig 4a**). More rapid extensive lysis of this strain (**Fig. 4b,c**) showed that limited YueB availability in wild-type SP *B. subtilis* extends the overall infection period in the bacterial population. Overall scarcity and cell-to-cell variation of YueB abundance (**Supplementary figure 5b,c**) possibly leads to some asynchrony of infection initiation. However, infected wild type bacteria still undergo a rather rapid extensive lysis phase (**Fig. 4b,c**) indicating that the long infection cycle duration minimizes this asynchrony effect. SPP1 multiplied faster in the YueB-overproducing strain but, strikingly, the final yield of infectious particles was identical to the wild type strain (**Fig. 4d,e**). Collectively, these results show that SPP1 adsorption to SP *B subtilis* impacts infection dynamics but does not determine the overall yield of virions produced during infection of a given SP bacterial population. Phage yield is therefore mostly determined by the limited metabolic capacity of nutrient-limited cells.

The program of SPP1 gene expression differs significantly between EP and SP infection both in temporal program and overall gene expression (**Fig. 5b**). Unlike EP infection, during persistent SP infection the expression of early and late genes remained roughly similar until extensive lysis. Importantly, late in infection, an ∼11-fold reduction of transcription of genes encoding proteins essential for virions assembly was observed relative to late EP infection (**Fig. 5b**; **Table S4**). This reduction correlates particularly well with the decrease in number of virions produced during persistent SP infection relative to rapid EP infection (**Supplementary figure 6c,f**). Reduction of expression of genes encoding phage structural proteins is particularly meaningful in the context of a nutrient-deprived host cell because production of virion components represents the highest biosynthetic and energetic cost during infection [38,57]. Therefore, SPP1 pervasively infects SP bacterial populations for a long period, effecting multiplication at reduced costs to the cell to achieve successful infection under adverse conditions.

Expression of SPP1 lysis genes *24*.*1* and *26* [58] is also lower than during late EP infection (**Fig. 5b**; **Table S4**). Such reduction does not hinder the capacity of SPP1 to promote SP cell lysis, and efficient release of virions into the environment is ensured. When a carbon/energy source (maltose) is made available to the infected SP culture lysis is more rapid. However, the time of maltose addition did not affect lysis behaviour (**Fig. 6b**). We conclude that the molecular clock for phage-induced host cell lysis is set upon infection initiation and that the added exogenous carbon source only renders lysis more effective. The exact mechanism by which this occurs is currently unknown. Interestingly, infected cultures supplemented with maltose re-grow after extensive phage-induced lysis and are clearly re-infected (**Fig. 6b-d**). Such re-growth, fueled by maltose and likely by nutrients released from lysed bacteria, offers SPP1 the opportunity for a new infection cycle mimicking conditions of host encounter of new nutrients. Indeed, un-supplemented cultures do not re-grow significantly but the low population of cells surviving mass culture lysis are extensively infected (**Fig. 3d,e**) and feature a pattern of SPP1 gene expression reminiscent of early EP infection (**Fig. 5b**).

Infection in stationary phase was previously reported for several phages [17,59–63] but their modes and dynamics of infection remain largely uncharacterised [62]. Phage T7 has the remarkable capacity for productive infection of stationary phase bacteria [59]. Other lytic phages adopt a dormant state in such nutrient-limited bacteria. In pseudo-lysogeny lytic phage DNA resides in the cell in an inactive state [18,63] while in the hibernation mode, described for phage T4, some early genes are expressed but the infection cycle is halted until further nutrients become available to the host cell [17]. In contrast, the above presented SPP1 persistent infection mode involves a complete productive infection process. Here, we investigated this infection mode in a simplified single phage-host pair laboratory based setup in order to carry out a detailed characterisation of their interaction. The soil environment imposes much more complex, multi-factorial, constraints to the SPP1-*B subtilis* interaction [62,64]. Nevertheless, SPP1’s persistent infection mode appears to represent an adaptive strategy that would aid persistence of lytic phage populations under frequently encountered environmental conditions that heavily restrict host bacterial growth. Infected nutrient-limited cells are killed and phage particles released. Environmental decay of free phage particles is a threat for these free particles [65]. However, nutrients liberated by lysis can also fuel new infections of surviving susceptible host cells. This will sustain local phage populations in the medium term while awaiting nutrient influx, while additional phage particles may be disseminated to find new susceptible hosts.

## Author contributions

JD and PT designed the research. JD performed most experimental work with help from PT. NN and EJ performed qRT-PCR, including preceding validation and quality assessment work. ACL designed and constructed SPP1^mNeonGreen^. Data were analysed by NN, EJ and JD. Data were interpreted by JD and PT. The manuscript was written by JD, EJ and PT. Work was read and critically revised by JD and PT. All authors approved the final version of the manuscript.

## Acknowledgments

We thank Sriram Tiruvadi-Krishnan for providing the plasmid bearing the *mNeonGreen* gene codon-optimised for *B. subtilis*. This work was funded by Fondation pour la Recherche Médicale (Equipe FRM), grants ANR BacVirRemodel and ANR BioBrickEvolver, and institutional funding from CNRS.

