## Supplementary_information for "Persistent lytic bacteriophage infection as a novel strategy for exploitation of nutrient-limited host bacteria"


##### This PDF file includes:


- **Supplementary Materials and Methods**
- **Supplementary figure 1** (*Correspondence between manual counts of cells infected with SPP1$^{mNeonGreen}$ and counts derived from semi-automatic identification*)
- **Supplementary figure 2** (B. subtilis *growth and morphology during exponential, transition and stationary phase*)
- **Supplementary figure 3** (*Infection of* B. subtilis *by SPP1 in semi-solid medium*)
- **Supplementary figure 4** (*Visualisation of mNeonGreen reporter during exponential phase infection*)
- **Supplementary figure 5** (*YueB presentation at the cell surface during growth of wild-type and YueB-overproducing* B. subtilis)
- **Supplementary figure 6** (*Characterization of phage infection cultures used for SPP1 gene expression analyses*).
- **Table S1** (*Bacterial strains, phage strains and plasmids used in this study*)
- **Table S2** (*Primers used in this study*)
- **Legend for Table S3**
- **Table S4** (*SPP1 gene expression during stationary phase (SP) infection compared to exponential phase (EP) late stage of infection*)
- **Supplementary Data Analysis** (*This data analysis summary includes statistics, statistical models used, their outputs and any multiple comparison testing made for each experiment. Data are organised by Figure and their respective panels. All model / test outputs are as given by R, minimally edited to make interpretation easier for the reader*)


##### Other supplementary materials for this manuscript include the following:

- **Table S3** (.xlsx file) (*Supporting information on bacterial growth, phage induced lysis, and PFU production for main Figs. 1 to 4 and 6 and to Supplementary figures 2, 5, and 6*)

### Supplementary Materials and Methods

#### Bacterial strains and growth

Bacterial strains are described in **Table S1.** *B. subtilis* 168 strain 1A1 [1], designated $Bs^{wt}$, was used as the wild-type strain for infection experiments. The prophage-free strain YB886 [2], was used for phage amplification and for phage titration unless stated otherwise.

*B. subtilis* strain $Bs^{Pspac::yueB}$ was constructed by SPP1-mediated transduction of the $P_{spac}$*::yueB* cassette from the donor strain YB886.PspacYB [3] into $Bs^{wt}$. Correct insertion of the cassette into the native *yueB* locus was verified by genomic DNA (gDNA) extraction followed by PCR and Sanger sequencing (Eurofins; Ebersberg, Germany) using primers listed in **Table S2**.

Pre-cultures were normally grown overnight in LB medium (Conda-Pronadisa; Madrid, Spain) at 30°C for 16-18h with agitation at 150rpm. Pre-cultures for stationary phase infection experiments were grown at 37°C for 12h. Pre-cultures were diluted to an optical density at 600nm ($OD_{600}$) of 0.05 and grown at 37°C. Strain $Bs^{Pspac::yueB}$ was grown in presence of 1mM IPTG for overproduction of YueB [3,4]. Bacterial growth was followed by monitoring $OD_{600}$. For some experiments, bacteria were cultured in 96-well flat-bottomed plates in a TECAN 200 infinite Pro plate reader to follow long-term bacterial growth or infection dynamics. In these experiments bacteria were cultured at 37°C with continuous shaking, taking $OD_{600}$ readings every 5min.

#### Phage strains

Phage strains are described in **Table S1.** Phages were amplified in *B. subtilis* YB886, purified by isopycnic CsCl density gradient centrifugation [3] and stocked in TBT buffer (100mM Tris-HCl, 100mM NaCl, 10mM $MgCl_2$ [pH 7.5]) [5].

SPP1$^{mNeonGreen}$ was constructed by insertion of an RBS-*mNeonGreen*$^{opt}$-STOP cassette, bearing a *mNeonGreen* gene codon-optimised for *B. subtilis* [6], into the SPP1 early gene region under the control of *P*E2. A PCR fragment bearing SphI restriction sites flanking the cassette was digested with SphI and ligated with SPP1$^{wt}$ DNA cleaved at its unique SphI restriction site, disrupting SPP1 gene *43*. The ligation reaction was transfected into YB886 competent cells and pure phage clones were isolated as described in [3]. Correct insertion was confirmed by PCR and Sanger sequencing. Primers are listed in **Table S2**.

**Quantification of bacterial colony forming units (CFU)**

Bacterial culture samples were first diluted 1:10 in Bott and Wilson (BW) salts (71.2mM $K_2HPO_4$, 58.8mM $KH_2PO_4$, 3.87mM trisodium citrate, 54.4mM di-ammonium sulphate). This sample was serially diluted in the same buffer and appropriate dilutions plated on LB 2% agar plates before incubation overnight at 37°C. To determine spore CFU counts the initial 1:10 dilution was heated to 80°C for 20 minutes, allowed to cool, and then processed as above.

**Phage infection, lysis curves, quantification of plaque forming units (PFU) and irreversible adsorption assays**

Phage infections were initiated with $2\times10^9$ PFU ml$^{-1}$ as standard, ensuring a multiplicity of infection (MOI) of ~3-5 in high-density cultures while maintaining a comparable PFU input baseline between infections. Cultures were supplemented with 10mM $CaCl_2$ prior to infection. Bacterial lysis was followed by measurement of cultures $OD_{600}$. Extracellular infectious phages were separated from cells by centrifugation (16,000*g*, 3min, room temperature (RT)) and isolation of the supernatant. For total (extra and intracellular) infectious phage counts, a 1:10 dilution of infected bacterial culture in TBT was mixed vigorously with chloroform for 30s, settled on ice for >2h and the aqueous phase isolated. These samples were diluted and phages enumerated by titration [7]. SPP1 irreversible adsorption (IA) was measure as described in [8].

**Cell labelling, wide-field fluorescence microscopy and image processing**

Immunofluorescent labelling of YueB at the surface of live *B. subtilis* cells was performed as described in [3], using a polyclonal rabbit YueB antibody followed by detection with a goat anti-rabbit IgG AlexaFluor 488 fluorescent dye conjugated secondary antibody (Molecular Probes). For membrane staining 25μM TMA-DPH (Molecular Probes) was added to culture samples and incubated 3-5min at 37°C with 150rpm shaking. For imaging of bacteria, 3μl of labelled culture was placed onto a thin agarose pad (0.1% w/v agarose in water) on a glass microscope slide and covered with a coverslip.

Microscopy was performed using an Axio Observer.Z1 (Zeiss; Marly Le Roi, France) microscope equipped with a 63x oil immersion lens (Zeiss), coupled to a Orca-Flash4.0 digital CMOS camera (Hamamatsu; Hamamatsu City, Japan). DAPI (Ex. 381-392, Em. 417-477) and GFP (Ex. 457-487, Em. 502-537) filter sets were used to image TMA-DPH and mNeonGreen/AlexaFluor 488 fluorophores, respectively. An exposure time of 200ms was used for all fluorescence acquisitions. Post-capture image processing was carried out using Fiji [9]. A linear adjustment of contrast was applied across entire images to enhance clarity, while ensuring no clipping of high or low intensity signal. No further processing of images was performed.

**Extraction of data from microscopy images and semi-automatic identification of infected cells**

MicrobeJ [10] was used to extract morphological and fluorescence data from individual cells in wide-field microscopy images. Cut-offs for cell area, circularity and intensity were used to exclude cell debris and lysed cell 'ghosts' during cell detection steps using the phase contrast channel. Any out-of-focus or incompletely visible cells were removed by manual curation. Short chains or paired cells were manually segmented into individual cells using membrane staining signal to identify complete division septa. All cells exhibited some autofluorescence at a similar wavelength to mNeonGreen fluorescence. Extracellular background fluorescence measurements (4 per image, each $100px^2$) were used to correct original fluorescence measurements by subtraction of the mean background measurement. Following inspection and analysis of the distribution of fluorescence data in non-infected and infected samples, plus validation through comparison with manual counts of infected cells (**Supplementary figure 1**), a cut-off equal to 4- standard deviations above the mean auto-fluorescence of non-infected cells was determined as appropriate to delimit infected from non-infected cells. Assigning infection status based on mNeonGreen fluorescence intensity of each measured cell in `R`, then allowed rapid determination of proportions of infected cells among very large total numbers of cells.

**RNA extraction and qRT-PCR**

Cultures were infected in exponential phase (EP) or SP with $2\times10^9$ PFU $ml^{-1}$ SPP1$^{mNeonGreen}$ as described above. At EP time-points 8 and 25min post-infection (p.i.) and SP time-points 1, 6, 10, and 30h p.i., culture samples were mixed 9:1 with 10x ice-cold sample stop buffer (100mM Tris-HCl [pH 7.5], 100mM NaCl, 10mM $MgCl_2$, 100mM $NaN_3$), centrifuged at 8000*g* for 5min, their supernatant removed, centrifuged again for 1min, remaining supernatant removed, and finally snap frozen by immersion in liquid nitrogen. Sample volumes were normalised across all time-points by $OD_{600}$ for isolation of RNA except SP 30h p.i. where 30ml samples were taken as sufficient RNA proved difficult to isolate from smaller volumes. RNA was isolated from samples using a 'NucleoSpin RNA Mini' RNA purification kit (Macherey-Nagel) according to the manufacturer's instructions, with some slight modifications. All samples were purified twice, with a digestion of 15min at 37°C using DNase I (NEB) in between, to effectively remove contaminating gDNA from the samples. Large volume samples (30ml) were re-suspended and split into aliquots equivalent to 2ml culture each for isolation, as initial tests revealed that this was required due to the high ratio of cellular debris to intact cells in these samples. Individual purifications were pooled after the first round of purification. All RNA was stored at -80°C before further treatment.

RNA concentration was quantified by Nanodrop One spectrophotometer (ThermoFisher Scientific) and RNA integrity verified using the Agilent 2100 bioanalyzer with an RNA 6000 Nano kit (Agilent Technologies). A High Capacity cDNA Reverse Transcription Kit (Applied Biosystems) was used to reverse-transcribe 200ng of total RNA using random primers in a final reaction volume of 20µL, in the presence of RNase inhibitor.

Quantitative real-time PCR (qRT-PCR) was performed on a QuantStudio 12K Flex Real-Time PCR System (Life Technologies) with a SYBR green detection protocol. 1.5ng of cDNA was mixed with Fast SYBR Green Master Mix and either 500 or 750nM of each primer in a final volume of 10µL. The reaction mixture was loaded into 384-well microplates and submitted to 40 cycles of PCR (95°C, 20s; [95°C, 1s; 60°C, 20s] × 40) followed by a fusion cycle to analyze melting curves of PCR products. qPCR in the absence of a reverse transcription step was performed on all RNA samples to check the absence of DNA contamination. Primers were designed using the Primer-Blast tool from NCBI and the Primer Express 3.0 software (Life Technologies). Specificity and the absence of multilocus matching at the primer site was verified by BLAST analysis. The amplification efficiencies of primers were determined using the slopes of standard curves obtained over a five-fold dilution series. Amplification specificity for each real-time PCR reaction was confirmed by analysis of dissociation curves. Technical duplicate measurements were made for each independent biological replicate. Determined Ct values were then used for further analysis.

For data normalisation nine *B. subtilis* reference genes were tested (*gyrA*, *rplP*, *rpsE*, *era*, *secA*, *gmk*, *rpoB*, *dnaG*, *ftsZ*). The most stable reference genes throughout the cell cycle of *B subtilis* (*gyrA*, *secA*, *rpoB*, *dnaG*, *ftsZ*) were selected by GenEx software (MultiD) and the geometric mean of the five most stable genes was used to normalise the data. Relative gene expression ratios were determined using the ΔΔCt method. Calibration was performed relative to control EP non-infected samples (**Fig. 5b** *top*) grown in parallel to infected cells sampled at 8min (**Supplementary figure 6a**), to 8min EP post-infection infected samples (**Fig. 5b** *bottom*) or to 25min EP post-infection infected samples (**Table S4**). All values, including that of the calibrator, were divided by those of the calibrator to give fold-change values relative to the calibrator. The limit of detection for all genes was set at a Ct limit of 33 cycles. This in turn determined the 'phage gene expression' in non-infected samples used for calibration to 8min (**Fig. 5b** *top*).

**Protein extraction, polyacrylamide gel electrophoresis and Western blotting**

Protein samples were prepared from culture samples normalised across time-points by $OD_{600}$ as described for RNA extraction. Protein was extracted as described previously [11] and run immediately on two parallel 15% polyacrylamide gels. One gel was stained with

Coomassie blue and the other used for Western blotting. Given these initial Coomassie staining results, sample concentrations were adjusted and gels repeated with approximately equal amounts of protein in each sample. For Western-blotting, gp11 was detected using a rabbit polyclonal α-gp11 antibody [12], followed by a goat anti-rabbit IgG secondary HRP-conjugated antibody (Sigma) and revealed using ECL Western Blotting Detection Reagents (Amersham).

**Quantitative and statistical data analysis**

Quantitative and statistical analyses were performed using `R` version 3.6.3 [13].

Growth and lysis rates were derived by fitting cubic smoothing splines to time-series data (either $OD_{600}$ or PFU $ml^{-1}$) using the `smooth.spline` package [13] and predicting this fit at the same x-values (time). The first derivative (gradient or rate) of the predicted fit at each x-value was calculated using the `predict` package [13]. Maximum or minimum rates were extracted where appropriate. For extraction of lysis rates from multiphasic lysis curves, the second derivative of the curve was first calculated and used to identify inflection points, accompanied by manual inspection. Mid-points between these inflection points were then used to delimit different zones of the curves (see **Supplementary Data Analysis** for determined break-points). Maximum decline (or lysis) rates were then taken from each respective curve region. Where possible to determine, the time to lysis initiation was calculated as the time between infection and the switch from a positive to negative gradient of the curve. Proportion loss of $OD_{600}$ was calculated as a ratio between the maximum and minimum $OD_{600}$ values reached during the analysis period. The time to 50% loss of maximum $OD_{600}$ was calculated from the time of infection until the first time-point at which $OD_{600}$ was less than 50% of the maximum recorded in the respective culture. Duration of infection was calculated from the time of infection until the first time-point at which the rate of decrease in $OD_{600}$ fell below the average rate of decline of the non-infected culture during stationary phase of each respective experiment.

Fold-change in PFU was calculated as a ratio between an initial input of $2\times10^9$ PFU $ml^{-1}$ and the final phage titre within the assay period. Where possible, average burst size was calculated as the number of PFU produced per each CFU lost during a given time-period.

All statistical analyses involving more than 2 independent groups were performed using linear modelling. The package `lm` was used to fit general linear models (LM's) to normally distributed, constant variance (or transformed) data, and `glm` to fit generalised linear models (GLM's) to untransformed data requiring different error structures or link functions. GLM error structures and link functions were chosen based on the interpretation of the fit of LM's to untransformed data. Maximal models containing all possible variable interactions were constructed, followed by iterative model simplification by likelihood ratio testing to produce minimal adequate models [14] for statistical analysis (**Supplementary Data Analysis**). For piece-wise regression analyses, breakpoints were determined using the `segmented R` package, supplying basic linear models (see **Supplementary Data Analysis**) and manually estimated breakpoints as input.

Results are given from computed analysis of variance (F; LM) or deviance ($\chi^2$; GLM) tables. Multiple comparisons were made using either `multcomp` [15] or `emmeans` [16] packages, as required, to calculate Tukey's honest significant difference contrasts (HSD). Where data were transformed before analysis or when generalised linear models using link functions producing outputs not on the response scale, contrasts were appropriately back-transformed for interpretation. Model predictions for visual representation were generated using the `predict` package [13]. For LM's predicted confidence intervals (CI's) were generated automatically, and for GLM's by multiplying the fitted value on the link scale ± 2·SE by the inverse link function of the model. In some cases two-tailed t-tests or Wilcoxon rank sum tests were run using appropriate `R` packages [13]. In one case (**Fig. 1e**) data were log-transformed to conform to t-test assumptions. Correlations were assessed through calculation of Pearson correlation coefficients using the `R cor` package [13].

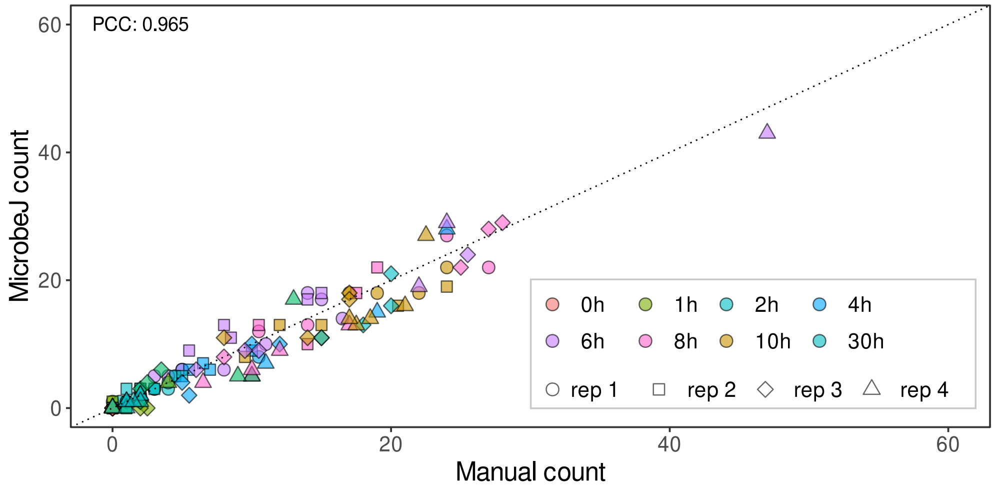


**Supplementary figure 1 Correspondence between manual counts of cells infected with SPP1$^{mNeonGreen}$ and counts derived from semi-automatic identification.** Infected cells were identified by the presence of mNeonGreen fluorescence intensities >4·SD above the mean corrected fluorescence intensity of non-infected cells (see **Materials and Methods**). Visibly infected cells were also counted manually to allow an assessment of the accuracy of the identification of infected and non-infected cells using our semi-automatic method. Each data point is derived from a single fluorescence image. This analysis was performed using the entire dataset presented in **Fig. 3d,e** from four independent biological replicates (rep 1 to 4). The Pearson's correlation coefficient of the two counts is given in the panel top left-hand corner. The diagonal dotted line represents a perfect correlation of 1.

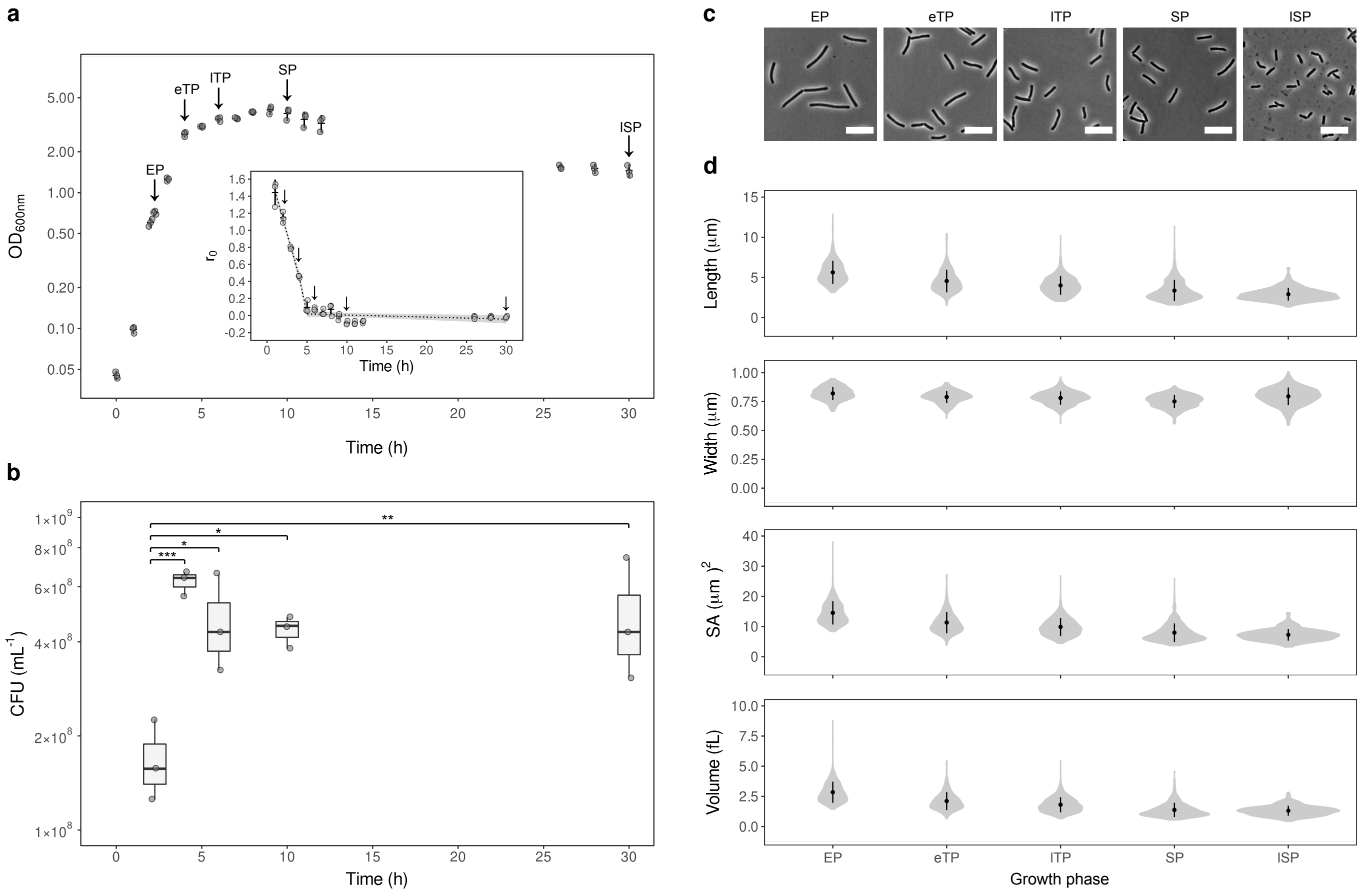


**Supplementary figure 2 *B. subtilis* growth and morphology during exponential, transition and stationary phase. a** $Bs^{wt}$ growth in flasks in LB medium at 37°C. Arrows identify the four post-inoculation time points selected for subsequent infection experiments at different stages of the bacterial growth cycle: exponential phase (EP), early transition phase (eTP), late transition phase (lTP) and stationary phase (SP). A fifth arrow identifies a late stationary phase (lSP) time-point in this experiment. Bacterial growth rates ($r_0$) at each time-point were calculated as described in **Supplementary Materials and Methods** and the time of growth arrest identified using a piece-wise linear regression model fit to $r_0$ data (dotted line, inset) as detailed in **Supplementary Data Analysis**. **b** CFU counts at the selected time points in **a**. Tukey's honest significant difference contrasts (THC's) are displayed in the upper part of the plot. Statistical significance levels: * $p < 0.05$, ** $p < 0.01$, *** $p < 0.001$. **c** Phase contrast images of $Bs^{wt}$ at different phases of culture growth. The white scale bars represent 10µm. **d** Morphological parameters of $Bs^{wt}$ bacterial populations at different phases of growth. The upper two subpanels show cell length and width measurements that were used to calculate cell surface area (SA) and volume using a spherical-ended cylinder as a proxy for bacterial shape. Shaded areas of the violin-plots display the probability density of the data at a given value. Data are from three independent biological replicates. The piecewise linear regression model fit to the data for statistical analysis in **a** (inset) is represented graphically by a dotted line (model fit) and shaded areas (95% confidence intervals). The mean (horizontal lines in **a**, solid circles in **d**) and standard deviation (vertical bars) are displayed in **a** and **d**. The median, upper and lower quartiles (boxes) and the limits of the data (whiskers) of the data are given in **b**. Sample sizes were as follows; EP; n = 305, eTP; n = 285, lTP; n = 369, SP; n = 395, lSP; n = 568. Additional data and analyses related to these experiments are presented in **Table S3** and **Supplementary Data Analysis**.

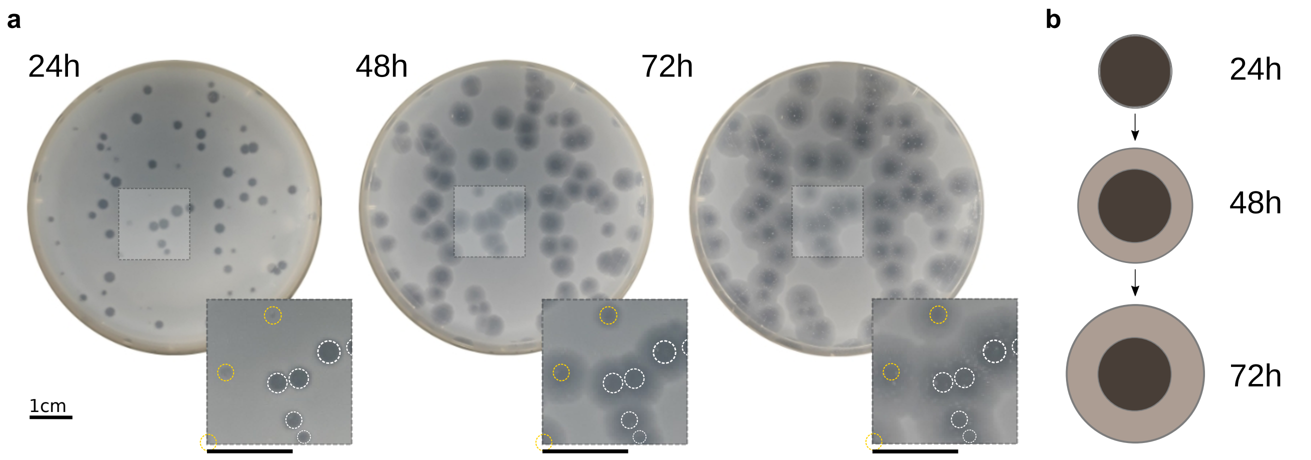


**Supplementary figure 3. Infection of *B. subtilis* by SPP1 in semi-solid medium.** $Bs^{wt}$ bacteria grown in LB medium at 37°C were sampled at mid-exponential phase ($OD_{600}$ ~0.8, 2.25h post-inoculation (**Supplementary figure 2a**)), mixed with 0.7% LB agar and infected with $SPP1^{wt}$ to yield a final count of ~100 PFU per plate. **a** Development of phage plaques over 72h. White dashed outlines in magnifications indicate plaques that had fully developed by 24h p.i. while yellow dashed outlines indicate plaques that had fully developed by 48h p.i. Diffuse halo areas around these outlines represent lysis either due to diffusion of lytic components in the medium or continued partial lysis due to infection. Scale bars (main and inset) represent 1cm. **b** Phage plaque development in schematic form.

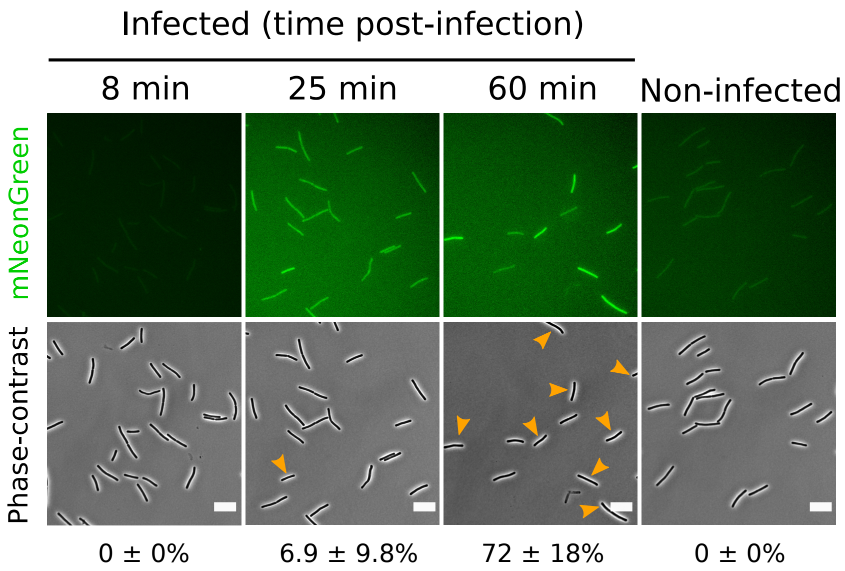


**Supplementary figure 4. Visualisation of mNeonGreen reporter during exponential phase infection.** *Bs*$^{wt}$ was infected with $2\times10^{9}$ PFU $ml^{-1}$ of SPP1$^{mNeonGreen}$ at mid-exponential phase and imaged at the indicated time-points. An MOI of 20 ensures infection of the complete bacterial population. Individual cells were counted at each time-point by microscopy and the percentage of infected cells in the culture determined computationally (see **Supplementary Materials and Methods**). A cut-off of +2·SD above the mean background-corrected fluorescence of non-infected cells was used to delimit infected from non-infected cells. This cut-off was chosen in place of +4·SD (see **Supplementary figure 1**) as background fluorescence at 25 and 60min post-exponential phase infection was high, probably due to release of mNeonGreen to the medium resulting from cell lysis. Orange arrowheads in the phase contrast images identify bright cells in the mNeonGreen channel. Data are representative of 3 independent biological replicates. The mean and standard deviation of the percentage of visibly infected cells is shown. Scale bars represent 10μm.

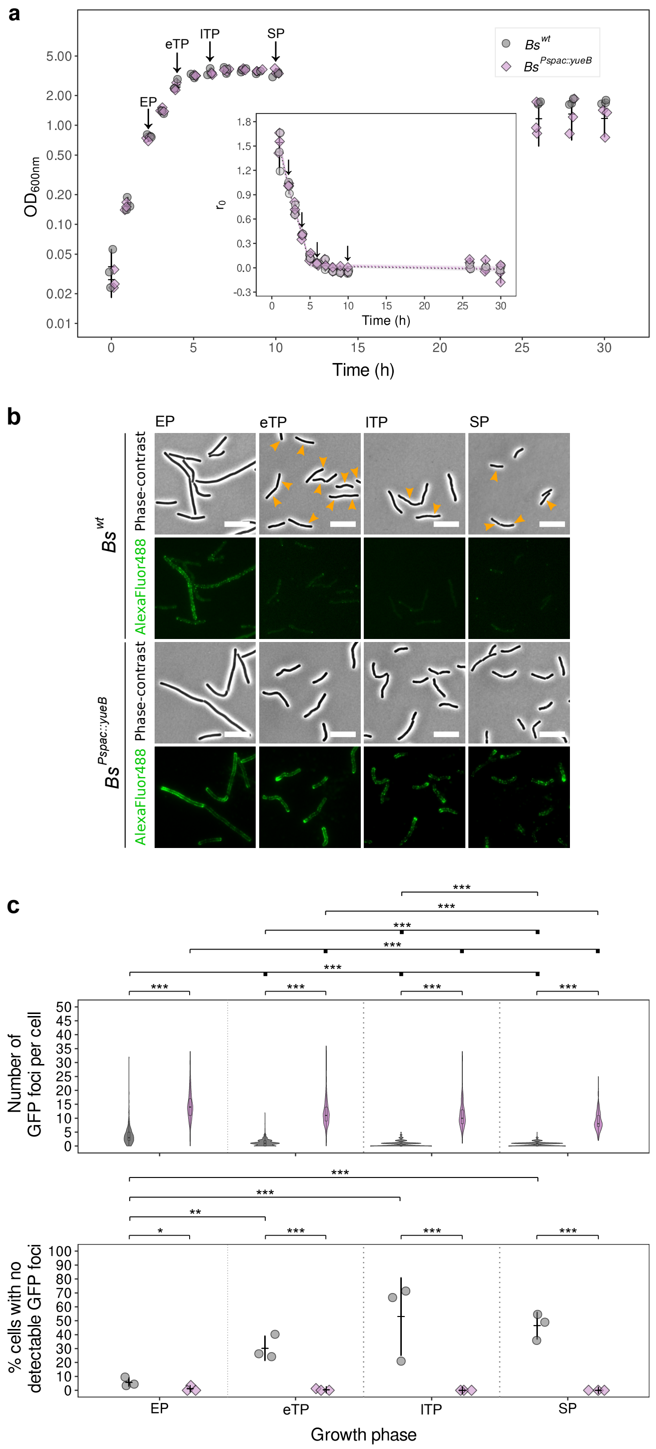


**Supplementary figure 5.**

**YueB presentation at the cell surface during growth of wild-type and YueB-overproducing *B. subtilis.*** **a** Growth curves of $Bs^{wt}$ (grey circles) and $Bs^{Pspac::yueB}$ (magenta diamonds) monitored until 30h post-inoculation. Growth rates ($r_0$; inset) declined significantly over time and significantly more slowly after 5.7h post-inoculation in both strains (see also **Supplementary Data Analysis**). Growth phase time-points defined for the two strains (vertical arrows) were identical. **b** Immunofluorescence microscopy of cell surface YueB in $Bs^{wt}$ and $Bs^{Pspac::yueB}$. Alexa Fluor 488-labeled YueB was imaged in the GFP channel. Orange arrowheads over the phase channel highlight the localisation bacteria presenting fluorescent foci (GFP channel) on the surface of $Bs^{wt}$ in transition and stationary phase. The white scale bars represent 10µm. **c** Quantification of the number of YueB foci per cell. Violins display the population distribution of the number of independent foci per cell of $Bs^{wt}$ (grey) and $Bs^{Pspac::yueB}$ (magenta) (*top*). Grey circles ($Bs^{wt}$) or magenta diamonds ($Bs^{Pspac::yueB}$) (*bottom*) show the percentage of cells possessing no detectable surface labeled YueB. Sample sizes were as follows; EP; $n^{wt}$ = 378, $n^{Pspac::yueB}$ = 467, eTP; $n^{wt}$ = 511, $n^{Pspac::yueB}$ = 527, ITP; $n^{wt}$ = 317, $n^{Pspac::yueB}$ = 335, SP; $n^{wt}$ = 416, $n^{Pspac::yueB}$ = 424. THC's are displayed above each plot. Statistical significance levels; * $p < 0.05$, ** $p < 0.01$, *** $p < 0.001$. Where multiple samples are compared, comparisons are made from the leftmost sample (thin vertical tick) to each other sample (thick vertical ticks). Data are from three independent biological replicates. The linear regression model fit to the data for statistical analysis in **a** (inset) is represented graphically by dotted lines (model fit) and shaded areas (95% confidence intervals). The mean (horizontal lines) and standard deviation (vertical bars) are displayed in **a** and **c** (*bottom*). The median, upper and lower quartiles (boxes) and the limits of the data (whiskers) of the data are given in **c** (*top*). Data and analyses related to these experiments are presented in **Table S3** and **Supplementary Data Analysis**.

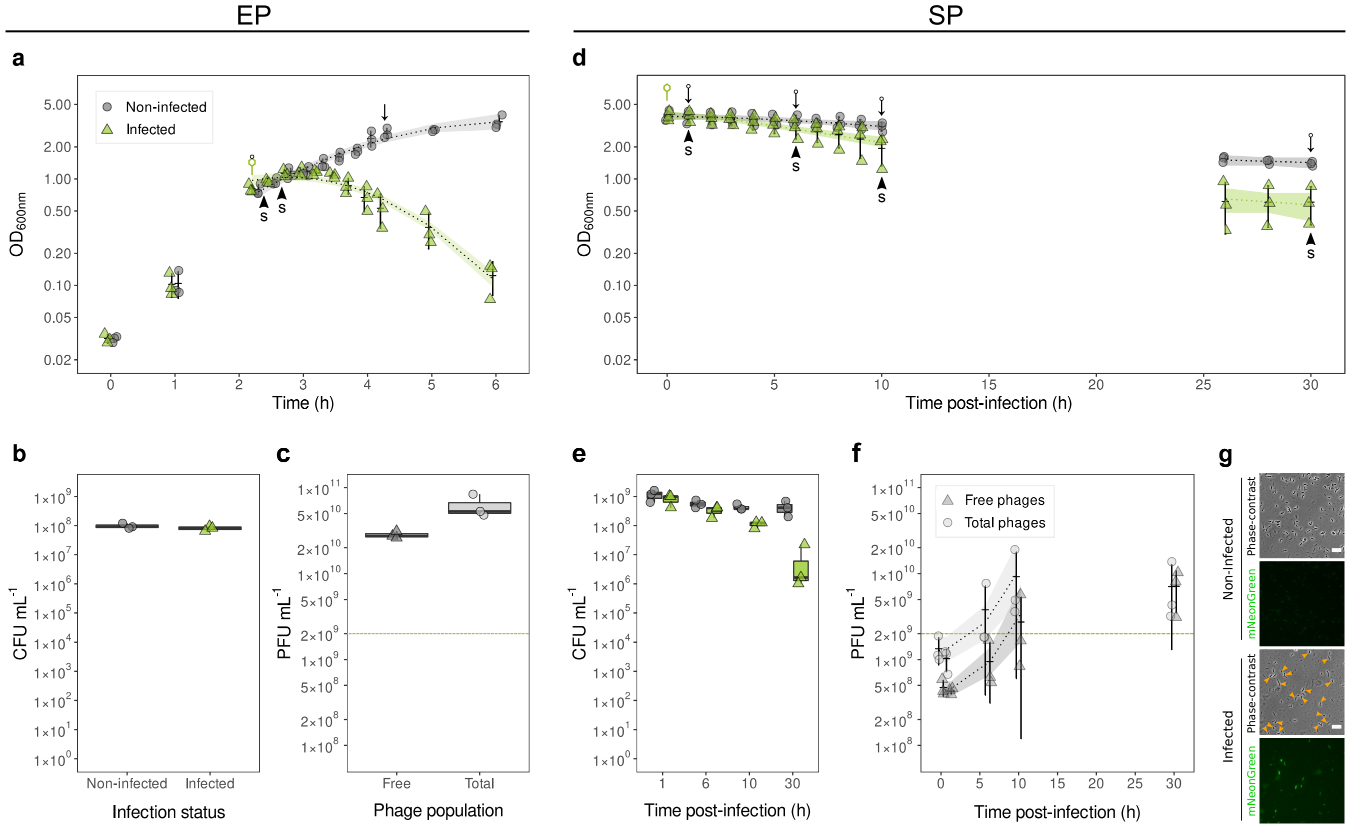


**Supplementary figure 6. Characterization of phage infection cultures used for SPP1 gene expression analyses.** $Bs^{wt}$ grown in flasks were infected with 2×10$^{9}$ PFU ml$^{-1}$ of SPP1$^{mNeonGreen}$ either at mid-EP (**a**) or at 10h post-inoculation in SP (**d**). Green phage symbols denote the times of infection, black arrows denote times of PFU titration, small circles attached to these symbols mark times of CFU sampling and black arrowheads labelled with an 'S' denote times of sampling for RNA and protein isolation. CFU were enumerated at the time of infection during EP (**b**) or throughout SP infection (**e**). PFU were enumerated after 2h of infection in EP (**c**) or throughout infection during SP (**f**). Green dashed lines denote phage input (**c**, **f**). Infected cells were imaged at 30h p.i. during stationary phase infection (**g**). Data are from 3 independent biological replicates. Linear regression models fit to the data for statistical analysis in **a**, **d** and **f** are represented graphically by dotted lines (model fits) and shaded areas (95% confidence intervals). The mean (horizontal bars) and standard deviation (vertical bars) is given in **a**, **d** and **f**. The median, upper and lower quartiles (boxes) and the limits of the data (whiskers) of the data are given in **b**, **c** and **e**. Scale bars in **g** represent 10µm. Data and analyses related to these experiments are presented in **Table S3** and **Supplementary Data Analysis**.

#### Table S1. Bacterial strains, phage strains and plasmids used in this study.

| ID | Description | Source |
|---|---|---|
| | **Bacterial strains** | |
| *Bs*$^{wt}$ | Wild-type *Bacillus subtilis subsp. subtilis str. 168 trpC2* (NCBI genome ID: 665). Strain 1A1$^{WT}$. Contains prophages PBSX and SPβ | BGSC |
| *Bs*$^{Pspac::yueB}$ | *B. subtilis* 1A1$^{WT}$ derivative expressing *yueB* under the control of the IPTG inducible promoter *Pspac* in the native locus. Contains a transcriptional fusion of the 5' end of *yueB* to *spoVG-lacZ*, and an erythromycin resistance gene. Erm$^{R}$ 0.5µg ml$^{-1}$ | This study |
| YB886 | Derivative of wild-type *B. subtilis* subsp. *subtilis* str. 168 *trpC*2 (NCBI genome ID: 665). Cured of prophages PBSX and SPβ. Used for titration and ampification of phage stocks | [1] |
| | **Phage strains** | |
| SPP1$^{wt}$ | Wild-type Subtilis Phage Pavia 1 (SPP1; NCBI genome ID: 4782) | [2] |
| SPP1$^{mNeonGreen}$ | SPP1$^{WT}$ derivative containing a *B. subtilis* codon optimised RBS-mNeonGreen-STOP construct at the unique SphI site of the phage genome (genome pos. 39725, within gene *43*) under control of *P*E2 | This study |
| | **Plasmids** | |
| pEX-A128:: *mNeonGreen* | Plasmid containing *B. subtilis* codon optimised RBS-mNeonGreen-STOP construct used for PCR amplification for construction of SPP1$^{mNeonGreen}$. Confers ampicillin resistance (Amp$^{R}$ 100µg ml$^{-1}$) in *Escherichia coli* | [3] |


[1] Yasbin RE, Fields PI, Andersen BJ. Properties of *Bacillus subtilis* 168 derivatives freed of their natural prophages. *Gene* 1980; **12**: 155–9.

[2] Riva S, Polsinelli M, Falaschi A. A New Phage of *Bacillus subtilis* with Infectious DNA having Separable Strands. *J Mol Biol* 1968; **35**: 347–56.

[3] Bisson-Filho AW, Hsu YP, Squyres GR, Kuru E, Wu F, Jukes C, et al. Treadmilling by FtsZ filaments drives peptidoglycan synthesis and bacterial cell division. *Science* 2017; **355**: 739-43

**Table S2. Primers used in this study.**

| Primer | Sequence | Description | Source |
|---|---|---|---|
| yueB-fw2 | CGGCCGTCCGAAATTCAATC | Forward primer for amplification of insertion region in *B. subtilis* 1A1$^{Pspac::yueB}$ | This study |
| yueB-rv2 | CCGAGCAGGATCAGTACGAC | Reverse primer for amplification of insertion region in *B. subtilis* 1A1$^{Pspac::yueB}$ | This study |
| pspac::yueB_seq-r1 | AGACGGCAAACGACTGTC | | This study |
| pspac::yueB_seq-f1 | ACACCAACGTAACCTATCCC | | This study |
| pspac::yueB_seq-r2 | GTACATCGGGCAAATAATATCGG | | This study |
| pspac::yueB_seq-f2 | GATCGTAATCACCCGAGTG | | This study |
| pspac::yueB_seq-r3 | TCGCCACTTCAACATCAAC | | This study |
| pspac::yueB_seq-f3 | CTAACGCCTGGGTCGAAC | | This study |
| pspac::yueB_seq-r4 | CTGGCACCCAGTTGATCG | | This study |
| pspac::yueB_seq-f4 | TTTCCCGCGTGGTGAAC | Sequencing of *B. subtilis* 1A1$^{Pspac::yueB}$ *spoVG-lacZ::ermR::Pspac::yueB* insertion region | This study |
| pspac::yueB_seq-r5 | CTACGTGACTGGGTCATGG | | This study |
| pspac::yueB_seq-f5 | AGTTAGGCATCGCATCCTG | | This study |
| pspac::yueB_seq-r6 | GCTGAGATAGGTGCCTCAC | | This study |
| pspac::yueB_seq-f6 | GCAAGCAGCAGATTACGC | | This study |
| pspac::yueB_seq-r7 | GGTTTCTTAGACGTCAGGTG | | This study |
| pspac::yueB_seq-f7 | GGAATAAGGGCGACACGG | | This study |
| pspac::yueB_seq-r8 | GGCGTGTTTCATTGCTTG | | This study |
| pspac::yueB_seq-f8 | CAGCGGAATGCTTTCATCC | | This study |
| pspac::yueB_seq-r9 | GCATCCTCAAGCTCACGC | | This study |
| pspac::yueB_seq-f9 | CGCGTTTCGGTGATGAAG | | This study |
| mNeonGreen_SphI-f1 | GGTAGCATGCTAAAGGAGAAATATAC**ATGTCGGCTGGCTCCGCTGC** | Forward primer for amplification of RBS-*mNeonGreen*-STOP construct from pEX-A128::*mNeonGreen*. SphI site underlined, *mNeonGreen* sequence in bold | This study |
| mNeonGreen_Sph1-r1 | AAGAGCATGCATTA**TCACTTATAGAGTTCATCCATACC** | Reverse primer for amplification of RBS-*mNeonGreen*-STOP construct from pEX-A128::*mNeonGreen*. SphI site underlined, *mNeonGreen* sequence in bold | This study |
| SPP1_mNeonGreen-f1 | GGTTTTATGTGTCCGAATTGCG | Forward primer for amplification of insertion region in SPP1$^{mNeonGreen}$ | This study |
| SPP1_mNeonGreen-r1 | CCTGCTCCCAATTGTTCTGC | Forward primer for amplification of insertion region in SPP1$^{mNeonGreen}$ | This study |

| | | | |
|---|---|---|---|
| 3066 | AAAACAGCGGGTGCAGG | Sequencing of SPP1mNeonGreen RBS-*mNeonGreen*-STOP insertion region | This study |
| SPP1_mNeon_seq-1 | CGGTTGACTCGTAATCTGAA | | This study |
| SPP1_mNeon_seq-2 | GGTTCATGCCACTTGTTACC | | This study |
| 1033-Bs-gyrA-F | GAATACGGCAGAACGGCAAA | Forward primer for qPCR : *B. subtilis gyrA* | This study |
| 1034-Bs-gyrA-R | TTCGTTTTGAAACCCCATGC | Reverse primer for qPCR : *B. subtilis gyrA* | This study |
| 1035-Bs-rplP-F | AGCTGCACGTATTGCGATGA | Forward primer for qPCR : *B. subtilis rplP* | This study |
| 1036-Bs-rplP-R | CTTCTGGAGCCCCTTTACCG | Reverse primer for qPCR : *B. subtilis rplP* | This study |
| 1037-Bs-rpsE-F | TTATCGCTGGAGGCCCTGTA | Forward primer for qPCR : *B. subtilis rpsE* | This study |
| 1038-Bs-rpsE-R | AGCTTCGCAACGTCTTCAGC | Reverse primer for qPCR : *B. subtilis rpsE* | This study |
| 1737-Bs-era-F | GTAGCGGCAACGATTGTGGT | Forward primer for qPCR : *B. subtilis era* | This study |
| 1738-Bs-era-R | GTAGACACGGGAGCCCAACA | Reverse primer for qPCR : *B. subtilis era* | This study |
| 1741-Bs-secA-F | GGCGACGATTACGTTCCAAA | Forward primer for qPCR : *B. subtilis secA* | This study |
| 1742-Bs-secA-R | GGGATCGTGACAACCTGCAT | Reverse primer for qPCR : *B. subtilis secA* | This study |
| 1745-Bs-gmk-F | TCCCGGAAGGCCTGTTTATT | Forward primer for qPCR : *B. subtilis gmk* | This study |
| 1746-Bs-gmk-R | TCAGCTTTTGCGGCTTTCAT | Reverse primer for qPCR : *B. subtilis gmk* | This study |
| 1747-Bs-rpoB-F | CGTATGAACATCGGGCAGGT | Forward primer for qPCR : *B. subtilis rpoB* | This study |
| 1748-Bs-rpoB-R | TGCCGGCTTCTTCAAGTGTT | Reverse primer for qPCR : *B. subtilis rpoB* | This study |
| 1767-Bs-dnaG-F | GATCATCACGGGGCTGTTGT | Forward primer for qPCR : *B. subtilis dnaG* | This study |
| 1768-Bs-dnaG-R | CGGGGTTTCAGGACTGTTCA | Reverse primer for qPCR : *B. subtilis dnaG* | This study |
| 1769-Bs-ftsZ-F | CAGTCGGCGTTGTGACAAGA | Forward primer for qPCR : *B. subtilis ftsZ* | This study |
| 1770-Bs-ftsZ-R | GATACGGTCGTTCGGGATCA | Reverse primer for qPCR : *B. subtilis ftsZ* | This study |
| 1029-SPP1-g6-F | CGGGCTGAAATACCTGTGGA | Forward primer for qPCR : SPP1 gene *6* | This study |
| 1030-SPP1-g6-R | TAGCCCCTCCTCCGATTGTT | Reverse primer for qPCR : SPP1 gene *6* | This study |
| 1031-SPP1-g11-F | ACGAGATAGCGGTGCGAAGA | Forward primer for qPCR : SPP1 gene *11* | This study |
| 1032-SPP1-g11-R | TCCTTGTACGCCTCCGTTTG | Reverse primer for qPCR : SPP1 gene *11* | This study |
| 1749-SPP1-g46-F | GACCCAGAGGCTAACCCTTATCA | Forward primer for qPCR : SPP1 gene *46* | This study |
| 1750-SPP1-g46-R | GGCGTTCTCCATAACCCAATG | Reverse primer for qPCR : SPP1 gene *46* | This study |
| 1977-SPP1-g24.1-F | CAGCGTATCAGCTTACTGGAGAGA | Forward primer for qPCR : SPP1 gene *24.1* | This study |
| 1978-SPP1-g24.1-R | CCAAGTGGTGTTTTCATGGATTT | Reverse primer for qPCR : SPP1 gene *24.1* | This study |
| 1981-SPP1-g26-F | ACAGCGCAACAGGTTTATGATG | Forward primer for qPCR : SPP1 gene *26* | This study |
| 1982-SPP1-g26-R | CTTCTCGCTGTTTGCGACCT | Reverse primer for qPCR : SPP1 gene *26* | This study |
| 1983-SPP1-g35-F | ACCAGTCAAGCCGCAGGATA | Forward primer for qPCR : SPP1 gene *35* | This study |
| 1984-SPP1-g35-R | GATGCGGGAAAGTCGATCAG | Reverse primer for qPCR : SPP1 gene *35* | This study |

#### Legend of Supplementary file “Table S3.xlsx”

**Table S3. Supporting information on bacterial growth, phage induced lysis, and PFU production for main Figs. 1 to 4 and 6 and to Supplementary figures 2, 5, and 6.**

**Table S4. SPP1 gene expression during stationary phase (SP) infection compared to exponential phase (EP) late stage of infection.**

| Time post-infection in stationary phase (h) | Gene | Mean relative fold-change ± SD [a] | Mean fold-reduction ± SD [b] |
|---|---|---|---|
| 1 | *46* | 0.153 ± 0.0436 | 6.85 ± 1.68 |
| 6 | *46* | 0.974 ± 0.663 | 1.32 ± 0.666 |
| 10 | *46* | 3.83 ± 4.23 | -3.83 ± 4.23 |
| 30 | *46* | 7.03 ± 4.98 | -7.03 ± 4.98 |
| 1 | *35* | 0.0626 ± 0.0191 | 17.2 ± 5.96 |
| 6 | *35* | 0.445 ± 0.131 | 2.41 ± 0.832 |
| 10 | *35* | 1.46 ± 1.23 | -1.46 ± 1.23 |
| 30 | *35* | 17.4 ± 13.1 | -17.4 ± 13.1 |
| 1 | *24.1* | 0.00812 ± 0.00520 | 187 ± 158 |
| 6 | *24.1* | 0.142 ± 0.0579 | 7.75 ± 2.72 |
| 10 | *24.1* | 0.503 ± 0.391 | 3.74 ± 3.74 |
| 30 | *24.1* | 0.0685 ± 0.0418 | 22.1 ± 19.0 |
| 1 | *26* | 0.00762 ± 0.00469 | 211 ± 196 |
| 6 | *26* | 0.107 ± 0.0652 | 11.5 ± 5.32 |
| 10 | *26* | 0.468 ± 0.392 | 6.39 ± 8.31 |
| 30 | *26* | 0.178 ± 0.124 | 11.2 ± 12.3 |
| 1 | *6* | 0.00600 ± 0.00403 | 323 ± 350 |
| 6 | *6* | 0.0547 ± 0.0336 | 23.2 ± 12.7 |
| 10 | *6* | 0.245 ± 0.283 | 10.9 ± 10.9 |
| 30 | *6* | 0.0370 ± 0.0286 | 37.2 ± 19.9 |
| 1 | *11* | 0.00300 ± 0.00223 | 619 ± 624 |
| 6 | *11* | 0.0501 ± 0.0310 | 25.9 ± 15.2 |
| 10 | *11* | 0.182 ± 0.176 | 11.5 ± 11.1 |
| 30 | *11* | 0.0235 ± 0.0223 | 73.7 ± 53.0 |

[a] Mean relative fold-change using late exponential phase (25min post-infection) as the calibrator.

[b] Mean fold reduction relative to late exponential phase infection. The reciprocals of relative fold-change values less than 1 are given. Negative values represent a fold-increase relative to late exponential phase infection.

Index

#### Supplementary data analysis

This data analysis summary includes summary statistics, statistical models used, their outputs and any multiple comparison testing made for each experiment. Data are organised by Figure and their respective panels. All model / test outputs are as given by R, minimally edited to make interpretation easier for the reader.

==============================================================

#### Contents:


| Figure | Panel | Pages |
|---|---|---|
| 1 | b | 24-26 |
| 1 | c | 26 |
| 1 | d | 26-27 |
| 1 | e | 27 |
| 2 | a | 28-31 |
| 2 | b | 31 |
| 3 | a | 32-33 |
| 3 | b | 33-34 |
| 3 | c | 34-35 |
| 3 | e | 35 |
| 4 | a | 36 |
| 4 | b | 36-40 |
| 4 | c | 41-44 |
| 4 | d | 44-45 |
| 4 | e | 46 |
| 5 | b | 47-49 |
| 6 | b | 50-55 |
| 6 | c | 55 |
| 6 | d | 56 |
| S2 | a | 57 |
| S2 | b | 58 |
| S2 | d | 58-59 |
| S5 | a | 60-61 |
| S5 | c | 61-62 |
| S6 | a | 63 |
| S6 | b | 63 |
| S6 | c | 63 |
| S6 | d | 64-65 |
| S6 | e | 65 |
| S6 | f | 66 |


#### Abbreviations (inc. model terms):

| Abbreviation | Meaning | Notes |
|---|---|---|
| bp | breakpoint for multiphasic lysis rates | |
| CFU | colony forming units per ml-1 | |
| dur | infection duration | determined by lysis rate |
| fch / log10fch | fold change | raw / log10 transformation |
| gene | phage gene | |
| gphase | growth phase | |
| gphasepd | combined variable: gphase, pd | |
| infstat | infection status | |
| ISSP | combined variable: infstat, time.sup, pd | |
| len | cell length | |
| maxima | number of maxima per cell-1 | |
| max | maximum | |
| min | minimum | |
| mxdecrt | maximum rate of decline | |
| mxpfurt | maximum rate of PFU production | |
| NA | not applicable | |
| ND | not determined | |
| p.i. | post-infection | |
| pd | period | |
| percdec / propdec | proportion decrease in OD600 | percentage / proportion |
| percNF / propNF | proportion of cells without foci | percentage / proportion |
| percrgw / proprgw | proportion of cell regrowth | percentage / proportion |
| percspr / propspr | proportion sporulating cells | percentage / proportion |
| PFU | plaque forming units per ml-1 | |
| ppn | phage population | |
| r0 | growth rate | |
| strain | bacterial strain | |
| suppd | combined variable: time.sup, pd | |
| surf | cell surface area | |
| time / time.fac | time | continuous / categorical |
| time.sup | time of supplementation | |
| time50 | time to reach 50% maximum OD600 | determined by loss OD600 |
| timelysinit | time to initiation of lysis | |
| tmxdec | time at max. rate decline | |
| tmxpfu | time at max. rate of PFU production | |
| tpd | time period | |
| vol | cell volume | |
| wid | cell width | |

Fig. 1

### Figure 1

#### *Fig. 1b*

##### OD~time*infection_status relationships

**Model 2.25h (top left facet)**

```
Call:
glm(formula = od ~ time + I(time^2) * infstat, family =
Gamma(link = log))

Deviance Residuals:
     Min        1Q    Median        3Q       Max
-0.39559  -0.15596  -0.00678   0.11104   0.51466

Coefficients:
                    Estimate Std. Error t value Pr(>|t|)
(Intercept)         -1.902e-01  8.560e-02  -2.222  0.03375 *
time                 9.792e-03  3.041e-03   3.220  0.00301 **
I(time^2)           -1.153e-04  2.488e-05  -4.633 6.13e-05 ***
infstatnon           6.003e-03  1.150e-01   0.052  0.95870
I(time^2):infstatnon 1.209e-04  1.402e-05   8.622 9.75e-10 ***
---
Signif. codes:  0 ‘***’ 0.001 ‘**’ 0.01 ‘*’ 0.05 ‘.’ 0.1 ‘ ’ 1

(Dispersion parameter for Gamma family taken to be 0.04317806)
Null deviance: 8.9087  on 35  degrees of freedom
Residual deviance: 1.3169  on 31  degrees of freedom
AIC: -8.4459
Number of Fisher Scoring iterations: 6
```

**Analysis of Deviance Table (top left facet)**

```
Model: Gamma, link: log
Response: od
Terms added sequentially (first to last)
                 Df Deviance Resid. Df Resid. Dev  Pr(>Chi)
NULL                               35     8.9087
time              1   0.7327       34     8.1760 3.799e-05 ***
I(time^2)         1   0.0125       33     8.1635    0.5904
infstat           1   3.6064       32     4.5571 < 2.2e-16 ***
I(time^2):infstat 1   3.2401       31     1.3169 < 2.2e-16 ***
---
Signif. codes:  0 ‘***’ 0.001 ‘**’ 0.01 ‘*’ 0.05 ‘.’ 0.1 ‘ ’ 1
```

**Model 4h (top right facet)**

```
Call:
glm(formula = od ~ time + I(time^2) * infstat, family =
Gamma(link = log))

Deviance Residuals:
     Min        1Q    Median        3Q       Max
-0.36881  -0.06007  -0.01226   0.09207   0.29076

Coefficients:
                      Estimate Std. Error t value Pr(>|t|)
(Intercept)          1.002e+00  5.887e-02  17.020  < 2e-16 ***
time                -1.522e-03  2.091e-03  -0.728  0.47235
I(time^2)           -4.988e-05  1.711e-05  -2.915  0.00655 **
infstatnon           8.885e-02  7.907e-02   1.124  0.26976
I(time^2):infstatnon 7.637e-05  9.644e-06   7.919 6.13e-09 ***
---
Signif. codes:  0 ‘***’ 0.001 ‘**’ 0.01 ‘*’ 0.05 ‘.’ 0.1 ‘ ’ 1

(Dispersion parameter for Gamma family taken to be 0.02041815)
Null deviance: 4.63566  on 35  degrees of freedom
Residual deviance: 0.65489  on 31  degrees of freedom
AIC: 25.662
Number of Fisher Scoring iterations: 5
```

**Analysis of Deviance Table (top right facet)**

```
Model: Gamma, link: log
Response: od
Terms added sequentially (first to last)
                  Df Deviance Resid. Df Resid. Dev  Pr(>Chi)
NULL                                35     4.6357
time               1  0.63225       34     4.0034 2.627e-08 ***
I(time^2)          1  0.06750       33     3.9359   0.06904 .
infstat            1  1.98858       32     1.9473 < 2.2e-16 ***
I(time^2):infstat  1  1.29244       31     0.6549 1.776e-15 ***
---
Signif. codes:  0 ‘***’ 0.001 ‘**’ 0.01 ‘*’ 0.05 ‘.’ 0.1 ‘ ’ 1
```

**Model 6h  (bottom left facet)**

```
Call:
glm(formula = od ~ time + infstat, family = gaussian(link =
"inverse"))

Deviance Residuals:
     Min        1Q    Median        3Q       Max
-0.39902  -0.14552   0.04722   0.11826   0.31594

Coefficients:
              Estimate Std. Error t value Pr(>|t|)
(Intercept)  2.874e-01  4.446e-03  64.651  < 2e-16 ***
time        -2.016e-04  5.462e-05  -3.690 0.000803 ***
infstatnon  -1.477e-02  5.010e-03  -2.947 0.005846 **
---
Signif. codes:  0 ‘***’ 0.001 ‘**’ 0.01 ‘*’ 0.05 ‘.’ 0.1 ‘ ’ 1

(Dispersion parameter for gaussian family taken to be
0.03502253)
Null deviance: 1.9109  on 35  degrees of freedom
Residual deviance: 1.1557  on 33  degrees of freedom
AIC: -13.632
Number of Fisher Scoring iterations: 4
```

**Analysis of Deviance Table**

```
Model: gaussian, link: inverse
Response: od
Terms added sequentially (first to last)
        Df Deviance Resid. Df Resid. Dev  Pr(>Chi)
NULL                       35     1.9109
time     1  0.46598        34     1.4449 0.0002647 ***
infstat  1  0.28920        33     1.1557 0.0040584 **
---
Signif. codes:  0 ‘***’ 0.001 ‘**’ 0.01 ‘*’ 0.05 ‘.’ 0.1 ‘ ’ 1
```

**Model 10h  (bottom right facet)**

```
Call:
glm(formula = od ~ time + infstat, family = gaussian(link =
"inverse"))

Deviance Residuals:
    Min       1Q   Median       3Q      Max
-1.1003  -0.7453   0.3542   0.4551   0.5686

Coefficients:
              Estimate Std. Error t value Pr(>|t|)
(Intercept)  0.2536401  0.0129723  19.552   <2e-16 ***
time         0.0003770  0.0001811   2.082   0.0452 *
infstatnon  -0.0157060  0.0163467  -0.961   0.3436
---
Signif. codes:  0 ‘***’ 0.001 ‘**’ 0.01 ‘*’ 0.05 ‘.’ 0.1 ‘ ’ 1

(Dispersion parameter for gaussian family taken to be 0.3820679)
Null deviance: 14.669  on 35  degrees of freedom
Residual deviance: 12.608  on 33  degrees of freedom
AIC: 72.393
Number of Fisher Scoring iterations: 5
```

**Analysis of Deviance Table**

```
Model: gaussian, link: inverse
Response: od
Terms added sequentially (first to last)
        Df Deviance Resid. Df Resid. Dev Pr(>Chi)
NULL                       35     14.669
time     1  1.72745        34     12.942  0.03348 *
infstat  1  0.33374        33     12.608  0.34999
---
Signif. codes:  0 ‘***’ 0.001 ‘**’ 0.01 ‘*’ 0.05 ‘.’ 0.1 ‘ ’ 1
```

#### Fig. 1

#### Growth rates at selected time-points (**Table S3,** also relates to panel 1d)

**Model**
```
Call:
lm(formula = log(r0 + 1) ~ time.fac)

Residuals:
      Min        1Q    Median        3Q       Max 
-0.096450 -0.011084 -0.002757  0.032061  0.065352 

Coefficients:
            Estimate Std. Error t value Pr(>|t|)    
(Intercept)  0.80840    0.03086   26.20 1.51e-10 ***
time.fac4   -0.47871    0.04364  -10.97 6.77e-07 ***
time.fac6   -0.74530    0.04364  -17.08 9.99e-09 ***
time.fac10  -0.84779    0.04364  -19.43 2.85e-09 ***
time.fac28  -0.82609    0.04364  -18.93 3.67e-09 ***
---
Signif. codes:  0 ‘***’ 0.001 ‘**’ 0.01 ‘*’ 0.05 ‘.’ 0.1 ‘ ’ 1

Residual standard error: 0.05345 on 10 degrees of freedom
Multiple R-squared:  0.9815,  Adjusted R-squared:  0.9742 
F-statistic: 132.9 on 4 and 10 DF,  p-value: 1.268e-08
```

**Analysis of Variance Table**
```
Response: log(r0 + 1)
          Df  Sum Sq Mean Sq F value    Pr(>F)    
time.fac   4 1.51878 0.37969   132.9 1.268e-08 ***
Residuals 10 0.02857 0.00286                      
---
Signif. codes:  0 ‘***’ 0.001 ‘**’ 0.01 ‘*’ 0.05 ‘.’ 0.1 ‘ ’ 1
```

**Simultaneous Tests for General Linear Hypotheses**
```
 contrast  ratio     SE df null t.ratio p.value
 2.25 / 4  1.614 0.0704 10    1  10.969  <.0001
 2.25 / 6  2.107 0.0920 10    1  17.078  <.0001
 2.25 / 10 2.334 0.1019 10    1  19.426  <.0001
 2.25 / 28 2.284 0.0997 10    1  18.929  <.0001
 4 / 6     1.306 0.0570 10    1   6.109  0.0008
 4 / 10    1.446 0.0631 10    1   8.457  0.0001
 4 / 28    1.415 0.0618 10    1   7.960  0.0001
 6 / 10    1.108 0.0484 10    1   2.348  0.2070
 6 / 28    1.084 0.0473 10    1   1.851  0.3989
 10 / 28   0.979 0.0427 10    1  -0.497  0.9858

P value adjustment: tukey method for comparing a family of 5
estimates
Tests are performed on the log scale
```

#### Maximum decline rates (**Table S3**)

**Model**
```
Call:
lm(formula = mxdecrt ~ time.fac)

Residuals:
     Min       1Q   Median       3Q      Max 
-0.33989 -0.22618 -0.08246  0.13689  0.56482 

Coefficients:
             Estimate Std. Error t value Pr(>|t|)    
(Intercept)  -1.31021    0.18082  -7.246 8.84e-05 ***
time.fac4    -0.03702    0.25571  -0.145  0.88846    
time.fac6     1.20664    0.25571   4.719  0.00150 ** 
time.fac10    1.13550    0.25571   4.441  0.00217 ** 
---
Signif. codes:  0 ‘***’ 0.001 ‘**’ 0.01 ‘*’ 0.05 ‘.’ 0.1 ‘ ’ 1

Residual standard error: 0.3132 on 8 degrees of freedom
Multiple R-squared:  0.8443,  Adjusted R-squared:  0.7859 
F-statistic: 14.46 on 3 and 8 DF,  p-value: 0.001353
```

**Analysis of Variance Table**
```
Response:  mxdecrt
          Df Sum Sq Mean Sq F value   Pr(>F)   
time.fac   3 4.2550 1.41832   14.46 0.001353 **
Residuals  8 0.7847 0.09808                    
---
Signif. codes:  0 ‘***’ 0.001 ‘**’ 0.01 ‘*’ 0.05 ‘.’ 0.1 ‘ ’ 1
```

**Simultaneous Tests for General Linear Hypotheses**
```
Multiple Comparisons of Means: Tukey Contrasts
Fit: lm(formula = mxdecrt ~ time.fac)
Linear Hypotheses:
contrast       Estimate Std. Error t value Pr(>|t|)   
4 - 2.25 == 0  -0.03702    0.25571  -0.145  0.99882   
6 - 2.25 == 0   1.20664    0.25571   4.719  0.00666 **
10 - 2.25 == 0  1.13550    0.25571   4.441  0.00927 **
6 - 4 == 0      1.24367    0.25571   4.864  0.00562 **
10 - 4 == 0     1.17252    0.25571   4.585  0.00765 **
10 - 6 == 0    -0.07114    0.25571  -0.278  0.99188   
---
Signif. codes:  0 ‘***’ 0.001 ‘**’ 0.01 ‘*’ 0.05 ‘.’ 0.1 ‘ ’ 1
(Adjusted p values reported -- single-step method)
```

#### Time at maximum decline rates (**Table S3**)

**Model**
```
Call:
lm(formula = tmxdec ~ time.fac, data = maxR.comb)
Residuals:
   Min     1Q Median     3Q    Max 
   -60      0      0     20     30 

Coefficients:
              Estimate Std. Error t value Pr(>|t|)  
(Intercept)   6.000e+01  1.803e+01   3.328   0.0104 *
time.fac4    -1.540e-14  2.550e+01   0.000   1.0000  
time.fac6     2.500e+01  2.550e+01   0.981   0.3555  
time.fac10    1.500e+01  2.550e+01   0.588   0.5725  
---
Signif. codes:  0 ‘***’ 0.001 ‘**’ 0.01 ‘*’ 0.05 ‘.’ 0.1 ‘ ’ 1

Residual standard error: 31.22 on 8 degrees of freedom
Multiple R-squared:  0.1475,  Adjusted R-squared:  -0.1721 
F-statistic: 0.4615 on 3 and 8 DF,  p-value: 0.7168
```

**Analysis of Variance Table**
```
Response:  tmxdec
          Df Sum Sq Mean Sq F value Pr(>F)
time.fac   3   1350     450  0.4615 0.7168
Residuals  8   7800     975               
```

#### Time to lysis initiation (**Table S3**)

**Welch Two Sample t-test**
```
t = 1.4142, df = 4, p-value = 0.2302
alternative hypothesis: true difference in means is not equal to 0
95 percent confidence interval:
 -0.1605405  0.4938739
sample estimates:
mean of x mean of y 
0.8333333 0.6666667 
```

### Fig. 1

#### % decrease OD (**Table S3**)

```
Model
NOTE: analysed as proportions rather than percentages to allow
arcsine transformation

Call:
lm(formula = asin(propdec) ~ time.fac)

Residuals:
     Min       1Q   Median       3Q      Max 
-0.18430 -0.05470 -0.00522  0.04329  0.15472 

Coefficients:
             Estimate Std. Error t value Pr(>|t|)    
(Intercept)   0.53606    0.06249   8.578 2.63e-05 ***
time.fac4     0.03865    0.08838   0.437 0.673450    
time.fac6    -0.48305    0.08838  -5.466 0.000598 ***
time.fac10   -0.33306    0.08838  -3.769 0.005477 ** 
---
Signif. codes:  0 '***' 0.001 '**' 0.01 '*' 0.05 '.' 0.1 ' ' 1

Residual standard error: 0.1082 on 8 degrees of freedom
Multiple R-squared:  0.8617,  Adjusted R-squared:  0.8098 
F-statistic: 16.61 on 3 and 8 DF,  p-value: 0.0008493
```

```
Analysis of Variance Table
Response: asin(propdec)
          Df  Sum Sq  Mean Sq F value    Pr(>F)    
time.fac   3 0.58395 0.194650  16.613 0.0008493 ***
Residuals  8 0.09373 0.011716                      
---
Signif. codes:  0 '***' 0.001 '**' 0.01 '*' 0.05 '.' 0.1 ' ' 1

Simultaneous Tests for General Linear Hypotheses
Multiple Comparisons of Means: Tukey Contrasts
Fit: lm(formula = asin(propdec) ~ tpt.fac)
Linear Hypotheses:
contrast       Estimate Std. Error t value Pr(>|t|)   
4 - 2.25 == 0   0.03865    0.08838   0.437  0.97026   
6 - 2.25 == 0  -0.48305    0.08838  -5.466  0.00264 **
10 - 2.25 == 0 -0.33306    0.08838  -3.769  0.02296 * 
6 - 4 == 0     -0.52170    0.08838  -5.903  0.00169 **
10 - 4 == 0    -0.37171    0.08838  -4.206  0.01254 * 
10 - 6 == 0     0.14999    0.08838   1.697  0.38426   
---
Signif. codes:  0 '***' 0.001 '**' 0.01 '*' 0.05 '.' 0.1 ' ' 1
(Adjusted p values reported -- single-step method)
```

---

#### *Fig. 1c*
#### Fold change in PFU **(Table S3)**

```
Model
Call:
glm(formula = fch ~ time.fac, family = Gamma(link = log))

Deviance Residuals: 
    Min       1Q   Median       3Q      Max  
-1.2950  -0.5412   0.2406   0.3236   0.5030  

Coefficients:
            Estimate Std. Error t value Pr(>|t|)    
(Intercept)   3.2677     0.3410   9.583 1.17e-05 ***
time.fac4    -1.5599     0.4822  -3.235    0.012 *  
time.fac6    -3.6320     0.4822  -7.532 6.72e-05 ***
time.fac10   -4.0016     0.4822  -8.298 3.35e-05 ***
---
Signif. codes:  0 '***' 0.001 '**' 0.01 '*' 0.05 '.' 0.1 ' ' 1

(Dispersion parameter for Gamma family taken to be 0.3488201)
Null deviance: 31.9048  on 11  degrees of freedom
Residual deviance:  4.5653  on  8  degrees of freedom
AIC: 51.912
Number of Fisher Scoring iterations: 5
```

```
Analysis of Deviance Table
Model: Gamma, link: log
Response: fch
Terms added sequentially (first to last)

         Df Deviance Resid. Df Resid. Dev  Pr(>Chi)    
NULL                        11     31.905              
time.fac  3    27.34         8      4.565 < 2.2e-16 ***
---
Signif. codes:  0 '***' 0.001 '**' 0.01 '*' 0.05 '.' 0.1 ' ' 1

Simultaneous Tests for General Linear Hypotheses
 contrast  ratio     SE df null t.ratio p.value
 2.25 / 4   4.76  2.295  8    1   3.235  0.0478
 2.25 / 6  37.79 18.223  8    1   7.532  0.0003
 2.25 / 10 54.69 26.372  8    1   8.298  0.0002
 4 / 6      7.94  3.830  8    1   4.297  0.0112
 4 / 10    11.49  5.542  8    1   5.063  0.0043
 6 / 10     1.45  0.698  8    1   0.767  0.8673

P value adjustment: tukey method for comparing a family of 4
estimates
Tests are performed on the log scale
```

---

#### *Fig. 1d*
#### OD~time*infection status relationship

```
Model (panel)
Call:
lm(formula = od ~ time * infstat)

Residuals:
     Min       1Q   Median       3Q      Max 
-0.50281 -0.09882 -0.01904  0.08289  0.79430 
Coefficients:
                  Estimate Std. Error t value Pr(>|t|)    
(Intercept)       5.560367   0.110667  50.244  < 2e-16 ***
time             -0.122889   0.006071 -20.243  < 2e-16 ***
infstatnon        0.835612   0.136669   6.114 1.45e-07 ***
time:infstatnon  -0.094288   0.008075 -11.676 6.79e-16 ***
---
Signif. codes:  0 '***' 0.001 '**' 0.01 '*' 0.05 '.' 0.1 ' ' 1

Residual standard error: 0.1902 on 50 degrees of freedom
Multiple R-squared:  0.9765,  Adjusted R-squared:  0.9751 
F-statistic: 692.7 on 3 and 50 DF,  p-value: < 2.2e-16
```

```
Analysis of Variance Table
Response: od
             Df Sum Sq Mean Sq F value    Pr(>F)    
time          1 65.846  65.846 1819.53 < 2.2e-16 ***
infstat       1  4.420   4.420  122.13 5.004e-15 ***
time:infstat  1  4.934   4.934  136.34 6.788e-16 ***
Residuals    50  1.809   0.036                      
---
Signif. codes:  0 '***' 0.001 '**' 0.01 '*' 0.05 '.' 0.1 ' ' 1
```

Fig. 1

#### Maximum decline rates & times at max decline rates **(Table S3)**

```
Welch Two Sample t-test (Max decline rate)
t = 4.7089, df = 3.2974, p-value = 0.01462
alternative hypothesis: true difference in means is not equal to 0
95 percent confidence interval:
 0.0428848 0.1971202
sample estimates:
  mean of x   mean of y
-0.09221581 -0.21221829
```

```
Wilcoxon rank sum test with continuity correction (Time at max decline rate)
W = 0, p-value = 0.05935
alternative hypothesis: true location shift is not equal to 0
```

#### Time to lysis **(Table S3)**

Not determined – rate already negative from start

#### % decrease OD (**Table S3**)

```
Wilcoxon rank sum test
W = 0, p-value = 0.1
alternative hypothesis: true location shift is not equal to 0
```

---

***Fig. 1e***

#### Fold change in PFU (**Table S3**)

```
Welch Two Sample t-test
NOTE: data were log-transformed to conform to assumptions for analysis
t = -8.7877, df = 3.2861, p-value = 0.002158
alternative hypothesis: true difference in means is not equal to 0
95 percent confidence interval:
 -3.836968 -1.868797
sample estimates:
mean of x mean of y
-1.151702  1.701181
```

=====================================================================

### Figure 2

#### *Fig. 2a*

##### Growth rate at infection (**Table S3**)

```
Model
Call:
glm(formula = r0 ~ gphase, family = gaussian(link = "inverse"))

Deviance Residuals:
       Min          1Q      Median          3Q         Max
-0.0216326  -0.0051915  -0.0007281   0.0045066   0.0239647

Coefficients:
               Estimate Std. Error t value Pr(>|t|)
(Intercept)      8.2563     0.5344  15.450 3.06e-07 ***
gphaseexp       -6.8510     0.5346 -12.815 1.30e-06 ***
gphaselTP        2.8023     1.0976   2.553   0.0340 *
gphasestat      17.9200     5.3954   3.321   0.0105 *
---
Signif. codes:  0 ‘***’ 0.001 ‘**’ 0.01 ‘*’ 0.05 ‘.’ 0.1 ‘ ’ 1

(Dispersion parameter for gaussian family taken to be
0.0001843839)
Null deviance: 0.9003710  on 11  degrees of freedom
Residual deviance: 0.0014748  on  8  degrees of freedom
AIC: -63.995
Number of Fisher Scoring iterations: 4
```

```
Analysis of Deviance Table
Model: gaussian, link: inverse
Response:  r0
Terms added sequentially (first to last)
       Df Deviance Resid. Df Resid. Dev  Pr(>Chi)
NULL                      11    0.90037
gphase   3   0.8989        8    0.00147 < 2.2e-16 ***
---
Signif. codes:  0 ‘***’ 0.001 ‘**’ 0.01 ‘*’ 0.05 ‘.’ 0.1 ‘ ’ 1


Simultaneous Tests for General Linear Hypotheses
 contrast     estimate     SE df t.ratio p.value
 eTP - exp     -0.5905 0.0111  8 -53.258  <.0001
 eTP - lTP      0.0307 0.0111  8   2.768  0.0923
 eTP - stat     0.0829 0.0111  8   7.481  0.0003
 exp - lTP      0.6212 0.0111  8  56.027  <.0001
 exp - stat     0.6734 0.0111  8  60.754  <.0001
 lTP - stat     0.0522 0.0111  8   4.712  0.0066

P value adjustment: tukey method for comparing a family of 4
estimates
```

##### Time to lysis initiation (**Table S3**)

```
Model (Table S3)
Call:
glm(formula = timelysinit ~ gphase, family = Gamma(link =
"inverse"))

Deviance Residuals:
     Min        1Q    Median        3Q       Max
-0.39361  -0.09987   0.00963   0.11533   0.24519
Coefficients:
                Estimate Std. Error t value Pr(>|t|)
(Intercept)       0.9231     0.1167   7.907 4.75e-05 ***
gphaseexp         0.9717     0.2665   3.645  0.00654 **
gphaselTP        -0.3606     0.1367  -2.638  0.02982 *
gphasestat       -0.3858     0.1351  -2.856  0.02128 *
---
Signif. codes:  0 ‘***’ 0.001 ‘**’ 0.01 ‘*’ 0.05 ‘.’ 0.1 ‘ ’ 1

(Dispersion parameter for Gamma family taken to be 0.04798035)
Null deviance: 3.11503  on 11  degrees of freedom
Residual deviance: 0.41362  on  8  degrees of freedom
AIC: 7.1223
Number of Fisher Scoring iterations: 4
```

```
Analysis of Deviance Table
Model: Gamma, link: inverse
Response:  timelysinit
Terms added sequentially (first to last)
       Df Deviance Resid. Df Resid. Dev  Pr(>Chi)
NULL                      11    3.11503
gphase   3   2.7014        8    0.41362 3.621e-12 ***
---
Signif. codes:  0 ‘***’ 0.001 ‘**’ 0.01 ‘*’ 0.05 ‘.’ 0.1 ‘ ’ 1


Simultaneous Tests for General Linear Hypotheses
 contrast     estimate    SE df t.ratio p.value
 eTP - exp      0.5556 0.152  8   3.645  0.0270
 eTP - lTP     -0.6944 0.263  8  -2.638  0.1109
 eTP - stat    -0.7778 0.272  8  -2.856  0.0815
 exp - lTP     -1.2500 0.235  8  -5.330  0.0031
 exp - stat    -1.3333 0.245  8  -5.450  0.0027
 lTP - stat    -0.0833 0.325  8  -0.256  0.9936

P value adjustment: tukey method for comparing a family of 4
estimates
```

#### Fig. 2

### Maximum decline rates & times at max decline rates (**Table S3**)

**Breakpoints for multiphasic decline rate calculation**

| growth phase | initiation of lysis (h) | bp 1 (h) | bp 2 (h) | bp 3 (h) | bp 4 (h) | replicate |
|---|---|---|---|---|---|---|
| EP | 3.50 | 7.88 | NA | NA | NA | 1 |
| EP | 3.50 | 8.50 | NA | NA | NA | 2 |
| EP | 3.58 | 7.88 | NA | NA | NA | 3 |
| eTP | 7.33 | 9.79 | 13.3 | 17.4 | 23.2 | 1 |
| eTP | 7.00 | 9.83 | 13.1 | 17.7 | 22.7 | 2 |
| eTP | 6.92 | 9.71 | 13.0 | 17.8 | 22.4 | 3 |
| lTP | 9.17 | 15.00 | 27.3 | NA | NA | 1 |
| lTP | 10.25 | 16.29 | 24.5 | NA | NA | 2 |
| lTP | 9.92 | 16.29 | 25.3 | NA | NA | 3 |
| SP | 12.42 | 21.25 | 27.6 | NA | NA | 1 |
| SP | 13.00 | 21.92 | 27.6 | NA | NA | 2 |
| SP | 13.17 | 21.13 | 26.7 | NA | NA | 3 |

* periods noted in Table S3 (1-4) correspond to time periods between the above listed breakpoints

```
Model (Max decline rate)
NOTE: growth phase and lysis period combined into one variable
to directly compare lysis rates

Call:
lm(formula = log(mxdecrt * -1) ~ gphasepd)

Residuals:
     Min       1Q   Median       3Q      Max
-0.63749 -0.08795 -0.03175  0.09431  0.42841

Coefficients:
               Estimate Std. Error t value Pr(>|t|)
(Intercept)    -2.16746    0.15614 -13.881 9.96e-12 ***
gphasepdeTP2    0.44260    0.22082   2.004 0.058760 .
gphasepdeTP3   -0.16149    0.22082  -0.731 0.473050
gphasepdeTP4    0.24789    0.22082   1.123 0.274899
gphasepdexp1    2.36518    0.22082  10.711 9.85e-10 ***
gphasepdlTP1   -0.07195    0.22082  -0.326 0.747924
gphasepdlTP2    0.57047    0.22082   2.583 0.017750 *
gphasepdnon1   -0.88323    0.22082  -4.000 0.000704 ***
gphasepdstat1   0.17215    0.22082   0.780 0.444755
gphasepdstat2   0.92205    0.22082   4.176 0.000467 ***
---
Signif. codes:  0 ‘***’ 0.001 ‘**’ 0.01 ‘*’ 0.05 ‘.’ 0.1 ‘ ’ 1

Residual standard error: 0.2704 on 20 degrees of freedom
Multiple R-squared:  0.9309,  Adjusted R-squared:  0.8998
F-statistic: 29.94 on 9 and 20 DF,  p-value: 1.081e-09


Analysis of Variance Table
Response: log(mxdecrt * -1)
          Df  Sum Sq Mean Sq F value    Pr(>F)
gphasepd   9 19.7081 2.18979   29.94 1.081e-09 ***
Residuals 20  1.4628 0.07314
---
Signif. codes:  0 ‘***’ 0.001 ‘**’ 0.01 ‘*’ 0.05 ‘.’ 0.1 ‘ ’ 1
```

```
Simultaneous Tests for General Linear Hypotheses
 contrast          ratio     SE df null  t.ratio p.value
 eTP1 / eTP2      0.6424 0.1418 20    1  -2.004  0.6057
 eTP1 / eTP3      1.1753 0.2595 20    1   0.731  0.9989
 eTP1 / eTP4      0.7804 0.1723 20    1  -1.123  0.9761
 eTP1 / exp1      0.0939 0.0207 20    1 -10.711  <.0001
 eTP1 / lTP1      1.0746 0.2373 20    1   0.326  1.0000
 eTP1 / lTP2      0.5653 0.1248 20    1  -2.583  0.2863
 eTP1 / non1      2.4187 0.5341 20    1   4.000  0.0192
 eTP1 / stat1     0.8419 0.1859 20    1  -0.780  0.9981
 eTP1 / stat2     0.3977 0.0878 20    1  -4.176  0.0132
 eTP2 / eTP3      1.8296 0.4040 20    1   2.736  0.2244
 eTP2 / eTP4      1.2150 0.2683 20    1   0.882  0.9954
 eTP2 / exp1      0.1462 0.0323 20    1  -8.707  <.0001
 eTP2 / lTP1      1.6729 0.3694 20    1   2.330  0.4126
 eTP2 / lTP2      0.8800 0.1943 20    1  -0.579  0.9998
 eTP2 / non1      3.7653 0.8314 20    1   6.004  0.0002
 eTP2 / stat1     1.3106 0.2894 20    1   1.225  0.9592
 eTP2 / stat2     0.6191 0.1367 20    1  -2.171  0.5042
 eTP3 / eTP4      0.6641 0.1466 20    1  -1.854  0.6967
 eTP3 / exp1      0.0799 0.0176 20    1 -11.442  <.0001
 eTP3 / lTP1      0.9144 0.2019 20    1  -0.405  1.0000
 eTP3 / lTP2      0.4810 0.1062 20    1  -3.315  0.0785
 eTP3 / non1      2.0580 0.4544 20    1   3.268  0.0858
 eTP3 / stat1     0.7163 0.1582 20    1  -1.511  0.8726
 eTP3 / stat2     0.3384 0.0747 20    1  -4.907  0.0027
 eTP4 / exp1      0.1204 0.0266 20    1  -9.588  <.0001
 eTP4 / lTP1      1.3769 0.3040 20    1   1.448  0.8967
 eTP4 / lTP2      0.7243 0.1599 20    1  -1.461  0.8922
 eTP4 / non1      3.0991 0.6843 20    1   5.122  0.0017
 eTP4 / stat1     1.0787 0.2382 20    1   0.343  1.0000
 eTP4 / stat2     0.5096 0.1125 20    1  -3.053  0.1289
 exp1 / lTP1     11.4402 2.5262 20    1  11.037  <.0001
 exp1 / lTP2      6.0177 1.3288 20    1   8.128  <.0001
 exp1 / non1     25.7493 5.6859 20    1  14.711  <.0001
 exp1 / stat1     8.9623 1.9790 20    1   9.931  <.0001
 exp1 / stat2     4.2339 0.9349 20    1   6.535  0.0001
 lTP1 / lTP2      0.5260 0.1162 20    1  -2.909  0.1669
 lTP1 / non1      2.2508 0.4970 20    1   3.674  0.0381
 lTP1 / stat1     0.7834 0.1730 20    1  -1.105  0.9783
 lTP1 / stat2     0.3701 0.0817 20    1  -4.501  0.0065
 lTP2 / non1      4.2789 0.9449 20    1   6.583  0.0001
 lTP2 / stat1     1.4893 0.3289 20    1   1.804  0.7259
 lTP2 / stat2     0.7036 0.1554 20    1  -1.592  0.8371
 non1 / stat1     0.3481 0.0769 20    1  -4.779  0.0035
 non1 / stat2     0.1644 0.0363 20    1  -8.175  <.0001
 stat1 / stat2    0.4724 0.1043 20    1  -3.396  0.0669
P value adjustment: tukey method for comparing a family of 10
estimates, Tests are performed on the log scale
```

#### Fig. 2

**Model (Time at max lysis rate)**

```
Call:
lm(formula = tmxdec ~ gphasepd, data = MPmax_lys.1)

Residuals:
    Min      1Q  Median      3Q     Max
-0.8889 -0.3750  0.0000  0.3681  1.3611

Coefficients:
               Estimate Std. Error t value Pr(>|t|)
(Intercept)      8.2500     0.3837  21.501 2.70e-15 ***
gphasepdeTP2     2.6667     0.5426   4.914 8.37e-05 ***
gphasepdeTP3     4.9722     0.5426   9.163 1.35e-08 ***
gphasepdeTP4    11.5278     0.5426  21.244 3.40e-15 ***
gphasepdexp1    -3.6111     0.5426  -6.655 1.77e-06 ***
gphasepdlTP1     5.3889     0.5426   9.931 3.55e-09 ***
gphasepdlTP2    13.1667     0.5426  24.265 2.62e-16 ***
gphasepdnon1    12.0556     0.5426  22.217 1.44e-15 ***
gphasepdstat1   13.1667     0.5426  24.265 2.62e-16 ***
gphasepdstat2   16.7778     0.5426  30.920  < 2e-16 ***
---
Signif. codes:  0 ‘***’ 0.001 ‘**’ 0.01 ‘*’ 0.05 ‘.’ 0.1 ‘ ’ 1

Residual standard error: 0.6646 on 20 degrees of freedom
MulTPiple R-squared:  0.9927, Adjusted R-squared:  0.9894
F-statistic: 302.7 on 9 and 20 DF,  p-value: < 2.2e-16
```

**Analysis of Variance Table**

```
Response: tmxdec
          Df  Sum Sq Mean Sq F value    Pr(>F)
gphasepd   9 1203.20 133.689  302.69 < 2.2e-16 ***
Residuals 20    8.83   0.442
---
Signif. codes:  0 ‘***’ 0.001 ‘**’ 0.01 ‘*’ 0.05 ‘.’ 0.1 ‘ ’ 1
```

**Simultaneous Tests for General Linear Hypotheses**

```
 contrast      estimate    SE df t.ratio p.value
 eTP1 - eTP2     -2.667 0.543 20  -4.914  0.0026
 eTP1 - eTP3     -4.972 0.543 20  -9.163  <.0001
 eTP1 - eTP4    -11.528 0.543 20 -21.244  <.0001
 eTP1 - exp1      3.611 0.543 20   6.655  0.0001
 eTP1 - lTP1     -5.389 0.543 20  -9.931  <.0001
 eTP1 - lTP2    -13.167 0.543 20 -24.265  <.0001
 eTP1 - non1    -12.056 0.543 20 -22.217  <.0001
 eTP1 - stat1   -13.167 0.543 20 -24.265  <.0001
 eTP1 - stat2   -16.778 0.543 20 -30.920  <.0001
 eTP2 - eTP3     -2.306 0.543 20  -4.249  0.0113
 eTP2 - eTP4     -8.861 0.543 20 -16.330  <.0001
 eTP2 - exp1      6.278 0.543 20  11.569  <.0001
 eTP2 - lTP1     -2.722 0.543 20  -5.017  0.0021
 eTP2 - lTP2    -10.500 0.543 20 -19.350  <.0001
 eTP2 - non1     -9.389 0.543 20 -17.303  <.0001
 eTP2 - stat1   -10.500 0.543 20 -19.350  <.0001
 eTP2 - stat2   -14.111 0.543 20 -26.005  <.0001
 eTP3 - eTP4     -6.556 0.543 20 -12.081  <.0001
 eTP3 - exp1      8.583 0.543 20  15.818  <.0001
 eTP3 - lTP1     -0.417 0.543 20  -0.768  0.9984
 eTP3 - lTP2     -8.194 0.543 20 -15.101  <.0001
 eTP3 - non1     -7.083 0.543 20 -13.054  <.0001
 eTP3 - stat1    -8.194 0.543 20 -15.101  <.0001
 eTP3 - stat2   -11.806 0.543 20 -21.756  <.0001
 eTP4 - exp1     15.139 0.543 20  27.899  <.0001
 eTP4 - lTP1      6.139 0.543 20  11.313  <.0001
 eTP4 - lTP2     -1.639 0.543 20  -3.020  0.1368
 eTP4 - non1     -0.528 0.543 20  -0.973  0.9908
 eTP4 - stat1    -1.639 0.543 20  -3.020  0.1368
 eTP4 - stat2    -5.250 0.543 20  -9.675  <.0001
 exp1 - lTP1     -9.000 0.543 20 -16.586  <.0001
 exp1 - lTP2    -16.778 0.543 20 -30.920  <.0001
 exp1 - non1    -15.667 0.543 20 -28.872  <.0001
 exp1 - stat1   -16.778 0.543 20 -30.920  <.0001
 exp1 - stat2   -20.389 0.543 20 -37.574  <.0001
 lTP1 - lTP2     -7.778 0.543 20 -14.334  <.0001
 lTP1 - non1     -6.667 0.543 20 -12.286  <.0001
 lTP1 - stat1    -7.778 0.543 20 -14.334  <.0001
 lTP1 - stat2   -11.389 0.543 20 -20.988  <.0001
 lTP2 - non1      1.111 0.543 20   2.048  0.5792
 lTP2 - stat1     0.000 0.543 20   0.000  1.0000
 lTP2 - stat2    -3.611 0.543 20  -6.655  0.0001
 non1 - stat1    -1.111 0.543 20  -2.048  0.5792
 non1 - stat2    -4.722 0.543 20  -8.703  <.0001
 stat1 - stat2   -3.611 0.543 20  -6.655  0.0001
P value adjustment: tukey method for comparing a family of 10
estimates
```

#### % decrease OD (**Table S3**)

**Model**

```
Call:
glm(formula = percdec ~ gphase, family = quasi(link =
"identity", variance = "mu"))

Deviance Residuals:
     Min        1Q    Median        3Q       Max
-0.24729  -0.12227  -0.00907   0.10248   0.38716

Coefficients:
              Estimate Std. Error t value Pr(>|t|)
(Intercept)    33.3777     0.6689   49.90 2.52e-13 ***
gphaseeTP      45.3371     1.2258   36.99 4.97e-12 ***
gphaseexp      50.2897     1.2525   40.15 2.20e-12 ***
gphaselTP      50.6436     1.2544   40.37 2.08e-12 ***
gphasestat     49.8589     1.2502   39.88 2.35e-12 ***
---
Signif. codes:  0 ‘***’ 0.001 ‘**’ 0.01 ‘*’ 0.05 ‘.’ 0.1 ‘ ’ 1

(Dispersion parameter for quasi family taken to be 0.0402114)
Null deviance: 96.02773  on 14  degrees of freedom
Residual deviance:  0.40037  on 10  degrees of freedom
AIC: NA
Number of Fisher Scoring iterations: 3
```

**Analysis of Deviance Table**

```
Model: quasi, link: identity
Response: percdec
Terms added sequentially (first to last)
       Df Deviance Resid. Df Resid. Dev  Pr(>Chi)
NULL                      14     96.028
gphase  4   95.627        10      0.400 < 2.2e-16 ***
---
Signif. codes:  0 ‘***’ 0.001 ‘**’ 0.01 ‘*’ 0.05 ‘.’ 0.1 ‘ ’ 1
```

**Simultaneous Tests for General Linear Hypotheses**

```
 contrast      estimate   SE  df z.ratio p.value
 non - eTP      -45.337 1.23 Inf -36.987  <.0001
 non - exp      -50.290 1.25 Inf -40.150  <.0001
 non - lTP      -50.644 1.25 Inf -40.372  <.0001
 non - stat     -49.859 1.25 Inf -39.880  <.0001
 eTP - exp       -4.953 1.48 Inf  -3.357  0.0071
 eTP - lTP       -5.306 1.48 Inf  -3.593  0.0030
 eTP - stat      -4.522 1.47 Inf  -3.069  0.0183
 exp - lTP       -0.354 1.50 Inf  -0.236  0.9993
 exp - stat       0.431 1.50 Inf   0.288  0.9985
 lTP - stat       0.785 1.50 Inf   0.524  0.9849

P value adjustment: tukey method for comparing a family of 5
estimates
```

Fig. 2

#### Time to 50% OD loss (**Table S3**)

**Model**

```
Call:
glm(formula = time50 ~ gphase, family = Gamma(link =
"identity"))

Deviance Residuals:
     Min        1Q    Median        3Q       Max
-0.48654  -0.06529  -0.01056   0.01876   0.42775

Coefficients:
             Estimate Std. Error t value Pr(>|t|)
(Intercept)      9.056      1.265   7.158 9.63e-05 ***
gphaseexp       -8.806      1.266  -6.958 0.000117 ***
gphaselTP        2.806      2.085   1.346 0.215259
gphasestat       2.861      2.091   1.368 0.208379
---
Signif. codes:  0 ‘***’ 0.001 ‘**’ 0.01 ‘*’ 0.05 ‘.’ 0.1 ‘ ’ 1

(Dispersion parameter for Gamma family taken to be 0.05854593)
Null deviance: 16.57266  on 11  degrees of freedom
Residual deviance:  0.47674  on  8  degrees of freedom
AIC: 39.559
Number of Fisher Scoring iterations: 3
```

**Analysis of Deviance Table**

```
Model: Gamma, link: identity
Response:  time50
Terms added sequentially (first to last)
      Df Deviance Resid. Df Resid. Dev  Pr(>Chi)
NULL                     11    16.5727
phase  3   16.096         8     0.4767 < 2.2e-16 ***
---
Signif. codes:  0 ‘***’ 0.001 ‘**’ 0.01 ‘*’ 0.05 ‘.’ 0.1 ‘ ’ 1
```

**Simultaneous Tests for General Linear Hypotheses**

```
 contrast     estimate   SE df t.ratio p.value
 eTP - exp      8.8056 1.27  8   6.958  0.0005
 eTP - lTP     -2.8056 2.08  8  -1.346  0.5624
 eTP - stat    -2.8611 2.09  8  -1.368  0.5500
 exp - lTP    -11.6111 1.66  8  -7.006  0.0005
 exp - stat   -11.6667 1.67  8  -7.007  0.0005
 lTP - stat    -0.0556 2.35  8  -0.024  1.0000

P value adjustment: tukey method for comparing a family of 4
estimates
```

------------------------------------------------------------------------------------------------------------------------------------

#### *Fig. 2b*
#### Duration of infection

**Model (also in panel)**

```
Call:
glm(formula = dur ~ gphase, family = Gamma(link = "identity"))

Deviance Residuals:
     Min        1Q    Median        3Q       Max
-0.09014  -0.05038  -0.02226   0.04452   0.13458

Coefficients:
              Estimate Std. Error t value Pr(>|t|)
(Intercept)    18.0833     0.8869  20.390 3.50e-08 ***
gphaseexp     -10.7778     0.9565 -11.268 3.46e-06 ***
gphaselTP       0.3056     1.2649   0.242    0.815
gphasestat     -0.8333     1.2257  -0.680    0.516
---
Signif. codes:  0 ‘***’ 0.001 ‘**’ 0.01 ‘*’ 0.05 ‘.’ 0.1 ‘ ’ 1

(Dispersion parameter for Gamma family taken to be 0.007215803)
Null deviance: 1.597989  on 11  degrees of freedom
Residual deviance: 0.056262  on  8  degrees of freedom
AIC: 43.515
Number of Fisher Scoring iterations: 3
```

**Analysis of Deviance Table**

```
Model: Gamma, link: identity
Response:  dur
Terms added sequentially (first to last)
      Df Deviance Resid. Df Resid. Dev  Pr(>Chi)
NULL                     11    1.59799
phase  3   1.5417         8    0.05626 < 2.2e-16 ***
---
Signif. codes:  0 ‘***’ 0.001 ‘**’ 0.01 ‘*’ 0.05 ‘.’ 0.1 ‘ ’ 1
```

**Simultaneous Tests for General Linear Hypotheses**

```
 contrast     estimate    SE df t.ratio p.value
 eTP - exp      10.778 0.957  8  11.268  <.0001
 eTP - lTP      -0.306 1.265  8  -0.242  0.9946
 eTP - stat      0.833 1.226  8   0.680  0.9019
 exp - lTP     -11.083 0.970  8 -11.421  <.0001
 exp - stat     -9.944 0.919  8 -10.824  <.0001
 lTP - stat      1.139 1.237  8   0.921  0.7949

P value adjustment: tukey method for comparing a family of 4
estimates
```

=================================================================

### Figure 3

#### *Fig. 3a*

##### OD~time*infection status relationship

```
Model 0-10h (panel)
Call:
lm(formula = od ~ time + I(time^2) * infstat)

Residuals:
     Min       1Q   Median       3Q      Max
-0.37771 -0.15942 -0.03794  0.11761  0.50180

Coefficients:
                      Estimate Std. Error t value Pr(>|t|)
(Intercept)           3.164481   0.067449  46.916  < 2e-16 ***
time                 -0.283755   0.027164 -10.446  < 2e-16 ***
I(time^2)             0.015994   0.002709   5.903 7.51e-08 ***
infstatinf           -0.101539   0.067546  -1.503    0.137
I(time^2):infstatinf -0.006633   0.001408  -4.712 9.74e-06 ***
---
Signif. codes:  0 ‘***’ 0.001 ‘**’ 0.01 ‘*’ 0.05 ‘.’ 0.1 ‘ ’ 1

Residual standard error: 0.2168 on 83 degrees of freedom
Multiple R-squared:  0.8708,  Adjusted R-squared:  0.8646
F-statistic: 139.8 on 4 and 83 DF,  p-value: < 2.2e-16

Analysis of Variance Table
Response: od
                  Df  Sum Sq Mean Sq F value    Pr(>F)
time               1 21.6848 21.6848 461.536 < 2.2e-16 ***
I(time^2)          1  1.1032  1.1032  23.481 5.792e-06 ***
infstat            1  2.4496  2.4496  52.136 2.264e-10 ***
I(time^2):infstat  1  1.0433  1.0433  22.206 9.741e-06 ***
Residuals         83  3.8997  0.0470
---
Signif. codes:  0 ‘***’ 0.001 ‘**’ 0.01 ‘*’ 0.05 ‘.’ 0.1 ‘ ’ 1
```

```
Model 26-30h (panel)
Call:
lm(formula = od ~ time * infstat)

Residuals:
    Min      1Q  Median      3Q     Max
-0.2917 -0.1533 -0.1118  0.1535  0.4603

Coefficients:
                Estimate Std. Error t value Pr(>|t|)
(Intercept)      1.02217    1.20068   0.851    0.405
time             0.02025    0.04281   0.473    0.641
infstatinf      -0.57350    1.69802  -0.338    0.739
time:infstatinf -0.02031    0.06054  -0.336    0.741

Residual standard error: 0.2422 on 20 degrees of freedom
Multiple R-squared:  0.8699,  Adjusted R-squared:  0.8504
F-statistic: 44.57 on 3 and 20 DF,  p-value: 4.83e-09

Analysis of Variance Table
Response: od
             Df Sum Sq Mean Sq  F value    Pr(>F)
time          1 0.0065  0.0065   0.1112    0.7423
infstat       1 7.8284  7.8284 133.4931 2.647e-10 ***
time:infstat  1 0.0066  0.0066   0.1126    0.7407
Residuals    20 1.1729  0.0586
---
Signif. codes:  0 ‘***’ 0.001 ‘**’ 0.01 ‘*’ 0.05 ‘.’ 0.1 ‘ ’ 1
```

##### Maximum decline rates & times at max decline rates (**Table S3**)

```
Model (Max decline rate, comparison different periods)
Call:
lm(formula = mxdecrt ~ infstat + tpd)

Residuals:
      Min        1Q    Median        3Q       Max
-0.037320 -0.016260 -0.005387  0.008332  0.051994

Coefficients:
             Estimate Std. Error t value Pr(>|t|)
(Intercept)  -0.12271    0.01194 -10.273 1.32e-07 ***
infstatnon    0.02279    0.01379   1.653    0.122
tpd26-30      0.11970    0.01379   8.679 9.07e-07 ***
---
Signif. codes:  0 ‘***’ 0.001 ‘**’ 0.01 ‘*’ 0.05 ‘.’ 0.1 ‘ ’ 1

Residual standard error: 0.02758 on 13 degrees of freedom
Multiple R-squared:  0.8572,  Adjusted R-squared:  0.8353
F-statistic: 39.03 on 2 and 13 DF,  p-value: 0.0000032

Analysis of Variance Table
Response:  mxdecrt
            Df   Sum Sq  Mean Sq F value    Pr(>F)
infstat      1 0.002078 0.002078   2.731    0.1224
tpd          1 0.057314 0.057314  75.322 9.068e-07 ***
Residuals   13 0.009892 0.000761
---
Signif. codes:  0 ‘***’ 0.001 ‘**’ 0.01 ‘*’ 0.05 ‘.’ 0.1 ‘ ’ 1

Simultaneous Tests for General Linear Hypotheses
Multiple Comparisons of Means: Tukey Contrasts
Fit: lm(formula = mxdecrt ~ infstat + tpd)
Linear Hypotheses:
contrast       Estimate Std. Error t value Pr(>|t|)
non - inf == 0  0.02279    0.01379   1.653    0.122
(Adjusted p values reported -- single-step method)

Linear Hypotheses:
 contrast          Estimate Std. Error t value Pr(>|t|)
26-30 - 0-10 == 0  0.11970    0.01379   8.679 9.07e-07 ***
---
Signif. codes:  0 ‘***’ 0.001 ‘**’ 0.01 ‘*’ 0.05 ‘.’ 0.1 ‘ ’ 1
(Adjusted p values reported -- single-step method)
```

```
Model (Time at max decline rate, comparison different periods)
Call:
lm(formula = tmxdec ~ tpd + infstat)

Residuals:
    Min      1Q  Median      3Q     Max
-3.4375 -1.5313 -0.8125  0.8125  4.5625

Coefficients:
            Estimate Std. Error t value Pr(>|t|)
(Intercept)   4.4375     0.9853   4.504 0.000593 ***
tpd26-30     24.3750     1.1377  21.424 1.59e-11 ***
infstatnon   -1.6250     1.1377  -1.428 0.176800
---
Signif. codes:  0 ‘***’ 0.001 ‘**’ 0.01 ‘*’ 0.05 ‘.’ 0.1 ‘ ’ 1

Residual standard error: 2.275 on 13 degrees of freedom
Multiple R-squared:  0.9726,  Adjusted R-squared:  0.9684
F-statistic: 230.5 on 2 and 13 DF,  p-value: 7.046e-11

Analysis of Variance Table
Response:  tmxdec
            Df  Sum Sq Mean Sq  F value    Pr(>F)
tpd          1 2376.56 2376.56 458.9833 1.592e-11 ***
infstat      1   10.56   10.56   2.0399    0.1768
Residuals   13   67.31    5.18
---
Signif. codes:  0 ‘***’ 0.001 ‘**’ 0.01 ‘*’ 0.05 ‘.’ 0.1 ‘ ’ 1

Simultaneous Tests for General Linear Hypotheses
Multiple Comparisons of Means: Tukey Contrasts
Fit: lm(formula = tmxdec ~ tpd + infstat)

Linear Hypotheses:
contrast          Estimate Std. Error t value Pr(>|t|)
26-30 - 0-10 == 0   24.375      1.138   21.42 1.59e-11 ***
---
Signif. codes:  0 ‘***’ 0.001 ‘**’ 0.01 ‘*’ 0.05 ‘.’ 0.1 ‘ ’ 1
(Adjusted p values reported -- single-step method)
```

##### Time to lysis initiation (**Table S3**)

Not determined – rate already negative from start

### Fig. 3

#### % decrease OD (**Table S3**)

**Model (Comparison different periods)**
**NOTE:** analysed as proportion data rather than percentage data

```
Call:
lm(formula = propdec ~ tpd * infstat)

Residuals:
     Min       1Q   Median       3Q      Max 
-0.12177 -0.03390  0.01401  0.03508  0.10320 

Coefficients:
                      Estimate Std. Error t value Pr(>|t|)    
(Intercept)            0.62389    0.03580  17.426 6.92e-10 ***
tpd26-30               0.23674    0.05063   4.676 0.000536 ***
infstatnon            -0.22122    0.05063  -4.369 0.000913 ***
tpd26-30:infstatnon   -0.16470    0.07160  -2.300 0.040187 *  
---
Signif. codes:  0 '***' 0.001 '**' 0.01 '*' 0.05 '.' 0.1 ' ' 1

Residual standard error: 0.0716 on 12 degrees of freedom
Multiple R-squared:  0.8887,  Adjusted R-squared:  0.8608 
F-statistic: 31.93 on 3 and 12 DF,  p-value: 5.31e-06
```

**Analysis of Variance Table**

```
Response:  propdec
               Df  Sum Sq Mean Sq F value    Pr(>F)    
tpd             1 0.09535 0.09535 18.5981  0.001009 ** 
infstat         1 0.36862 0.36862 71.8990 2.062e-06 ***
tpd:infstat     1 0.02712 0.02712  5.2907  0.040187 *  
Residuals      12 0.06152 0.00513                      
---
Signif. codes:  0 '***' 0.001 '**' 0.01 '*' 0.05 '.' 0.1 ' ' 1
```

**Simultaneous Tests for General Linear Hypotheses**

```
infstat = inf:
 contrast     estimate     SE df t.ratio p.value
 0-10 - 26-30   -0.237 0.0506 12  -4.676  0.0005

infstat = non:
 contrast     estimate     SE df t.ratio p.value
 0-10 - 26-30   -0.072 0.0506 12  -1.423  0.1802


tpd = 0-10:
 contrast  estimate     SE df t.ratio p.value
 inf - non    0.221 0.0506 12   4.369  0.0009

tpd = 26-30:
 contrast  estimate     SE df t.ratio p.value
 inf - non    0.386 0.0506 12   7.622  <.0001
```

---

#### *Fig. 3b*

#### CFU~time*infection status relationship

| time p.i. (h) | infection status | mean CFU $ml^{-1}$ | SD CFU $ml^{-1}$ |
|---|---|---|---|
| 0 | non-infected | 3.95E+08 | 1.11E+08 |
| 0 | infected | 4.28E+08 | 9.57E+07 |
| 10 | non-infected | 3.64E+08 | 1.18E+08 |
| 10 | infected | 9.55E+07 | 1.87E+07 |
| 30 | non-infected | 3.42E+08 | 3.18E+08 |
| 30 | infected | 5.17E+06 | 7.38E+06 |

**Model (panel)**

```
Call:
glm(formula = CFU ~ time * infstat, family = quasipoisson(link = 
"identity"))

Deviance Residuals: 
   Min      1Q  Median      3Q     Max  
-15242   -6824   -1913    4420   21617  

Coefficients:
                  Estimate Std. Error t value Pr(>|t|)    
(Intercept)      389033589   75094899   5.181 4.54e-05 ***
time              -1646269    3980387  -0.414   0.6836    
infstatinf       -75631272   97611488  -0.775   0.4475    
time:infstatinf   -8654592    4506009  -1.921   0.0691 .  
---
Signif. codes:  0 '***' 0.001 '**' 0.01 '*' 0.05 '.' 0.1 ' ' 1

(Dispersion parameter for quasipoisson family taken to be 
82415651)
Null deviance: 4344000838  on 23  degrees of freedom
Residual deviance: 1550776151  on 20  degrees of freedom
AIC: NA
Number of Fisher Scoring iterations: 6
```

**Analysis of Deviance Table**

```
Model: quasipoisson, link: identity
Response:  CFU
Terms added sequentially (first to last)
               Df   Deviance Resid. Df Resid. Dev  Pr(>Chi)    
NULL                                23 4344000838              
time            1  690011186        22 3653989652   0.00381 ** 
infstat         1 1780193411        21 1873796241 3.358e-06 ***
time:infstat    1  323020089        20 1550776151   0.04773 *  
---
Signif. codes:  0 '***' 0.001 '**' 0.01 '*' 0.05 '.' 0.1 ' ' 1
```

### Fig. 3

#### % spore~time*infection status relationship

**Model (panel)**

```
Call:
lm(formula = propspr ~ time + infstat)

Residuals:
      Min        1Q    Median        3Q       Max
-0.020918 -0.006176 -0.004416  0.003298  0.054051

Coefficients:
                Estimate Std. Error t value Pr(>|t|)
(Intercept)   -0.0033718  0.0062609  -0.539   0.5959
time           0.0009420  0.0002832   3.327   0.0032 **
infstatinf     0.0002274  0.0070632   0.032   0.9746
---
Signif. codes:  0 ‘***’ 0.001 ‘**’ 0.01 ‘*’ 0.05 ‘.’ 0.1 ‘ ’ 1

Residual standard error: 0.0173 on 21 degrees of freedom
Multiple R-squared:  0.3452,  Adjusted R-squared:  0.2828
F-statistic: 5.534 on 2 and 21 DF,  p-value: 0.01173
```

**Analysis of Variance Table**

```
Response:  propspr
          Df    Sum Sq   Mean Sq F value   Pr(>F)
time       1 0.0033129 0.0033129  11.068 0.003203 **
infstat    1 0.0000003 0.0000003   0.001 0.974617
Residuals 21 0.0062861 0.0002993
---
Signif. codes:  0 ‘***’ 0.001 ‘**’ 0.01 ‘*’ 0.05 ‘.’ 0.1 ‘ ’ 1
```

---

***Fig. 3c***

#### PFU~time*population relationship

**Model 0-10h (panel)**

```
Call:
glm(formula = PFU ~ time + I(time^2) * ppn, family = Gamma(link
= "log"))

Deviance Residuals:
     Min        1Q    Median        3Q       Max
-1.31010  -0.16310  -0.00333   0.15945   0.50817

Coefficients:
                      Estimate Std. Error t value Pr(>|t|)
(Intercept)          20.071137   0.086956 230.820  < 2e-16 ***
time                  0.025415   0.038479   0.660 0.511916
I(time^2)             0.013856   0.003908   3.546 0.000849 ***
ppntotal              1.155159   0.094757  12.191  < 2e-16 ***
I(time^2):ppntotal   -0.005213   0.002003  -2.603 0.012086 *
---
Signif. codes:  0 ‘***’ 0.001 ‘**’ 0.01 ‘*’ 0.05 ‘.’ 0.1 ‘ ’ 1

(Dispersion parameter for Gamma family taken to be 0.06971564)
Null deviance: 29.811  on 55  degrees of freedom
Residual deviance:  4.607  on 51  degrees of freedom
AIC: 2392.2
Number of Fisher Scoring iterations: 5
```

**Analysis of Deviance Table**

```
Model: Gamma, link: log
Response: PFU
Terms added sequentially (first to last)
              Df Deviance Resid. Df Resid. Dev  Pr(>Chi)
NULL                             55    29.8113
time           1  10.8261        54    18.9853 < 2.2e-16 ***
I(time^2)      1   0.8576        53    18.1277 0.0004528 ***
ppn            1  13.0651        52     5.0626 < 2.2e-16 ***
I(time^2):ppn  1   0.4555        51     4.6070 0.0105823 *
---
Signif. codes:  0 ‘***’ 0.001 ‘**’ 0.01 ‘*’ 0.05 ‘.’ 0.1 ‘ ’ 1
```

**Test 30h (panel)**
**Wilcoxon rank sum test**

```
W = 6, p-value = 0.6857
alternative hypothesis: true location shift is not equal to 0
```

#### Maximum PFU rate & time at max PFU rate (**Table S3**)

**Max PFU rate**
**Wilcoxon rank sum test (Table S3)**

```
W = 14, p-value = 0.1143
alternative hypothesis: true location shift is not equal to 0
```

**Time at maximum PFU rate**
**Welch Two Sample t-test (Table S3)**

```
t = -1.2649, df = 5.6017, p-value = 0.256
alternative hypothesis: true difference in means is not equal to
0
95 percent confidence interval:
 -5.936587  1.936587
sample estimates:
mean of x mean of y
      4.5       6.5
```

Fig. 3

#### Fold change in PFU (**Table S3**)

```
Model 0-10h vs 10-30h
Call:
glm(formula = fch ~ time.fac * ppn, family = Gamma(link = log))

Deviance Residuals:
     Min        1Q    Median        3Q       Max
-0.57750  -0.23325   0.02226   0.20295   0.41372

Coefficients:
                     stimate Std. Error t value Pr(>|t|)
(Intercept)           0.1527     0.1527   1.000 0.336999
time.fac30            1.0298     0.2160   4.768 0.000458 ***
ppntotal              0.7686     0.2160   3.559 0.003932 **
time.fac30:ppntotal  -0.7273     0.3054  -2.381 0.034680 *
---
Signif. codes:  0 ‘***’ 0.001 ‘**’ 0.01 ‘*’ 0.05 ‘.’ 0.1 ‘ ’ 1

(Dispersion parameter for Gamma family taken to be 0.09327464)
Null deviance: 3.7553  on 15  degrees of freedom
Residual deviance: 1.2063  on 12  degrees of freedom
AIC: 40.882
Number of Fisher Scoring iterations: 4


Analysis of Deviance Table
Model: Gamma, link: log
Response: fch
Terms added sequentially (first to last)
             Df Deviance Resid. Df Resid. Dev  Pr(>Chi)
NULL                            15     3.7553
time.fac      1  1.39228        14     2.3631 0.0001118 ***
ppn           1  0.63075        13     1.7323 0.0093105 **
time.fac:ppn  1  0.52604        12     1.2063 0.0175582 *
---
Signif. codes:  0 ‘***’ 0.001 ‘**’ 0.01 ‘*’ 0.05 ‘.’ 0.1 ‘ ’ 1
05 ‘.’ 0.1 ‘ ’ 1
```

```
Simultaneous Tests for General Linear Hypotheses
ppn = free:
contrast ratio     SE df null t.ratio p.value
 10 / 30 0.357 0.0771 12    1  -4.768  0.0005

ppn = total:
contrast ratio     SE df null t.ratio p.value
 10 / 30 0.739 0.1596 12    1  -1.401  0.1866

Tests are performed on the log scale

------

time.fac = 10:
 contrast     ratio    SE df null t.ratio p.value
 free / total 0.464 0.100 12    1  -3.559  0.0039

time.fac = 30:
 contrast     ratio    SE df null t.ratio p.value
 free / total 0.960 0.207 12    1  -0.191  0.8516

Tests are performed on the log scale
```

#### Average burst sizes (**Table S3**)

```
Wilcoxon rank sum test (Comparison different time periods; 0-10h, 10-30h)
W = 0, p-value = 0.02857
alternative hypothesis: true location shift is not equal to 0
```

---

#### *Fig. 3e*
#### % infected cells~time relationship

```
Model (panel)
NOTE: ‘response’ is coded here as a matrix of the number of
infected and uninfected cells at each time point

Call:
glm(formula = response ~ time + I(time^2), family =
quasibinomial(link = "logit"))

Deviance Residuals:
    Min       1Q   Median       3Q      Max
-3.6804  -1.7051  -0.7802   1.3225   3.8155

Coefficients:
            Estimate Std. Error t value Pr(>|t|)
(Intercept) -4.75182    0.39525 -12.022 6.90e-12 ***
time         0.99601    0.13034   7.642 5.36e-08 ***
I(time^2)   -0.05748    0.01011  -5.683 6.44e-06 ***
---
Signif. codes:  0 ‘***’ 0.001 ‘**’ 0.01 ‘*’ 0.05 ‘.’ 0.1 ‘ ’ 1

(Dispersion parameter for quasibinomial family taken to be
3.722048)
Null deviance: 964.51  on 27  degrees of freedom
Residual deviance: 106.90  on 25  degrees of freedom
AIC: NA
Number of Fisher Scoring iterations: 5
```

```
Analysis of Deviance Table
Model: quasibinomial, link: logit
Response: response
Terms added sequentially (first to last)

          Df Deviance Resid. Df Resid. Dev  Pr(>Chi)
NULL                         27     964.51
time       1   717.60        26     246.92 < 2.2e-16 ***
I(time^2)  1   140.01        25     106.90 8.609e-10 ***
---
Signif. codes:  0 ‘***’ 0.001 ‘**’ 0.01 ‘*’ 0.05 ‘.’ 0.1 ‘
```

=================================================================

### Fig. 4

#### Figure 4

##### *Fig. 4a*

##### Recovered PFU~growth phase*strain relationship

**Model (panel)**

```
Call:
lm(formula = log(PFU) ~ strain * time.fac)

Residuals:
     Min       1Q   Median       3Q      Max 
-1.57873 -0.15705 -0.01681  0.21582  1.36571 

Coefficients:
                      Estimate Std. Error t value Pr(>|t|)    
(Intercept)            11.2289     0.3326  33.761  < 2e-16 ***
strainpsyB             -4.7753     0.4704 -10.152 3.66e-10 ***
strainMO                4.1497     0.4704   8.822 5.35e-09 ***
time.fac4               2.8728     0.4704   6.107 2.62e-06 ***
time.fac6               2.9481     0.4704   6.268 1.77e-06 ***
time.fac10              2.7283     0.4704   5.800 5.58e-06 ***
strainpsyB:time.fac4   -3.1997     0.6652  -4.810 6.73e-05 ***
strainMO:time.fac4     -3.0881     0.6652  -4.642 0.000103 ***
strainpsyB:time.fac6   -3.2750     0.6652  -4.923 5.05e-05 ***
strainMO:time.fac6     -3.1055     0.6652  -4.669 9.65e-05 ***
strainpsyB:time.fac10  -2.9979     0.6652  -4.507 0.000146 ***
strainMO:time.fac10    -2.9895     0.6652  -4.494 0.000150 ***
---
Signif. codes:  0 ‘***’ 0.001 ‘**’ 0.01 ‘*’ 0.05 ‘.’ 0.1 ‘ ’ 1

Residual standard error: 0.5761 on 24 degrees of freedom
Multiple R-squared:  0.986,	Adjusted R-squared:  0.9796 
F-statistic: 153.5 on 11 and 24 DF,  p-value: < 2.2e-16
```

**Analysis of Variance Table**

```
Response: log(PFU)
                Df Sum Sq Mean Sq  F value    Pr(>F)    
strain           2 541.68 270.839 816.1015 < 2.2e-16 ***
time.fac         3   4.11   1.370   4.1272 0.0170847 *  
strain:time.fac  6  14.57   2.429   7.3193 0.0001569 ***
Residuals       24   7.96   0.332                       
---
Signif. codes:  0 ‘***’ 0.001 ‘**’ 0.01 ‘*’ 0.05 ‘.’ 0.1 ‘ ’ 1
```

**Simultaneous Tests for General Linear Hypotheses**

```
time.fac = 2.25:
 contrast        ratio        SE df null t.ratio p.value
 WT / psyB    118.5473   55.7608 24    1  10.152  <.0001
 WT / MO        0.0158    0.0074 24    1  -8.822  <.0001
 psyB / MO      0.0001    0.0001 24    1 -18.974  <.0001

time.fac = 4:
 contrast        ratio        SE df null t.ratio p.value
 WT / psyB   2907.4151 1367.5543 24    1  16.955  <.0001
 WT / MO        0.3459    0.1627 24    1  -2.257  0.0818
 psyB / MO      0.0001    0.0001 24    1 -19.212  <.0001

time.fac = 6:
 contrast        ratio        SE df null t.ratio p.value
 WT / psyB   3134.8825 1474.5476 24    1  17.115  <.0001
 WT / MO        0.3520    0.1656 24    1  -2.220  0.0880
 psyB / MO      0.0001    0.0001 24    1 -19.335  <.0001

time.fac = 10:
 contrast        ratio        SE df null t.ratio p.value
 WT / psyB   2376.1921 1117.6841 24    1  16.526  <.0001
 WT / MO        0.3134    0.1474 24    1  -2.467  0.0533
 psyB / MO      0.0001    0.0001 24    1 -18.993  <.0001

P value adjustment: tukey method for comparing a family of 3 estimates 
Tests are performed on the log scale 

------

strain = WT:
 contrast   ratio     SE df null t.ratio p.value
 2.25 / 4  0.0565 0.0266 24    1  -6.107  <.0001
 2.25 / 6  0.0524 0.0247 24    1  -6.268  <.0001
 2.25 / 10 0.0653 0.0307 24    1  -5.800  <.0001
 4 / 6     0.9274 0.4362 24    1  -0.160  0.9985
 4 / 10    1.1554 0.5435 24    1   0.307  0.9897
 6 / 10    1.2458 0.5860 24    1   0.467  0.9655

strain = psyB:
 contrast   ratio     SE df null t.ratio p.value
 2.25 / 4  1.3867 0.6523 24    1   0.695  0.8980
 2.25 / 6  1.3867 0.6523 24    1   0.695  0.8980
 2.25 / 10 1.3095 0.6160 24    1   0.573  0.9391
 4 / 6     1.0000 0.4704 24    1   0.000  1.0000
 4 / 10    0.9443 0.4442 24    1  -0.122  0.9993
 6 / 10    0.9443 0.4442 24    1  -0.122  0.9993

strain = MO:
 contrast   ratio     SE df null t.ratio p.value
 2.25 / 4  1.2403 0.5834 24    1   0.458  0.9674
 2.25 / 6  1.1705 0.5506 24    1   0.335  0.9868
 2.25 / 10 1.2984 0.6107 24    1   0.555  0.9442
 4 / 6     0.9437 0.4439 24    1  -0.123  0.9993
 4 / 10    1.0469 0.4924 24    1   0.097  0.9997
 6 / 10    1.1093 0.5218 24    1   0.221  0.9961

P value adjustment: tukey method for comparing a family of 4 estimates 
Tests are performed on the log scale
```

---

##### *Fig. 4b*

##### Maximum growth rate before infection (**Table S3**)

**Welch Two Sample t-test**

```
t = 3.1476, df = 2.4787, p-value = 0.06641
alternative hypothesis: true difference in means is not equal to 0
95 percent confidence interval:
 -0.005632195  0.084578128
sample estimates:
mean of x mean of y 
0.8208252 0.7813523 
```

##### Time at maximum growth rate before infection (**Table S3**)

**Welch Two Sample t-test**

```
t = -2.1213, df = 4, p-value = 0.1012
alternative hypothesis: true difference in means is not equal to 0
95 percent confidence interval:
 -0.38480480  0.05147146
sample estimates:
mean of x mean of y 
 1.944444  2.111111 
```

#### Fig. 4

### Growth/decline rates at time of infection (**Table S3**)

**Model**

```
Call:
lm(formula = (gdecrt * -1) ~ strain + infstat, family =
inverse.gaussian(link = "sqrt"))

Residuals:
       Min         1Q     Median         3Q        Max
-0.0110164 -0.0018789 -0.0005988  0.0023004  0.0101516

Coefficients:
              Estimate Std. Error t value Pr(>|t|)
(Intercept)  0.0725929  0.0027917  26.003 8.87e-10 ***
strainpsyB  -0.0215458  0.0032235  -6.684 9.02e-05 ***
infstatnon  -0.0005403  0.0032235  -0.168    0.871
---
Signif. codes:  0 ‘***’ 0.001 ‘**’ 0.01 ‘*’ 0.05 ‘.’ 0.1 ‘ ’ 1

Residual standard error: 0.005583 on 9 degrees of freedom
Multiple R-squared:  0.8324,  Adjusted R-squared:  0.7952
F-statistic: 22.35 on 2 and 9 DF,  p-value: 0.0003229
```

**Analysis of Variance Table**

```
Response: (gdecrt * -1)
          Df     Sum Sq    Mean Sq F value    Pr(>F)
strain     1 0.00139266 0.00139266 44.6744 9.018e-05 ***
infstat    1 0.00000088 0.00000088  0.0281    0.8706
Residuals  9 0.00028056 0.00003117
---
Signif. codes:  0 ‘***’ 0.001 ‘**’ 0.01 ‘*’ 0.05 ‘.’ 0.1 ‘ ’ 1
```

**Simultaneous Tests for General Linear Hypotheses**

```
infstat = inf:
 contrast    estimate      SE df t.ratio p.value
 WT - psyB     0.0215 0.00322  9   6.684  0.0001

infstat = non:
 contrast    estimate      SE df t.ratio p.value
 WT - psyB     0.0215 0.00322  9   6.684  0.0001

------

strain = WT:
 contrast    estimate      SE df t.ratio p.value
 inf - non    0.00054 0.00322  9   0.168  0.8706

strain = psyB:
 contrast    estimate      SE df t.ratio p.value
 inf - non    0.00054 0.00322  9   0.168  0.8706
```

Fig. 4

#### Time to lysis (**Table S3**)

Not determined – rate already negative from start

#### Maximum decline rates & times at max decline rates (**Table S3**)

- break point 1 for all samples was chosen to avoid artefacts in OD traces immediately following infection which could artificially inflate decline rates
- second break-points for non-infected samples set to avoid artefacts introduced due to condensation and clumping at late times during stationary phase
- **only** second period decline rates were analysed for non-infected samples

**Breakpoints for decline rate calculation**

| infection status | bacterial strain | infection time | bp1(h) | bp2(h) | bp3(h) | replicate |
|---|---|---|---|---|---|---|
| non-infected | wt | 11.0 | 13.2 | 33.2 | 40.0 | 1 |
| non-infected | wt | 11.0 | 12.5 | 34.3 | 40.0 | 2 |
| non-infected | wt | 11.0 | 12.0 | 30.3 | 40.0 | 3 |
| non-infected | pspac::yueB | 11.0 | 19.9 | 35.3 | 40.0 | 1 |
| non-infected | pspac::yueB | 11.0 | 23.5 | 35.8 | 40.0 | 2 |
| non-infected | pspac::yueB | 11.0 | 13.9 | 37.9 | 40.0 | 3 |
| infected | wt | 11.0 | 11.7 | 19.9 | 40.0 | 1 |
| infected | wt | 11.0 | 11.7 | 16.3 | 40.0 | 2 |
| infected | wt | 11.0 | 11.8 | 19.9 | 40.0 | 3 |
| infected | pspac::yueB | 11.0 | 12.3 | 15.8 | 40.0 | 1 |
| infected | pspac::yueB | 11.0 | 12.8 | 15.5 | 40.0 | 2 |
| infected | pspac::yueB | 11.0 | 13.2 | 16.9 | 40.0 | 3 |

* periods noted in Table S3 (1-2) correspond to time periods between the above listed breakpoints, starting from breakpoint

**Model max decline rate**
**NOTE:** data were box-cox transformed before analysis and infection status, bacterial strain and period were combined into a single variable for direct comparison between rates

```
Call:
lm(formula = (((mxdecrt * -1)^0.8282828 - 1)/0.8282828) ~ ISSP)

Residuals:
      Min        1Q    Median        3Q       Max
-0.071156 -0.007021 -0.000021  0.004868  0.101820

Coefficients:
                 Estimate Std. Error t value Pr(>|t|)
(Intercept)      -1.05393    0.02362 -44.612 1.05e-14 ***
ISSPnonpsyB1     -0.07562    0.03341  -2.263   0.0429 *
ISSPinfWT1       -0.01899    0.03341  -0.568   0.5802
ISSPinfWT2        0.32320    0.03341   9.674 5.12e-07 ***
ISSPinfpsyB1      0.08079    0.03341   2.418   0.0324 *
ISSPinfpsyB2      0.48029    0.03341  14.376 6.31e-09 ***
---
Signif. codes:  0 ‘***’ 0.001 ‘**’ 0.01 ‘*’ 0.05 ‘.’ 0.1 ‘ ’ 1

Residual standard error: 0.04092 on 12 degrees of freedom
Multiple R-squared:  0.9733,  Adjusted R-squared:  0.9621
F-statistic: 87.37 on 5 and 12 DF,  p-value: 5.171e-09
```

**Analysis of Variance Table**

```
Response: (((mxdecrt * -1)^ 0.8282828 – 1)/0.8282828)
          Df  Sum Sq  Mean Sq F value    Pr(>F)
infstat    5 0.73145 0.146290  87.372 5.171e-09 ***
Residuals 12 0.02009 0.001674
```

**Simultaneous Tests for General Linear Hypotheses**

```
 contrast              estimate     SE df t.ratio p.value
 nonWT1 - nonpsyB1       0.0756 0.0334 12   2.263  0.2793
 nonWT1 - infWT1         0.0190 0.0334 12   0.568  0.9914
 nonWT1 - infWT2        -0.3232 0.0334 12  -9.674  <.0001
 nonWT1 - infpsyB1      -0.0808 0.0334 12  -2.418  0.2240
 nonWT1 - infpsyB2      -0.4803 0.0334 12 -14.376  <.0001
 nonpsyB1 - infWT1      -0.0566 0.0334 12  -1.695  0.5590
 nonpsyB1 - infWT2      -0.3988 0.0334 12 -11.937  <.0001
 nonpsyB1 - infpsyB1    -0.1564 0.0334 12  -4.682  0.0054
 nonpsyB1 - infpsyB2    -0.5559 0.0334 12 -16.639  <.0001
 infWT1 - infWT2        -0.3422 0.0334 12 -10.242  <.0001
 infWT1 - infpsyB1      -0.0998 0.0334 12  -2.987  0.0925
 infWT1 - infpsyB2      -0.4993 0.0334 12 -14.944  <.0001
 infWT2 - infpsyB1       0.2424 0.0334 12   7.256  0.0001
 infWT2 - infpsyB2      -0.1571 0.0334 12  -4.702  0.0053
 infpsyB1 - infpsyB2    -0.3995 0.0334 12 -11.958  <.0001

P value adjustment: tukey method for comparing a family of 6
estimates
```

#### Fig. 4

**Model time at max decline rate**

```
Call:
glm(formula = tmxdec ~ ISSP, family = inverse.gaussian(link =
"inverse"))

Deviance Residuals:
      Min         1Q     Median         3Q        Max
-0.065944   0.000000   0.000000   0.001179   0.036571

Coefficients:
                  Estimate Std. Error t value Pr(>|t|)
(Intercept)      0.0517241  0.0034316  15.073 3.68e-09 ***
ISSPnonpsyB1    -0.0103448  0.0046040  -2.247   0.0442 *
ISSPinfWT1       0.0069078  0.0050124   1.378   0.1933
ISSPinfWT2      -0.0135076  0.0045251  -2.985   0.0114 *
ISSPinfpsyB1     0.0105596  0.0050947   2.073   0.0604 .
ISSPinfpsyB2     0.0009074  0.0048742   0.186   0.8554
---
Signif. codes:  0 ‘***’ 0.001 ‘**’ 0.01 ‘*’ 0.05 ‘.’ 0.1 ‘ ’ 1

(Dispersion parameter for inverse.gaussian family taken to be
0.0006829986)

    Null deviance: 0.037066  on 17  degrees of freedom
Residual deviance: 0.009326  on 12  degrees of freedom
AIC: 90.337

Number of Fisher Scoring iterations: 2
```

**Analysis of Deviance Table**

```
Model: inverse.gaussian, link: inverse
Response: tmxdec
Terms added sequentially (first to last)

     Df Deviance Resid. Df Resid. Dev  Pr(>Chi)
NULL                    17   0.037066
ISSP   5  0.02774        12   0.009326 1.122e-07 ***
---
Signif. codes:  0 ‘***’ 0.001 ‘**’ 0.01 ‘*’ 0.05 ‘.’ 0.1 ‘ ’ 1
```

**Simultaneous Tests for General Linear Hypotheses**

```
 contrast            estimate   SE  df z.ratio p.value
 nonWT1 - nonpsyB1     -4.833 2.20 Inf  -2.193  0.2410
 nonWT1 - infWT1        2.278 1.67 Inf   1.367  0.7466
 nonWT1 - infWT2       -6.833 2.39 Inf  -2.856  0.0491
 nonWT1 - infpsyB1      3.278 1.61 Inf   2.038  0.3208
 nonWT1 - infpsyB2      0.333 1.79 Inf   0.186  1.0000
 nonpsyB1 - infWT1      7.111 2.08 Inf   3.412  0.0085
 nonpsyB1 - infWT2     -2.000 2.70 Inf  -0.741  0.9769
 nonpsyB1 - infpsyB1    8.111 2.04 Inf   3.979  0.0010
 nonpsyB1 – infpsyB2    5.167 2.19 Inf   2.364  0.1689
 infWT1 - infWT2       -9.111 2.28 Inf  -3.992  0.0009
 infWT1 - infpsyB1      1.000 1.44 Inf   0.695  0.9826
 infWT1 - infpsyB2     -1.944 1.64 Inf  -1.185  0.8441
 infWT2 - infpsyB1     10.111 2.24 Inf   4.512  0.0001
 infWT2 - infpsyB2      7.167 2.37 Inf   3.018  0.0306
 infpsyB1 - infpsyB2   -2.944 1.58 Inf  -1.861  0.4266

P value adjustment: tukey method for comparing a family of 6
estimates
```

#### Duration of infection (**Table S3**)

**Wilcoxon rank sum test with continuity correction**

```
W = 9, p-value = 0.04685
alternative hypothesis: true location shift is not equal to 0
```

#### % decrease OD (**Table S3**)

**Model**

**NOTE:** data were analysed here as proportion data to allow arcsine transformation

```
Call:
lm(formula = asin(propdec) ~ infstat + strain)

Residuals:
      Min        1Q    Median        3Q       Max
-0.044260 -0.009163 -0.001441  0.015446  0.035928

Coefficients:
              Estimate Std. Error t value Pr(>|t|)
(Intercept)   0.459268   0.013007  35.308 5.80e-11 ***
infstatinf    0.635188   0.015020  42.291 1.15e-11 ***
strainpsyB   -0.009085   0.015020  -0.605     0.56
---
Signif. codes:  0 ‘***’ 0.001 ‘**’ 0.01 ‘*’ 0.05 ‘.’ 0.1 ‘ ’ 1

Residual standard error: 0.02601 on 9 degrees of freedom
Multiple R-squared:  0.995,   Adjusted R-squared:  0.9939
F-statistic: 894.4 on 2 and 9 DF,  p-value: 4.443e-11
```

**Analysis of Variance Table**

```
Response: asin(propdec)
          Df  Sum Sq Mean Sq   F value    Pr(>F)
infstat    1 1.21039 1.21039 1788.4980 1.153e-11 ***
strain     1 0.00025 0.00025    0.3659    0.5602
Residuals  9 0.00609 0.00068
---
Signif. codes:  0 ‘***’ 0.001 ‘**’ 0.01 ‘*’ 0.05 ‘.’ 0.1 ‘ ’ 1
```

**Simultaneous Tests for General Linear Hypotheses**

```
strain = WT:
 contrast  estimate     SE df t.ratio p.value
 non - inf   -0.635 0.0149  9 -42.706  <.0001

strain = psyB:
 contrast  estimate     SE df t.ratio p.value
 non - inf   -0.635 0.0149  9 -42.706  <.0001

------

infstat = non:
 contrast estimate     SE df t.ratio p.value
 WT - psyB      0.0199 0.0149  9   1.340  0.2131

infstat = inf:
 contrast estimate     SE df t.ratio p.value
 WT - psyB      0.0199 0.0149  9   1.340  0.2131
```

Fig. 4

### Time to 50% OD loss (**Table S3**)

**Model**

```
Call:
glm(formula = time50 ~ infstat * strain, family = Gamma(link =
"inverse"))

Deviance Residuals:
     Min        1Q    Median        3Q       Max
-0.06150  -0.01456   0.00000   0.01442   0.06346

Coefficients: (1 not defined because of singularities)
                       Estimate Std. Error t value Pr(>|t|)
(Intercept)            0.027108   0.000753  35.999 3.06e-08 ***
infstatinf             0.016370   0.001423  11.501 2.60e-05 ***
strainpsyB             0.012077   0.001960   6.163 0.000838 ***
infstatinf:strainpsyB        NA         NA      NA       NA
---
Signif. codes:  0 ‘***’ 0.001 ‘**’ 0.01 ‘*’ 0.05 ‘.’ 0.1 ‘ ’ 1

(Dispersion parameter for Gamma family taken to be 0.002315001)
Null deviance: 0.831336  on 8  degrees of freedom
Residual deviance: 0.013916  on 6  degrees of freedom
(3 observations deleted due to missingness)
AIC: 33.085
Number of Fisher Scoring iterations: 3
```

**Analysis of Deviance Table**

```
Model: Gamma, link: inverse
Response: time50
Terms added sequentially (first to last)

             Df Deviance Resid. Df Resid. Dev  Pr(>Chi)
NULL                             8    0.83134
infstat        1  0.72752        7    0.10382 < 2.2e-16 ***
strain         1  0.08990        6    0.01392 4.612e-10 ***
infstat:strain 0  0.00000        6    0.01392
---
Signif. codes:  0 ‘***’ 0.001 ‘**’ 0.01 ‘*’ 0.05 ‘.’ 0.1 ‘ ’ 1
```

**Simultaneous Tests for General Linear Hypotheses**

```
strain = WT:
 contrast   estimate   SE df t.ratio p.value
 non - inf     13.9 1.21  6  11.501  <.0001

strain = psyB:
 contrast   estimate   SE df t.ratio p.value
 non - inf   nonEst   NA NA      NA      NA

------

infstat = non:
 contrast     estimate    SE df t.ratio p.value
 WT - psyB      nonEst    NA NA      NA      NA

infstat = inf:
 contrast     estimate    SE df t.ratio p.value
 WT - psyB           5 0.811  6   6.163  0.0008
```

### % regrowth (**Table S3**)

**Model**

**NOTE:** data analysed as proportions rather than percentages

```
Call:
lm(formula = proprgw ~ infstat * strain)

Residuals:
      Min        1Q    Median        3Q       Max
-0.058283 -0.026141 -0.001702  0.011912  0.059876

Coefficients:
                 Estimate Std. Error t value Pr(>|t|)
(Intercept)           -0.002766   0.023594  -0.117 0.909573
infstatinf             0.011014   0.033367   0.330 0.749822
strainpsyB             0.028907   0.033367   0.866 0.411529
infstatinf:strainpsyB  0.244035   0.047188   5.172 0.000852 ***
---
Signif. codes:  0 ‘***’ 0.001 ‘**’ 0.01 ‘*’ 0.05 ‘.’ 0.1 ‘ ’ 1

Residual standard error: 0.04087 on 8 degrees of freedom
Multiple R-squared:  0.9255,  Adjusted R-squared:  0.8976
F-statistic: 33.15 on 3 and 8 DF,  p-value: 7.332e-05
```

**Analysis of Variance Table**

```
Response: proprgw
               Df   Sum Sq  Mean Sq F value    Pr(>F)
infstat         1 0.053092 0.053092  31.791 0.0004880 ***
strain          1 0.068335 0.068335  40.919 0.0002099 ***
infstat:strain  1 0.044665 0.044665  26.745 0.0008515 ***
Residuals       8 0.013360 0.001670
---
Signif. codes:  0 ‘***’ 0.001 ‘**’ 0.01 ‘*’ 0.05 ‘.’ 0.1 ‘ ’ 1
```

**Simultaneous Tests for General Linear Hypotheses**

```
strain = WT:
 contrast  estimate     SE df t.ratio p.value
 non - inf   -0.011 0.0334  8  -0.330  0.7498

strain = psyB:
 contrast  estimate     SE df t.ratio p.value
 non - inf   -0.255 0.0334  8  -7.644  0.0001
```

#### Fig. 4

##### *Fig. 4c*

#### OD~time*infection status relationship

**Model 0-10h (panel)**

```
Call:
lm(formula = od ~ time * strain * infstat)

Residuals:
     Min       1Q   Median       3Q      Max 
-0.87635 -0.15403  0.00273  0.19268  0.88700 

Coefficients:
                           Estimate Std. Error t value Pr(>|t|)    
(Intercept)                 3.46727    0.10635  32.603  < 2e-16 ***
time                       -0.13133    0.01798  -7.306 2.94e-11 ***
strainpsyB                 -0.30136    0.15040  -2.004   0.0473 *  
infstatinf                 -0.15364    0.15040  -1.022   0.3090    
time:strainpsyB             0.02045    0.02542   0.805   0.4226    
time:infstatinf            -0.05939    0.02542  -2.336   0.0211 *  
strainpsyB:infstatinf       0.14491    0.21269   0.681   0.4970    
time:strainpsyB:infstatinf -0.08015    0.03595  -2.229   0.0276 *  
---
Signif. codes:  0 ‘***’ 0.001 ‘**’ 0.01 ‘*’ 0.05 ‘.’ 0.1 ‘ ’ 1

Residual standard error: 0.3265 on 124 degrees of freedom
Multiple R-squared:  0.8132,	Adjusted R-squared:  0.8026 
F-statistic: 77.11 on 7 and 124 DF,  p-value: < 2.2e-16
```

**Analysis of Variance Table**

```
Response: od
                     Df Sum Sq Mean Sq  F value    Pr(>F)    
time                  1 38.526  38.526 361.2872 < 2.2e-16 ***
strain                1  3.529   3.529  33.0912 6.475e-08 ***
infstat               1 11.044  11.044 103.5729 < 2.2e-16 ***
time:strain           1  0.127   0.127   1.1911    0.2772    
time:infstat          1  3.265   3.265  30.6176 1.783e-07 ***
strain:infstat        1  0.540   0.540   5.0631    0.0262 *  
time:strain:infstat   1  0.530   0.530   4.9695    0.0276 *  
Residuals           124 13.223   0.107                       
---
Signif. codes:  0 ‘***’ 0.001 ‘**’ 0.01 ‘*’ 0.05 ‘.’ 0.1 ‘ ’ 1
```

**Model 26-30h (panel)**

```
Call:
glm(formula = od ~ time * strain * infstat, family = 
inverse.gaussian(link = "log"))

Deviance Residuals: 
     Min        1Q    Median        3Q       Max  
-0.30534  -0.10197  -0.02224   0.05785   0.52859  

Coefficients:
                            Estimate Std. Error t value Pr(>|t|)  
(Intercept)                5.518e-01  1.176e+00   0.469   0.6426  
time                      -1.131e-02  4.189e-02  -0.270   0.7892  
strainpsyB                -2.842e-01  1.699e+00  -0.167   0.8684  
infstatinf                -1.302e+00  1.327e+00  -0.982   0.3347  
time:strainpsyB            1.324e-02  6.056e-02   0.219   0.8286  
time:infstatinf            3.679e-05  4.725e-02   0.001   0.9994  
strainpsyB:infstatinf     -3.618e+00  1.890e+00  -1.914   0.0659 .
time:strainpsyB:infstatinf  1.197e-01  6.744e-02   1.775   0.0869 .
---
Signif. codes:  0 ‘***’ 0.001 ‘**’ 0.01 ‘*’ 0.05 ‘.’ 0.1 ‘ ’ 1

(Dispersion parameter for inverse.gaussian family taken to be 
0.03328725)
Null deviance: 28.69690  on 35  degrees of freedom
Residual deviance:  0.83681  on 28  degrees of freedom
AIC: -63.482
Number of Fisher Scoring iterations: 6
```

**Analysis of Deviance Table**

```
Model: inverse.gaussian, link: log
Response: od
Terms added sequentially (first to last)

                    Df Deviance Resid. Df Resid. Dev  Pr(>Chi)    
NULL                                   35    28.6969              
time                 1   0.0056        34    28.6913   0.68158    
strain               1   0.0154        33    28.6759   0.49638    
infstat              1  26.7958        32     1.8801 < 2.2e-16 ***
time:strain          1   0.5784        31     1.3017 3.067e-05 ***
time:infstat         1   0.1712        30     1.1305   0.02335 *  
strain:infstat       1   0.1889        29     0.9416   0.01720 *  
time:strain:infstat  1   0.1048        28     0.8368   0.07606 .  
---
Signif. codes:  0 ‘***’ 0.001 ‘**’ 0.01 ‘*’ 0.05 ‘.’ 0.1 ‘ ’ 1
```

Fig. 4

### Maximum decline rates (**Table S3**)

**Model (0-10h #1)**

**NOTE:** two models used for 0-10h, the second with an influential point removed for model validation

```
Call:
lm(formula = mxdecrt ~ strain * infstat)

Residuals:
      Min        1Q    Median        3Q       Max
-0.112315 -0.007126 -0.001099  0.002566  0.207824

Coefficients:
                      Estimate Std. Error t value Pr(>|t|)
(Intercept)           -0.10250    0.05222  -1.963   0.0853 .
strainpsyB            -0.22362    0.07385  -3.028   0.0164 *
infstatnon             0.05409    0.07385   0.732   0.4848
strainpsyB:infstatnon  0.20506    0.10445   1.963   0.0852 .
---
Signif. codes:  0 ‘***’ 0.001 ‘**’ 0.01 ‘*’ 0.05 ‘.’ 0.1 ‘ ’ 1

Residual standard error: 0.09045 on 8 degrees of freedom
Multiple R-squared:  0.695,   Adjusted R-squared:  0.5806
F-statistic: 6.075 on 3 and 8 DF,  p-value: 0.01852
```

**Analysis of Variance Table**

```
Response: mxdecrt
               Df   Sum Sq  Mean Sq F value  Pr(>F)
strain          1 0.043986 0.043986  5.3762 0.04902 *
infstat         1 0.073589 0.073589  8.9944 0.01710 *
strain:infstat  1 0.031538 0.031538  3.8547 0.08522 .
Residuals       8 0.065453 0.008182
---
Signif. codes:  0 ‘***’ 0.001 ‘**’ 0.01 ‘*’ 0.05 ‘.’ 0.1 ‘ ’ 1
```

**Simultaneous Tests for General Linear Hypotheses**

```
infstat = inf:
 contrast  estimate     SE df t.ratio p.value
 wt - psyB   0.2236 0.0739  8   3.028  0.0164

infstat = non:
 contrast  estimate     SE df t.ratio p.value
 wt - psyB   0.0186 0.0739  8   0.251  0.8080
```

**------**

```
strain = wt:
 contrast  estimate     SE df t.ratio p.value
 inf - non  -0.0541 0.0739  8  -0.732  0.4848

strain = psyB:
 contrast  estimate     SE df t.ratio p.value
 inf - non  -0.2592 0.0739  8  -3.509  0.0080
```

**Model (0-10h #2, influential point 8 removed)**

```
Call:
lm(formula = mxdecrt[-8] ~ strain[-8] * infstat[-8])

Residuals:
      Min        1Q    Median        3Q       Max
-0.013663 -0.003800 -0.001073  0.003734  0.016317

Coefficients:
                      Estimate Std. Error t value Pr(>|t|)
(Intercept)          -0.102502   0.005636 -18.188 3.76e-07 ***
strain[-8]psyB       -0.327530   0.008911 -36.756 2.87e-09 ***
infstat[-8]non        0.054088   0.007970   6.786 0.000256 ***
strain[-8]psyB:       0.308975   0.011955  25.844 3.32e-08 ***
infstat[-8]non
---
Signif. codes:  0 ‘***’ 0.001 ‘**’ 0.01 ‘*’ 0.05 ‘.’ 0.1 ‘ ’ 1

Residual standard error: 0.009761 on 7 degrees of freedom
Multiple R-squared:  0.9969,  Adjusted R-squared:  0.9956
F-statistic: 747.1 on 3 and 7 DF,  p-value: 3.916e-09
```

**Analysis of Variance Table**

```
Response: mxdecrt[-8]
                      Df   Sum Sq  Mean Sq F value    Pr(>F)
strain[-8]             1 0.050992 0.050992  535.15 7.154e-08 ***
infstat[-8]            1 0.098922 0.098922 1038.18 7.174e-09 ***
strain[-8]:infstat[-8] 1 0.063644 0.063644  667.93 3.320e-08 ***
Residuals              7 0.000667 0.000095
---
Signif. codes:  0 ‘***’ 0.001 ‘**’ 0.01 ‘*’ 0.05 ‘.’ 0.1 ‘ ’ 1
```

**Simultaneous Tests for General Linear Hypotheses**

```
infstat = inf:
 contrast  estimate      SE df t.ratio p.value
 wt - psyB   0.3275 0.00891  7  36.756  <.0001

infstat = non:
 contrast  estimate      SE df t.ratio p.value
 wt - psyB   0.0186 0.00797  7   2.328  0.0528
```

**------**

```
strain = wt:
 contrast  estimate      SE df t.ratio p.value
 inf - non  -0.0541 0.00797  7  -6.786  0.0003

strain = psyB:
 contrast  estimate      SE df t.ratio p.value
 inf - non  -0.3631 0.00891  7 -40.744  <.0001
```

**Model (26-30h)**

```
Call:
lm(formula = mxdecrt ~ strain * infstat)

Residuals:
      Min        1Q    Median        3Q       Max
-0.040730 -0.020260 -0.002445  0.011398  0.070128

Coefficients:
                       Estimate Std. Error t value Pr(>|t|)
(Intercept)           -0.022108   0.019645  -1.125 0.293066
strainpsyB             0.141857   0.0psyB3   5.106 0.000923 ***
infstatnon             0.002742   0.0psyB3   0.099 0.923805
strainpsyB:infstatnon -0.132118   0.039291  -3.363 0.009894 **
---
Signif. codes:  0 ‘***’ 0.001 ‘**’ 0.01 ‘*’ 0.05 ‘.’ 0.1 ‘ ’ 1

Residual standard error: 0.03403 on 8 degrees of freedom
Multiple R-squared:  0.8206,  Adjusted R-squared:  0.7533
F-statistic: 12.19 on 3 and 8 DF,  p-value: 0.002361
```

**Analysis of Variance Table**

```
Response: mxdecrt
              Df    Sum Sq   Mean Sq F value   Pr(>F)
strain         1 0.0172357 0.0172357  14.886 0.004821 **
infstat        1 0.0120272 0.0120272  10.388 0.012184 *
strain:infstat 1 0.0130915 0.0130915  11.307 0.009894 **
Residuals      8 0.0092626 0.0011578
---
Signif. codes:  0 ‘***’ 0.001 ‘**’ 0.01 ‘*’ 0.05 ‘.’ 0.1 ‘ ’ 1
```

**Simultaneous Tests for General Linear Hypotheses**

```
infstat = inf:
 contrast  estimate     SE df t.ratio p.value
 wt - psyB -0.14186 0.0278  8  -5.106  0.0009

infstat = non:
 contrast  estimate     SE df t.ratio p.value
 wt - psyB -0.00974 0.0278  8  -0.351  0.7350

------

strain = wt:
 contrast  estimate     SE df t.ratio p.value
 inf - non -0.00274 0.0278  8  -0.099  0.9238

strain = psyB:
 contrast  estimate     SE df t.ratio p.value
 inf - non  0.12938 0.0278  8   4.657  0.0016
```

Fig. 4

#### Times at max decline rates (**Table S3**)

**Model (0-10h)**

```
Call:
lm(formula = tmxdec ~ strain + infstat)
Residuals:
    Min      1Q  Median      3Q     Max
-2.5000 -2.1667 -0.3333  0.8333  5.5000

Coefficients:
             Estimate Std. Error t value Pr(>|t|)
(Intercept)    9.1667     1.4175   6.467 0.000116 ***
strainpsyB     0.3333     1.6368   0.204 0.843155
infstatnon    -6.0000     1.6368  -3.666 0.005189 **
---
Signif. codes:  0 ‘***’ 0.001 ‘**’ 0.01 ‘*’ 0.05 ‘.’ 0.1 ‘ ’ 1
Residual standard error: 2.835 on 9 degrees of freedom
Multiple R-squared:  0.5996,  Adjusted R-squared:  0.5107
F-statistic:  6.74 on 2 and 9 DF,  p-value: 0.01626
```

**Analysis of Variance Table**

```
Response: tmxdec
            Df  Sum Sq Mean Sq F value   Pr(>F)
strain       1   0.333   0.333  0.0415 0.843155
infstat      1 108.000 108.000 13.4378 0.005189 **
Residuals    9  72.333   8.037
Signif. codes:  0 ‘***’ 0.001 ‘**’ 0.01 ‘*’ 0.05 ‘.’ 0.1 ‘ ’ 1
```

**Simultaneous Tests for General Linear Hypotheses**

```
Multiple Comparisons of Means: Tukey Contrasts
Fit: lm(formula = tmxdec ~ strain + infstat, data = lsp.stsum)

Linear Hypotheses:
               Estimate Std. Error t value Pr(>|t|)
non - inf == 0   -6.000      1.637  -3.666  0.00519 **
---
Signif. codes:  0 ‘***’ 0.001 ‘**’ 0.01 ‘*’ 0.05 ‘.’ 0.1 ‘ ’ 1
(Adjusted p values reported -- single-step method)
```

**Model (26-30h)**

```
Call:
lm(formula = tmxdec ~ strain * infstat)

Residuals:
    Min      1Q  Median      3Q     Max
-0.6667 -0.6667  0.0000  0.0000  1.3333

Coefficients:
                      Estimate Std. Error t value Pr(>|t|)
(Intercept)            28.6667     0.4714  60.811 5.94e-12 ***
strainpsyB             -0.6667     0.6667  -1.000    0.347
infstatnon             -0.6667     0.6667  -1.000    0.347
strainpsyB:infstatnon   1.3333     0.9428   1.414    0.195
---
Signif. codes:  0 ‘***’ 0.001 ‘**’ 0.01 ‘*’ 0.05 ‘.’ 0.1 ‘ ’ 1

Residual standard error: 0.8165 on 8 degrees of freedom
Multiple R-squared:    0.2,   Adjusted R-squared:   -0.1
F-statistic: 0.6667 on 3 and 8 DF,  p-value: 0.5957
```

**Analysis of Variance Table**

```
Response: tmxdec
               Df Sum Sq Mean Sq F value Pr(>F)
strain          1 0.0000 0.00000       0  1.000
infstat         1 0.0000 0.00000       0  1.000
strain:infstat  1 1.3333 1.33333       2  0.195
Residuals       8 5.3333 0.66667
```

#### Time to lysis (**Table S3**)

Not determined – rate already negative from start

#### % decrease OD (**Table S3**)

**Model (0-10h)**

**NOTE:** data were analysed as proportions rather than percentage data to allow arcsine transformation

```
Call:
glm(formula = propdec ~ strain * infstat, family = Gamma(link =
"identity"))

Deviance Residuals:
     Min        1Q    Median        3Q       Max
-0.28521  -0.07887   0.01229   0.07807   0.17975

Coefficients:
                      Estimate Std. Error t value Pr(>|t|)
(Intercept)            0.57269    0.04883  11.727 2.55e-06 ***
strainpsyB             0.20563    0.08240   2.496   0.0372 *
infstatnon            -0.18225    0.05910  -3.084   0.0150 *
strainpsyB:infstatnon -0.22863    0.09423  -2.426   0.0414 *
---
Signif. codes:  0 ‘***’ 0.001 ‘**’ 0.01 ‘*’ 0.05 ‘.’ 0.1 ‘ ’ 1

(Dispersion parameter for Gamma family taken to be 0.02181414)
Null deviance: 1.32013  on 11  degrees of freedom
Residual deviance: 0.18511  on  8  degrees of freedom
AIC: -22.658
Number of Fisher Scoring iterations: 3
```

**Analysis of Deviance Table**

```
Model: Gamma, link: identity
Response: propdec
Terms added sequentially (first to last)

              Df Dev. Resid. Df Resid. Dev      F    Pr(>F)
NULL                         11    1.32013
strain         1 0.09034     10    1.22979  4.1415 0.0762566 .
infstat        1 0.91276     9     0.31702 41.8428 0.0001944 ***
strain:infstat 1 0.13191     8     0.18511  6.0472 0.0393742 *
---
Signif. codes:  0 ‘***’ 0.001 ‘**’ 0.01 ‘*’ 0.05 ‘.’ 0.1 ‘ ’ 1
```

**Simultaneous Tests for General Linear Hypotheses**

```
infstat = inf:
 2            estimate     SE df t.ratio p.value
 WT - psyB   -0.206 0.0824  8  -2.496  0.0372

infstat = non:
 2            estimate     SE df t.ratio p.value
 WT - psyB    0.023 0.0457  8   0.503  0.6286

------

strain = WT:
 2          estimate     SE df t.ratio p.value
 inf - non    0.182 0.0591  8   3.084  0.0150

strain = psyB:
 2          estimate     SE df t.ratio p.value
 inf - non    0.411 0.0734  8   5.598  0.0005
```

### Fig. 4

**Model (26-30h)**
```
Call:
lm(formula = asin(propdec) ~ strain + infstat)

Residuals:
     Min       1Q   Median       3Q      Max
-0.21643 -0.01829  0.02210  0.04121  0.08085

Coefficients:
               Estimate Std. Error t value Pr(>|t|)
(Intercept)    1.163260   0.043498  26.743 6.92e-10 ***
strainpsyB    -0.003295   0.050227  -0.066    0.949
infstatnon    -0.479128   0.050227  -9.539 5.29e-06 ***
---
Signif. codes:  0 ‘***’ 0.001 ‘**’ 0.01 ‘*’ 0.05 ‘.’ 0.1 ‘ ’ 1

Residual standard error: 0.087 on 9 degrees of freedom
Multiple R-squared:   0.91,   Adjusted R-squared:   0.89
F-statistic:  45.5 on 2 and 9 DF,  p-value: 0.00001968
```

**Analysis of Variance Table**
```
Response: asin(propdec)
          Df  Sum Sq Mean Sq F value      Pr(>F)
strain     1 0.00003 0.00003  0.0043      0.9491
infstat    1 0.68869 0.68869 90.9963 0.000005292 ***
Residuals  9 0.06812 0.00757
---
Signif. codes:  0 ‘***’ 0.001 ‘**’ 0.01 ‘*’ 0.05 ‘.’ 0.1 ‘ ’ 1
```

**Simultaneous Tests for General Linear Hypotheses**
```
Multiple Comparisons of Means: Tukey Contrasts
Fit: lm(formula = asin(propdec) ~ strain + infstat, data =
lsp.stsum)

Linear Hypotheses:
               Estimate Std. Error t value   Pr(>|t|)
non - inf == 0 -0.47913    0.05023  -9.539 0.00000529 ***
---
Signif. codes:  0 ‘***’ 0.001 ‘**’ 0.01 ‘*’ 0.05 ‘.’ 0.1 ‘ ’ 1
(Adjusted p values reported -- single-step method)
```

---

#### *Fig. 4d*

#### PFU~time*population relationship (**Free phages**)

**Determining breakpoints (mod_bp)**
```
Call:
lm(formula = PFU ~ time * strain)

Residuals:
       Min         1Q     Median         3Q        Max
-4.021e+09 -1.093e+09 -1.662e+07  5.226e+08  8.967e+09

Coefficients:
                   Estimate Std. Error t value Pr(>|t|)
(Intercept)      -212724687  840517664  -0.253  0.80156
time              458572241  149589024   3.066  0.00399 **
strainpsyB       -849262873 1188671480  -0.714  0.47931
time:strainpsyB    80972240  211550826   0.383  0.70403
---
Signif. codes:  0 ‘***’ 0.001 ‘**’ 0.01 ‘*’ 0.05 ‘.’ 0.1 ‘ ’ 1

Residual standard error: 2.371e+09 on 38 degrees of freedom
Multiple R-squared:  0.3756,  Adjusted R-squared:  0.3263
F-statistic: 7.619 on 3 and 38 DF,  p-value: 0.0004156
```

*****Regression Model with Segmented Relationship(s)*****
```
Call:
segmented.lm(obj = mod_bp, seg.Z = ~time, psi = c(1, 4))

Estimated Break-Point(s):
              Est. St.Err
psi1.time    1.383  5.971
psi2.time    7.313  1.126

Meaningful coefficients of the linear terms:
                  Estimate Std. Error t value Pr(>|t|)
(Intercept)      679633103 1083054860   0.628    0.535
time            -120436453 1309093272  -0.092    0.927
strainpsyB      -849262873 1133547754  -0.749    0.459
U1.time          278594500 1345370694   0.207       NA
U2.time         1561864167  729629702   2.141       NA
time:strainpsyB   80972240  201740319   0.401    0.691

Residual standard error: 2.261e+09 on 34 degrees of freedom
Multiple R-Squared: 0.4919,  Adjusted R-squared: 0.3873
Convergence attained in 2 iter. (rel. change 0)
```

**Model (panel)**
```
Call:
lm(formula = log(PFU) ~ time * (time < 1.383) + time *
    (time >= 1.383 & time <= 7.313) * strain)

Residuals:
    Min      1Q  Median      3Q     Max
-3.9493 -0.6495 -0.0513  0.6570  3.1714

Coefficients:


Estimate Std. Error t value Pr(>|t|)
(Intercept)             18.19975    4.08770   4.452 9.68e-05 ***
time                     0.41501    0.45436   0.913 0.367864
time < 1.383TRUE         2.68086    4.10969   0.652 0.518849
time >= 1.383 &          1.43008    4.31114   0.332 0.742265
time <= 7.313TRUE
strain2778              -7.06674    0.94215  -7.501 1.54e-08 ***
time:time               -2.62334    1.00266  -2.616 0.013451 *
< 1.383TRUE
time:time >= 1.383      -0.12727    0.55406  -0.230 0.819785
& time <= 7.313TRUE
time:strain2778          0.63583    0.14669   4.334 0.000136 ***
time >= 1.383           -1.21425    2.15427  -0.564 0.576924
& time <= 7.313TRUE:
strain2778
time:time >= 1.383       0.09799    0.47179   0.208 0.836787
& time <= 7.313TRUE:
strain2778
---
Signif. codes:  0 ‘***’ 0.001 ‘**’ 0.01 ‘*’ 0.05 ‘.’ 0.1 ‘ ’ 1

Residual standard error: 1.553 on 32 degrees of freedom
Multiple R-squared:  0.8572,  Adjusted R-squared:  0.8171
F-statistic: 21.35 on 9 and 32 DF,  p-value: 4.003e-11
```

**Analysis of Variance Table**
```
Response: log(PFU)

                        Df  Sum Sq Mean Sq F value    Pr(>F)
time                     1 165.170 165.170 68.4559 1.880e-09 ***
time < 1.383             1   0.614   0.614  0.2543  0.617509
time >= 1.383            1   0.318   0.318  0.1319  0.718835
& time <= 7.313
strain                   1 222.502 222.502 92.2178 6.050e-11 ***
time:time < 1.383        1  18.735  18.735  7.7649  0.008883 **
time:time >= 1.383       1   0.059   0.059  0.0244  0.876908
& time <= 7.313
time:strain              1  54.339  54.339 22.5211 4.156e-05 ***
time >= 1.383            1   1.689   1.689  0.7001  0.408961
& time <= 7.313:
strain
time:time >=             1   0.104   0.104  0.0431  0.836787
1.383 & time <=
7.313:strain
Residuals               32  77.209   2.413
---
Signif. codes:  0 ‘***’ 0.001 ‘**’ 0.01 ‘*’ 0.05 ‘.’ 0.1 ‘ ’ 1
```

#### Fig. 4

### Maximum PFU rate & time at max PFU rate (**Table S3**)

includes comparison between both phage populations (data from Fig. 5d)

**Model (Maximum PFU production rate)**
**NOTE:** also includes data from sections below on total PFU

```
Call:
glm(formula = mxpfurt ~ strain + ppn, family = Gamma(link = 
log))

Deviance Residuals:
     Min        1Q    Median        3Q       Max
-0.73627  -0.35097  -0.07997   0.18590   0.68587

Coefficients:
              Estimate Std. Error t value Pr(>|t|)
(Intercept)    -0.5599     0.2373  -2.359 0.042645 *
strainpsyB      1.4546     0.2740   5.308 0.000489 ***
ppntotal       -0.7549     0.2740  -2.755 0.022300 *
---
Signif. codes:  0 '***' 0.001 '**' 0.01 '*' 0.05 '.' 0.1 ' ' 1

(Dispersion parameter for Gamma family taken to be 0.2252953)
Null deviance: 9.1973  on 11  degrees of freedom
Residual deviance: 2.0737  on  9  degrees of freedom
AIC: 14.217
Number of Fisher Scoring iterations: 4
```

**Analysis of Deviance Table**

```
Model: Gamma, link: log
Response: mxpfurt
Terms added sequentially (first to last)

        Df Deviance Resid. Df Resid. Dev  Pr(>Chi)
NULL                       11     9.1973
strain    1   5.4615        10     3.7357 8.497e-07 ***
ppn  1   1.6620         9     2.0737  0.006606 **
---
Signif. codes:  0 '***' 0.001 '**' 0.01 '*' 0.05 '.' 0.1 ' ' 1
```

**Simultaneous Tests for General Linear Hypotheses**

```
ppn = free:
 contrast    ratio    SE df null t.ratio p.value
 WT / psyB   0.233 0.064  9    1  -5.308  0.0005

ppn = total:
 contrast    ratio    SE df null t.ratio p.value
 WT / psyB   0.233 0.064  9    1  -5.308  0.0005

Tests are performed on the log scale

------

strain = WT:
 contrast     ratio    SE df null t.ratio p.value
 free / total  2.13 0.583  9    1   2.755  0.0223

strain = psyB:
 contrast     ratio    SE df null t.ratio p.value
 free / total  2.13 0.583  9    1   2.755  0.0223

Tests are performed on the log scale
```

**Model (Time at max PFU production rate)**

```
Call:
lm(formula = log(tmxpfu) ~ strain + ppn)

Residuals:
    Min      1Q  Median      3Q     Max
-1.1293 -0.3576 -0.1307  0.5590  1.1733

Coefficients:
              Estimate Std. Error t value Pr(>|t|)
(Intercept)     1.1293     0.4097   2.756   0.0222 *
strainpsyB     -0.7979     0.4731  -1.687   0.1259
ppntotal       -0.2007     0.4731  -0.424   0.6814
---
Signif. codes:  0 '***' 0.001 '**' 0.01 '*' 0.05 '.' 0.1 ' ' 1

Residual standard error: 0.8194 on 9 degrees of freedom
Multiple R-squared:  0.2515,  Adjusted R-squared:  0.08522
F-statistic: 1.512 on 2 and 9 DF,  p-value: 0.2715
```

### Fold change in PFU (**Table S3**)

includes comparison between both phage populations (data from Fig. 5d)

**Model (10h)**
**NOTE:** also includes data from sections below on total PFU

```
Call:
lm(formula = fch ~ strain * ppn)

Residuals:
    Min      1Q  Median      3Q     Max
-3.0127 -1.2950 -0.4343  1.2600  3.4813

Coefficients:
                         Estimate Std. Error t value Pr(>|t|)
(Intercept)                2.8850     1.3219   2.182   0.0606 .
strainpsyB                 0.2837     1.8695   0.152   0.8832
ppntotal                  -0.4150     1.8695  -0.222   0.8299
strainpsyB:ppntotal        3.2463     2.6439   1.228   0.2544
---
Signif. codes:  0 '***' 0.001 '**' 0.01 '*' 0.05 '.' 0.1 ' ' 1

Residual standard error: 2.29 on 8 degrees of freedom
Multiple R-squared:  0.3561,  Adjusted R-squared:  0.1146
F-statistic: 1.475 on 3 and 8 DF,  p-value: 0.293
```

**Analysis of Variance Table**

```
Response: fch
                Df Sum Sq Mean Sq F value Pr(>F)
strain           1 10.908 10.9080  2.0807 0.1872
ppn              1  4.379  4.3790  0.8353 0.3875
strain:ppn       1  7.904  7.9040  1.5077 0.2544
Residuals        8 41.941  5.2426
```

**Model (30h)**

```
Call:
lm(formula = fch ~ strain + ppn)

Residuals:
    Min      1Q  Median      3Q     Max
-1.3292 -1.1479 -0.1458  0.8187  1.7208

Coefficients:
              Estimate Std. Error t value Pr(>|t|)
(Intercept)     4.7875     0.6269   7.636  3.2e-05 ***
strainpsyB     -2.0750     0.7239  -2.866   0.0186 *
ppntotal        0.8417     0.7239   1.163   0.2749
---
Signif. codes:  0 '***' 0.001 '**' 0.01 '*' 0.05 '.' 0.1 ' ' 1

Residual standard error: 1.254 on 9 degrees of freedom
Multiple R-squared:  0.5153,  Adjusted R-squared:  0.4076
F-statistic: 4.784 on 2 and 9 DF,  p-value: 0.03843
```

**Analysis of Variance Table**

```
Response: fch
          Df  Sum Sq Mean Sq F value  Pr(>F)
strain     1 12.9169 12.9169  8.2156 0.01859 *
ppn        1  2.1252  2.1252  1.3517 0.27488
Residuals  9 14.1502  1.5722
---
Signif. codes:  0 '***' 0.001 '**' 0.01 '*' 0.05 '.' 0.1 ' ' 1
```

**Simultaneous Tests for General Linear Hypotheses**

```
Multiple Comparisons of Means: Tukey Contrasts
Fit: lm(formula = fch ~ strain + ppn, data = prg.stsum)

Linear Hypotheses:
                 Estimate Std. Error t value Pr(>|t|)
psyB - WT == 0    -2.0750     0.7239  -2.866   0.0186 *
---
Signif. codes:  0 '***' 0.001 '**' 0.01 '*' 0.05 '.' 0.1 ' ' 1
(Adjusted p values reported -- single-step method)
```

---

#### *Fig. 4e*

#### PFU~time*population relationship (**Total phages**)

**Determining breakpoints (mod_bp)**

```
Call:
lm(formula = PFU ~ time * strain)

Residuals:
       Min         1Q     Median         3Q        Max 
-5.524e+09 -1.065e+09 -1.703e+07  7.536e+08  8.096e+09 

Coefficients:
                        Estimate Std. Error t value Pr(>|t|)    
(Intercept)            1.539e+09  9.038e+08   1.703 0.096714 .  
time                   3.485e+08  1.609e+08   2.167 0.036585 *  
strainpsyB            -3.514e+09  1.278e+09  -2.750 0.009085 ** 
time:strainpsyB        9.489e+08  2.275e+08   4.171 0.000169 ***
---
Signif. codes:  0 '***' 0.001 '**' 0.01 '*' 0.05 '.' 0.1 ' ' 1

Residual standard error: 2.549e+09 on 38 degrees of freedom
Multiple R-squared:  0.6498,  Adjusted R-squared:  0.6222 
F-statistic: 23.51 on 3 and 38 DF,  p-value: 8.992e-09
```

*****Regression Model with Segmented Relationship(s)*****

```
Call:
segmented.lm(obj = mod_bp, seg.Z = ~time, psi = c(4))

Estimated Break-Point(s):
           Est. St.Err
psi1.time 4.278  1.647

Meaningful coefficients of the linear terms:
                  Estimate Std. Error t value Pr(>|t|)    
(Intercept)      2.536e+09  9.916e+08   2.558 0.014898 *  
time            -2.594e+08  3.564e+08  -0.728 0.471448    
strainpsyB      -3.514e+09  1.232e+09  -2.853 0.007143 ** 
U1.time          1.063e+09  4.907e+08   2.166       NA    
time:strainpsyB  9.489e+08  2.193e+08   4.328 0.000115 ***
---
Signif. codes:  0 '***' 0.001 '**' 0.01 '*' 0.05 '.' 0.1 ' ' 1

Residual standard error: 2.457e+09 on 36 degrees of freedom
Multiple R-Squared: 0.6918,  Adjusted R-squared: 0.649 
Convergence attained in 2 iter. (rel. change 0)
```

**Model 0-10h (Table S3)**

```
Call:
lm(formula = log(PFU) ~ time * (time > 4.278) * strain)

Residuals:
     Min       1Q   Median       3Q      Max 
-1.17268 -0.44467  0.00597  0.29059  1.80693 

Coefficients:
                       Estimate Std. Error t value Pr(>|t|)    
(Intercept)            21.21330    0.31608  67.114  < 2e-16 ***
time                    0.11843    0.13795   0.858   0.3966    
time > 4.278TRUE        0.59749    1.21962   0.490   0.6274    
strainpsyB             -5.60886    0.44700 -12.548 2.59e-14 ***
time:time > 4.278TRUE  -0.07761    0.19961  -0.389   0.6998    
time:strainpsyB         1.14925    0.19509   5.891 1.19e-06 ***
time > 4.278TRUE:       2.80418    1.72480   1.626   0.1132    
 strainpsyB
time:time > 4.278TRUE: -0.75388    0.28229  -2.671   0.0115 *  
 strainpsyB
---
Signif. codes:  0 '***' 0.001 '**' 0.01 '*' 0.05 '.' 0.1 ' ' 1

Residual standard error: 0.7068 on 34 degrees of freedom
Multiple R-squared:  0.9218,  Adjusted R-squared:  0.9057 
F-statistic: 57.22 on 7 and 34 DF,  p-value: < 2.2e-16
```

**Analysis of Variance Table**

```
Response: log(PFU)
                        Df Sum Sq Mean Sq  F value    Pr(>F)    
time                     1 97.297  97.297 194.7784 1.233e-15 ***
time > 4.278             1  0.161   0.161   0.3229  0.573585    
strain                   1 37.995  37.995  76.0617 3.430e-10 ***
time:time > 4.278        1  5.181   5.181  10.3716  0.002816 ** 
time:strain              1 55.400  55.400 110.9054 3.049e-12 ***
time > 4.278:strain      1  0.491   0.491   0.9821  0.328684    
time:time > 4.278:strain 1  3.563   3.563   7.1322  0.011531 *  
Residuals               34 16.984   0.500                       
---
Signif. codes:  0 '***' 0.001 '**' 0.01 '*' 0.05 '.' 0.1 ' ' 1
```

**Test 30h**
**Welch Two Sample t-test (Table S3)**

```
t = 1.8562, df = 3.953, p-value = 0.1379
alternative hypothesis: true difference in means is not equal to
0
95 percent confidence interval:
 -2229153848 11095820515
sample estimates:
  mean of x   mean of y 
11400000000  6966666667 
```

==================================================================

### Figure 5

#### *Fig. 5b*

#### Fold change~time*infection status relationships

**Model 8 min EXP NON-INF normalised (upper section of panel)**

```
Call:
lm(formula = log10fch ~ time.fac * gene * infstat)

Residuals:
     Min       1Q   Median       3Q      Max 
-0.30556 -0.11217 -0.00792  0.10750  0.40135 

Coefficients:
                             Estimate Std. Error t value Pr(>|t|)    
(Intercept)                  0.179419   0.103268   1.737 0.088726 .  
time.fac8                   -0.179419   0.146043  -1.229 0.225233    
geneG24.1                    0.001971   0.146043   0.013 0.989286    
geneG26                      0.001971   0.146043   0.013 0.989286    
geneG35                      0.083054   0.146043   0.569 0.572214    
geneG46                      0.001971   0.146043   0.013 0.989286    
geneG6                       0.062381   0.146043   0.427 0.671186    
infstatinf                   5.492944   0.146043  37.612  < 2e-16 ***
time.fac8:geneG24.1         -0.001971   0.206535  -0.010 0.992424    
time.fac8:geneG26           -0.001971   0.206535  -0.010 0.992424    
time.fac8:geneG35           -0.083054   0.206535  -0.402 0.689375    
time.fac8:geneG46           -0.001971   0.206535  -0.010 0.992424    
time.fac8:geneG6            -0.062381   0.206535  -0.302 0.763930    
time.fac8:infstatinf        -1.671708   0.206535  -8.094 1.59e-10 ***
geneG24.1:infstatinf        -0.740861   0.206535  -3.587 0.000782 ***
geneG26:infstatinf          -0.331871   0.206535  -1.607 0.114647    
geneG35:infstatinf          -0.950510   0.206535  -4.602 3.08e-05 ***
geneG46:infstatinf          -1.524316   0.206535  -7.380 1.92e-09 ***
geneG6:infstatinf           -0.501161   0.206535  -2.427 0.019050 *  
time.fac8:geneG24.1:infstatinf -0.139022   0.292085  -0.476 0.636260    
time.fac8:geneG26:infstatinf  -0.029753   0.292085  -0.102 0.919288    
time.fac8:geneG35:infstatinf   1.719484   0.292085   5.887 3.73e-07 ***
time.fac8:geneG46:infstatinf   2.520733   0.292085   8.630 2.50e-11 ***
time.fac8:geneG6:infstatinf    0.100922   0.292085   0.346 0.731211    
---
Signif. codes:  0 ‘***’ 0.001 ‘**’ 0.01 ‘*’ 0.05 ‘.’ 0.1 ‘ ’ 1
Residual standard error: 0.1789 on 48 degrees of freedom
Multiple R-squared:  0.9958,  Adjusted R-squared:  0.9938 
F-statistic: 492.7 on 23 and 48 DF,  p-value: < 2.2e-16
```

**Analysis of Variance Table**

```
Response: log10fch
                    Df Sum Sq Mean Sq    F value    Pr(>F)    
time.fac             1   8.64    8.64   270.0472 < 2.2e-16 ***
gene                 5   1.34    0.27     8.3481 9.743e-06 ***
infstat              1 337.48  337.48 10548.6837 < 2.2e-16 ***
time.fac:gene        5   4.71    0.94    29.4144 1.546e-13 ***
time.fac:infstat     1   4.29    4.29   134.0731 1.684e-15 ***
gene:infstat         5   1.25    0.25     7.8027 1.943e-05 ***
time.fac:gene:infstat  5   4.83    0.97    30.1930 9.711e-14 ***
Residuals           48   1.54    0.03                         
---
Signif. codes:  0 ‘***’ 0.001 ‘**’ 0.01 ‘*’ 0.05 ‘.’ 0.1 ‘ ’ 1
```

**Simultaneous Tests for General Linear Hypotheses**

```
time.fac = 25, gene = G11:
 contrast  estimate    SE df t.ratio p.value
 non - inf    -5.49 0.146 48 -37.612  <.0001

time.fac = 8, gene = G11:
 contrast  estimate    SE df t.ratio p.value
 non - inf    -3.82 0.146 48 -26.165  <.0001

time.fac = 25, gene = G24.1:
 contrast  estimate    SE df t.ratio p.value
 non - inf    -4.75 0.146 48 -32.539  <.0001

time.fac = 8, gene = G24.1:
 contrast  estimate    SE df t.ratio p.value
 non - inf    -2.94 0.146 48 -20.140  <.0001

time.fac = 25, gene = G26:
 contrast  estimate    SE df t.ratio p.value
 non - inf    -5.16 0.146 48 -35.340  <.0001

time.fac = 8, gene = G26:
 contrast  estimate    SE df t.ratio p.value
 non - inf    -3.46 0.146 48 -23.689  <.0001

time.fac = 25, gene = G35:
 contrast  estimate    SE df t.ratio p.value
 non - inf    -4.54 0.146 48 -31.104  <.0001

time.fac = 8, gene = G35:
 contrast  estimate    SE df t.ratio p.value
 non - inf    -4.59 0.146 48 -31.431  <.0001

time.fac = 25, gene = G46:
 contrast  estimate    SE df t.ratio p.value
 non - inf    -3.97 0.146 48 -27.174  <.0001

time.fac = 8, gene = G46:
 contrast  estimate    SE df t.ratio p.value
 non - inf    -4.82 0.146 48 -32.988  <.0001

time.fac = 25, gene = G6:
 contrast  estimate    SE df t.ratio p.value
 non - inf    -4.99 0.146 48 -34.180  <.0001

time.fac = 8, gene = G6:
 contrast  estimate    SE df t.ratio p.value
 non - inf    -3.42 0.146 48 -23.425  <.0001

------

infstat = non, gene = G11:
 contrast  estimate    SE df t.ratio p.value
 25 - 8       0.179 0.146 48   1.229  0.2252

infstat = inf, gene = G11:
 contrast  estimate    SE df t.ratio p.value
 25 - 8       1.851 0.146 48  12.675  <.0001

infstat = non, gene = G24.1:
 contrast  estimate    SE df t.ratio p.value
 25 - 8       0.181 0.146 48   1.242  0.2203

infstat = inf, gene = G24.1:
 contrast  estimate    SE df t.ratio p.value
 25 - 8       1.992 0.146 48  13.641  <.0001

infstat = non, gene = G26:
 contrast  estimate    SE df t.ratio p.value
 25 - 8       0.181 0.146 48   1.242  0.2203

infstat = inf, gene = G26:
 contrast  estimate    SE df t.ratio p.value
 25 - 8       1.883 0.146 48  12.892  <.0001

infstat = non, gene = G35:
 contrast  estimate    SE df t.ratio p.value
 25 - 8       0.262 0.146 48   1.797  0.0786

infstat = inf, gene = G35:
 contrast  estimate    SE df t.ratio p.value
 25 - 8       0.215 0.146 48   1.470  0.1481

infstat = non, gene = G46:
 contrast  estimate    SE df t.ratio p.value
 25 - 8       0.181 0.146 48   1.242  0.2203

infstat = inf, gene = G46:
 contrast  estimate    SE df t.ratio p.value
 25 - 8      -0.668 0.146 48  -4.572  <.0001

infstat = non, gene = G6:
 contrast  estimate    SE df t.ratio p.value
 25 - 8       0.242 0.146 48   1.656  0.1043

infstat = inf, gene = G6:
 contrast  estimate    SE df t.ratio p.value
 25 - 8       1.813 0.146 48  12.411  <.0001

------

time.fac = 25, infstat = non:
 contrast     estimate    SE df t.ratio p.value
 G11 - G24.1 -0.00197 0.146 48  -0.013  1.0000
 G11 - G26   -0.00197 0.146 48  -0.013  1.0000
 G11 - G35   -0.08305 0.146 48  -0.569  0.9926
 G11 - G46   -0.00197 0.146 48  -0.013  1.0000
 G11 - G6    -0.06238 0.146 48  -0.427  0.9981
 G24.1 - G26  0.00000 0.146 48   0.000  1.0000
 G24.1 - G35 -0.08108 0.146 48  -0.555  0.9934
 G24.1 - G46  0.00000 0.146 48   0.000  1.0000
 G24.1 - G6  -0.06041 0.146 48  -0.414  0.9984
 G26 - G35   -0.08108 0.146 48  -0.555  0.9934
 G26 - G46    0.00000 0.146 48   0.000  1.0000
 G26 - G6    -0.06041 0.146 48  -0.414  0.9984
 G35 - G46    0.08108 0.146 48   0.555  0.9934
 G35 - G6     0.02067 0.146 48   0.142  1.0000
 G46 - G6    -0.06041 0.146 48  -0.414  0.9984

time.fac = 8, infstat = non:
 contrast     estimate    SE df t.ratio p.value
 G11 - G24.1  0.00000 0.146 48   0.000  1.0000
 G11 - G26    0.00000 0.146 48   0.000  1.0000
 G11 - G35    0.00000 0.146 48   0.000  1.0000
 G11 - G46    0.00000 0.146 48   0.000  1.0000
 G11 - G6     0.00000 0.146 48   0.000  1.0000
 G24.1 - G26  0.00000 0.146 48   0.000  1.0000
 G24.1 - G35  0.00000 0.146 48   0.000  1.0000
 G24.1 - G46  0.00000 0.146 48   0.000  1.0000
 G24.1 - G6   0.00000 0.146 48   0.000  1.0000
 G26 - G35    0.00000 0.146 48   0.000  1.0000
 G26 - G46    0.00000 0.146 48   0.000  1.0000
 G26 - G6     0.00000 0.146 48   0.000  1.0000
 G35 - G46    0.00000 0.146 48   0.000  1.0000
 G35 - G6     0.00000 0.146 48   0.000  1.0000
 G46 - G6     0.00000 0.146 48   0.000  1.0000
```

#### Fig. 5

```
time.fac = 25, infstat = inf:
 contrast    estimate    SE df t.ratio p.value
 G11 - G24.1  0.73889 0.146 48   5.059  0.0001
 G11 - G26    0.32990 0.146 48   2.259  0.2310
 G11 - G35    0.86746 0.146 48   5.940  <.0001
 G11 - G46    1.52235 0.146 48  10.424  <.0001
 G11 - G6     0.43878 0.146 48   3.004  0.0457
 G24.1 - G26 -0.40899 0.146 48  -2.800  0.0746
 G24.1 - G35  0.12857 0.146 48   0.880  0.9494
 G24.1 - G46  0.78346 0.146 48   5.365  <.0001
 G24.1 - G6  -0.30011 0.146 48  -2.055  0.3278
 G26 - G35    0.53756 0.146 48   3.681  0.0073
 G26 - G46    1.19245 0.146 48   8.165  <.0001
 G26 - G6     0.10888 0.146 48   0.746  0.9750
 G35 - G46    0.65489 0.146 48   4.484  0.0006
 G35 - G6    -0.42868 0.146 48  -2.935  0.0542
 G46 - G6    -1.08357 0.146 48  -7.420  <.0001
```

**Model 8 min EXP INF normalised (lower section of panel)**

```
Call:
lm(formula = log10fch ~ time.fac * gene)

Residuals:
     Min       1Q   Median       3Q      Max
-0.67700 -0.17978  0.03291  0.19143  0.59367

Coefficients:
                      Estimate Std. Error t value Pr(>|t|)
(Intercept)          -0.790912   0.199504  -3.964 0.000172 ***
time.fac10            1.741600   0.282141   6.173 3.56e-08 ***
time.fac25            2.642040   0.282141   9.364 4.40e-14 ***
time.fac30            0.881239   0.282141   3.123 0.002575 **
time.fac6             1.284381   0.282141   4.552 2.11e-05 ***
time.fac8             0.790912   0.282141   2.803 0.006497 **
geneG24.1             0.610862   0.282141   2.165 0.033696 *
geneG26               0.467275   0.282141   1.656 0.102039
geneG35              -0.213177   0.282141  -0.756 0.452373
geneG46              -0.702500   0.282141  -2.490 0.015088 *
geneG6                0.262351   0.282141   0.930 0.355553
time.fac10:geneG24.1  0.008472   0.399008   0.021 0.983118
time.fac25:geneG24.1 -0.469869   0.399008  -1.178 0.242837
time.fac30:geneG24.1  0.047558   0.399008   0.119 0.905456
time.fac6:geneG24.1   0.018989   0.399008   0.048 0.962175
time.fac8:geneG24.1  -0.610862   0.399008  -1.531 0.130164
time.fac10:geneG26   -0.061509   0.399008  -0.154 0.877918
time.fac25:geneG26   -0.435550   0.399008  -1.092 0.278656
time.fac30:geneG26    0.449678   0.399008   1.127 0.263489
time.fac6:geneG26    -0.099607   0.399008  -0.250 0.803579
time.fac8:geneG26    -0.467275   0.399008  -1.171 0.245423
time.fac10:geneG35   -0.480365   0.399008  -1.204 0.232572
time.fac25:geneG35   -1.423253   0.399008  -3.567 0.000647 ***
time.fac30:geneG35    1.402295   0.399008   3.514 0.000766 ***
time.fac6:geneG35    -0.432022   0.399008  -1.083 0.282537
time.fac8:geneG35     0.213177   0.399008   0.534 0.594802
time.fac10:geneG46   -0.540560   0.399008  -1.355 0.179730
time.fac25:geneG46   -1.816262   0.399008  -4.552 2.11e-05 ***
time.fac30:geneG46    0.662508   0.399008   1.660 0.101186
time.fac6:geneG46    -0.531132   0.399008  -1.331 0.187346
time.fac8:geneG46     0.702500   0.399008   1.761 0.082550 .
time.fac10:geneG6    -0.238629   0.399008  -0.598 0.551679
time.fac25:geneG6    -0.300892   0.399008  -0.754 0.453248
time.fac30:geneG6    -0.050094   0.399008  -0.126 0.900441
time.fac6:geneG6     -0.258793   0.399008  -0.649 0.518666
time.fac8:geneG6     -0.262351   0.399008  -0.658 0.512951
---
Signif. codes:  0 ‘***’ 0.001 ‘**’ 0.01 ‘*’ 0.05 ‘.’ 0.1 ‘ ’ 1

Residual standard error: 0.3456 on 72 degrees of freedom
Multiple R-squared:  0.9037,  Adjusted R-squared:  0.8569
F-statistic:  19.3 on 35 and 72 DF,  p-value: < 2.2e-16
```

**Analysis of Variance Table**

```
Response: log10fch
              Df Sum Sq Mean Sq F value    Pr(>F)
time.fac       5 39.862  7.9723 66.7667 < 2.2e-16 ***
gene           5 24.011  4.8022 40.2173 < 2.2e-16 ***
time.fac:gene 25 16.794  0.6718  5.6259 4.194e-09 ***
Residuals     72  8.597  0.1194
---
Signif. codes:  0 ‘***’ 0.001 ‘**’ 0.01 ‘*’ 0.05 ‘.’ 0.1 ‘ ’ 1
```

```
time.fac = 8, infstat = inf:
 contrast    estimate    SE df t.ratio p.value
 G11 - G24.1  0.87988 0.146 48   6.025  <.0001
 G11 - G26    0.36162 0.146 48   2.476  0.1517
 G11 - G35   -0.76897 0.146 48  -5.265  <.0001
 G11 - G46   -0.99642 0.146 48  -6.823  <.0001
 G11 - G6     0.40024 0.146 48   2.741  0.0857
 G24.1 - G26 -0.51826 0.146 48  -3.549  0.0107
 G24.1 - G35 -1.64886 0.146 48 -11.290  <.0001
 G24.1 - G46 -1.87630 0.146 48 -12.848  <.0001
 G24.1 - G6  -0.47964 0.146 48  -3.284  0.0222
 G26 - G35   -1.13060 0.146 48  -7.742  <.0001
 G26 - G46   -1.35804 0.146 48  -9.299  <.0001
 G26 - G6     0.03861 0.146 48   0.264  0.9998
 G35 - G46   -0.22744 0.146 48  -1.557  0.6297
 G35 - G6     1.16921 0.146 48   8.006  <.0001
 G46 - G6     1.39666 0.146 48   9.563  <.0001

P value adjustment: tukey method for comparing a family of 6
estimates
```

**Simultaneous Tests for General Linear Hypotheses**

```
gene = G11:
 contrast  estimate    SE df t.ratio p.value
   1 - 10   -1.7416 0.282 72  -6.173  <.0001
   1 - 25   -2.6420 0.282 72  -9.364  <.0001
   1 - 30   -0.8812 0.282 72  -3.123  0.0297
   1 - 6    -1.2844 0.282 72  -4.552  0.0003
   1 - 8    -0.7909 0.282 72  -2.803  0.0684
   10 - 25  -0.9004 0.282 72  -3.191  0.0246
   10 - 30   0.8604 0.282 72   3.049  0.0363
   10 - 6    0.4572 0.282 72   1.621  0.5879
   10 - 8    0.9507 0.282 72   3.370  0.0148
   25 - 30   1.7608 0.282 72   6.241  <.0001
   25 - 6    1.3577 0.282 72   4.812  0.0001
   25 - 8    1.8511 0.282 72   6.561  <.0001
   30 - 6   -0.4031 0.282 72  -1.429  0.7095
   30 - 8    0.0903 0.282 72   0.320  0.9995
   6 - 8     0.4935 0.282 72   1.749  0.5047

gene = G24.1:
 contrast  estimate    SE df t.ratio p.value
   1 - 10   -1.7501 0.282 72  -6.203  <.0001
   1 - 25   -2.1722 0.282 72  -7.699  <.0001
   1 - 30   -0.9288 0.282 72  -3.292  0.0185
   1 - 6    -1.3034 0.282 72  -4.620  0.0002
   1 - 8    -0.1801 0.282 72  -0.638  0.9877
   10 - 25  -0.4221 0.282 72  -1.496  0.6679
   10 - 30   0.8213 0.282 72   2.911  0.0522
   10 - 6    0.4467 0.282 72   1.583  0.6121
   10 - 8    1.5700 0.282 72   5.565  <.0001
   25 - 30   1.2434 0.282 72   4.407  0.0005
   25 - 6    0.8688 0.282 72   3.079  0.0335
   25 - 8    1.9921 0.282 72   7.061  <.0001
   30 - 6   -0.3746 0.282 72  -1.328  0.7688
   30 - 8    0.7487 0.282 72   2.654  0.0977
   6 - 8     1.1233 0.282 72   3.981  0.0022

gene = G26:
 contrast  estimate    SE df t.ratio p.value
   1 - 10   -1.6801 0.282 72  -5.955  <.0001
   1 - 25   -2.2065 0.282 72  -7.821  <.0001
   1 - 30   -1.3309 0.282 72  -4.717  0.0002
   1 - 6    -1.1848 0.282 72  -4.199  0.0010
   1 - 8    -0.3236 0.282 72  -1.147  0.8598
   10 - 25  -0.5264 0.282 72  -1.866  0.4314
   10 - 30   0.3492 0.282 72   1.238  0.8169
   10 - 6    0.4953 0.282 72   1.756  0.5005
   10 - 8    1.3565 0.282 72   4.808  0.0001
   25 - 30   0.8756 0.282 72   3.103  0.0314
   25 - 6    1.0217 0.282 72   3.621  0.0069
   25 - 8    1.8829 0.282 72   6.673  <.0001
   30 - 6    0.1461 0.282 72   0.518  0.9953
   30 - 8    1.0073 0.282 72   3.570  0.0081
   6 - 8     0.8611 0.282 72   3.052  0.0361

gene = G35:
 contrast  estimate    SE df t.ratio p.value
   1 - 10   -1.2612 0.282 72  -4.470  0.0004
   1 - 25   -1.2188 0.282 72  -4.320  0.0007
   1 - 30   -2.2835 0.282 72  -8.094  <.0001
   1 - 6    -0.8524 0.282 72  -3.021  0.0392
   1 - 8    -1.0041 0.282 72  -3.559  0.0084
   10 - 25   0.0424 0.282 72   0.150  1.0000
   10 - 30  -1.0223 0.282 72  -3.623  0.0069
   10 - 6    0.4089 0.282 72   1.449  0.6971
   10 - 8    0.2571 0.282 72   0.911  0.9424
   25 - 30  -1.0647 0.282 72  -3.774  0.0043
   25 - 6    0.3664 0.282 72   1.299  0.7848
   25 - 8    0.2147 0.282 72   0.761  0.9731
   30 - 6    1.4312 0.282 72   5.073  <.0001
   30 - 8    1.2794 0.282 72   4.535  0.0003
   6 - 8    -0.1517 0.282 72  -0.538  0.9944
```

#### Fig. 5

```
gene = G46:
 contrast  estimate    SE df t.ratio p.value
   1 - 10   -1.2010 0.282 72  -4.257  0.0008
   1 - 25   -0.8258 0.282 72  -2.927  0.0501
   1 - 30   -1.5437 0.282 72  -5.472  <.0001
   1 - 6    -0.7532 0.282 72  -2.670  0.0942
   1 - 8    -1.4934 0.282 72  -5.293  <.0001
   10 - 25   0.3753 0.282 72   1.330  0.7674
   10 - 30  -0.3427 0.282 72  -1.215  0.8283
   10 - 6    0.4478 0.282 72   1.587  0.6096
   10 - 8   -0.2924 0.282 72  -1.036  0.9041
   25 - 30  -0.7180 0.282 72  -2.545  0.1250
   25 - 6    0.0725 0.282 72   0.257  0.9998
   25 - 8   -0.6676 0.282 72  -2.366  0.1821
   30 - 6    0.7905 0.282 72   2.802  0.0686
   30 - 8    0.0503 0.282 72   0.178  1.0000
   6 - 8    -0.7402 0.282 72  -2.623  0.1048

gene = G6:
 contrast  estimate    SE df t.ratio p.value
   1 - 10   -1.5030 0.282 72  -5.327  <.0001
   1 - 25   -2.3411 0.282 72  -8.298  <.0001
   1 - 30   -0.8311 0.282 72  -2.946  0.0477
   1 - 6    -1.0256 0.282 72  -3.635  0.0066
   1 - 8    -0.5286 0.282 72  -1.873  0.4267
   10 - 25  -0.8382 0.282 72  -2.971  0.0447
   10 - 30   0.6718 0.282 72   2.381  0.1767
   10 - 6    0.4774 0.282 72   1.692  0.5415
   10 - 8    0.9744 0.282 72   3.454  0.0115
   25 - 30   1.5100 0.282 72   5.352  <.0001
   25 - 6    1.3156 0.282 72   4.663  0.0002
   25 - 8    1.8126 0.282 72   6.424  <.0001
   30 - 6   -0.1944 0.282 72  -0.689  0.9826
   30 - 8    0.3026 0.282 72   1.072  0.8907
   6 - 8     0.4970 0.282 72   1.762  0.4966

P value adjustment: tukey method for comparing a family of 6
estimates

------

time.fac = 1:
 contrast    estimate    SE df t.ratio p.value
 G11 - G24.1 -0.61086 0.282 72  -2.165  0.2669
 G11 - G26   -0.46727 0.282 72  -1.656  0.5648
 G11 - G35    0.21318 0.282 72   0.756  0.9739
 G11 - G46    0.70250 0.282 72   2.490  0.1408
 G11 - G6    -0.26235 0.282 72  -0.930  0.9375
 G24.1 - G26  0.14359 0.282 72   0.509  0.9957
 G24.1 - G35  0.82404 0.282 72   2.921  0.0509
 G24.1 - G46  1.31336 0.282 72   4.655  0.0002
 G24.1 - G6   0.34851 0.282 72   1.235  0.8181
 G26 - G35    0.68045 0.282 72   2.412  0.1660
 G26 - G46    1.16978 0.282 72   4.146  0.0012
 G26 - G6     0.20492 0.282 72   0.726  0.9781
 G35 - G46    0.48932 0.282 72   1.734  0.5142
 G35 - G6    -0.47553 0.282 72  -1.685  0.5458
 G46 - G6    -0.96485 0.282 72  -3.420  0.0128

time.fac = 10:
 contrast    estimate    SE df t.ratio p.value
 G11 - G24.1 -0.61933 0.282 72  -2.195  0.2528
 G11 - G26   -0.40577 0.282 72  -1.438  0.7038
 G11 - G35    0.69354 0.282 72   2.458  0.1507
 G11 - G46    1.24306 0.282 72   4.406  0.0005
 G11 - G6    -0.02372 0.282 72  -0.084  1.0000
 G24.1 - G26  0.21357 0.282 72   0.757  0.9737
 G24.1 - G35  1.31288 0.282 72   4.653  0.0002
 G24.1 - G46  1.86240 0.282 72   6.601  <.0001
 G24.1 - G6   0.59561 0.282 72   2.111  0.2934
 G26 - G35    1.09931 0.282 72   3.896  0.0029
 G26 - G46    1.64883 0.282 72   5.844  <.0001
 G26 - G6     0.38204 0.282 72   1.354  0.7538
 G35 - G46    0.54952 0.282 72   1.948  0.3824
 G35 - G6    -0.71726 0.282 72  -2.542  0.1257
 G46 - G6    -1.26678 0.282 72  -4.490  0.0004

time.fac = 25:
 contrast    estimate    SE df t.ratio p.value
 G11 - G24.1 -0.14099 0.282 72  -0.500  0.9960
 G11 - G26   -0.03172 0.282 72  -0.112  1.0000
 G11 - G35    1.63643 0.282 72   5.800  <.0001
 G11 - G46    2.51876 0.282 72   8.927  <.0001
 G11 - G6     0.03854 0.282 72   0.137  1.0000
 G24.1 - G26  0.10927 0.282 72   0.387  0.9988
 G24.1 - G35  1.77742 0.282 72   6.300  <.0001
 G24.1 - G46  2.65976 0.282 72   9.427  <.0001
 G24.1 - G6   0.17953 0.282 72   0.636  0.9879
 G26 - G35    1.66815 0.282 72   5.912  <.0001
 G26 - G46    2.55049 0.282 72   9.040  <.0001
 G26 - G6     0.07027 0.282 72   0.249  0.9999
 G35 - G46    0.88233 0.282 72   3.127  0.0294
 G35 - G6    -1.59789 0.282 72  -5.663  <.0001
 G46 - G6    -2.48022 0.282 72  -8.791  <.0001

time.fac = 30:
 contrast    estimate    SE df t.ratio p.value
 G11 - G24.1 -0.65842 0.282 72  -2.334  0.1944
 G11 - G26   -0.91695 0.282 72  -3.250  0.0209
 G11 - G35   -1.18912 0.282 72  -4.215  0.0010
 G11 - G46    0.03999 0.282 72   0.142  1.0000
 G11 - G6    -0.21226 0.282 72  -0.752  0.9744
 G24.1 - G26 -0.25853 0.282 72  -0.916  0.9411
 G24.1 - G35 -0.53070 0.282 72  -1.881  0.4221
 G24.1 - G46  0.69841 0.282 72   2.475  0.1453
 G24.1 - G6   0.44616 0.282 72   1.581  0.6133
 G26 - G35   -0.27217 0.282 72  -0.965  0.9275
 G26 - G46    0.95695 0.282 72   3.392  0.0139
 G26 - G6     0.70470 0.282 72   2.498  0.1385
 G35 - G46    1.22911 0.282 72   4.356  0.0006
 G35 - G6     0.97686 0.282 72   3.462  0.0112
 G46 - G6    -0.25225 0.282 72  -0.894  0.9467

time.fac = 6:
 contrast    estimate    SE df t.ratio p.value
 G11 - G24.1 -0.62985 0.282 72  -2.232  0.2361
 G11 - G26   -0.36767 0.282 72  -1.303  0.7824
 G11 - G35    0.64520 0.282 72   2.287  0.2130
 G11 - G46    1.23363 0.282 72   4.372  0.0006
 G11 - G6    -0.00356 0.282 72  -0.013  1.0000
 G24.1 - G26  0.26218 0.282 72   0.929  0.9376
 G24.1 - G35  1.27505 0.282 72   4.519  0.0003
 G24.1 - G46  1.86348 0.282 72   6.605  <.0001
 G24.1 - G6   0.62629 0.282 72   2.220  0.2416
 G26 - G35    1.01287 0.282 72   3.590  0.0076
 G26 - G46    1.60130 0.282 72   5.676  <.0001
 G26 - G6     0.36411 0.282 72   1.291  0.7892
 G35 - G46    0.58843 0.282 72   2.086  0.3064
 G35 - G6    -0.64876 0.282 72  -2.299  0.2078
 G46 - G6    -1.23719 0.282 72  -4.385  0.0005

time.fac = 8:
 contrast    estimate    SE df t.ratio p.value
 G11 - G24.1  0.00000 0.282 72   0.000  1.0000
 G11 - G26    0.00000 0.282 72   0.000  1.0000
 G11 - G35    0.00000 0.282 72   0.000  1.0000
 G11 - G46    0.00000 0.282 72   0.000  1.0000
 G11 - G6     0.00000 0.282 72   0.000  1.0000
 G24.1 - G26  0.00000 0.282 72   0.000  1.0000
 G24.1 - G35  0.00000 0.282 72   0.000  1.0000
 G24.1 - G46  0.00000 0.282 72   0.000  1.0000
 G24.1 - G6   0.00000 0.282 72   0.000  1.0000
 G26 - G35    0.00000 0.282 72   0.000  1.0000
 G26 - G46    0.00000 0.282 72   0.000  1.0000
 G26 - G6     0.00000 0.282 72   0.000  1.0000
 G35 - G46    0.00000 0.282 72   0.000  1.0000
 G35 - G6     0.00000 0.282 72   0.000  1.0000
 G46 - G6     0.00000 0.282 72   0.000  1.0000

P value adjustment: tukey method for comparing a family of 6
estimates
```

==============================================================

### Figure 6

#### *Fig. 6b*

#### Growth/decline rates at time of infection (**Table S3**)

**Model (Table S3)**

```
Call:
lm(formula = gdecrt ~ time.sup)

Residuals:
       Min         1Q     Median         3Q        Max
-0.0044258 -0.0026378 -0.0000919  0.0032270  0.0051431

Coefficients:
               Estimate Std. Error t value Pr(>|t|)
(Intercept)   -0.002490   0.002248  -1.108 0.293892
time.sup1h    -0.012147   0.003178  -3.822 0.003364 **
time.sup4h    -0.015018   0.003178  -4.725 0.000810 ***
time.sup8h    -0.015345   0.003178  -4.828 0.000694 ***
time.supnon   -0.006717   0.003178  -2.113 0.060696 .
---
Signif. codes:  0 ‘***’ 0.001 ‘**’ 0.01 ‘*’ 0.05 ‘.’ 0.1 ‘ ’ 1

Residual standard error: 0.003893 on 10 degrees of freedom
Multiple R-squared:  0.7699,  Adjusted R-squared:  0.6779
F-statistic: 8.365 on 4 and 10 DF,  p-value: 0.003128
```

**Analysis of Variance Table**

```
Response:  gdecrt
            Df     Sum Sq    Mean Sq F value   Pr(>F)
time.sup     4 0.00050708 1.2677e-04  8.3652 0.003128 **
Residuals   10 0.00015154 1.5154e-05
---
Signif. codes:  0 ‘***’ 0.001 ‘**’ 0.01 ‘*’ 0.05 ‘.’ 0.1 ‘ ’ 1
```

**Simultaneous Tests for General Linear Hypotheses**

```
 contrast   estimate      SE df t.ratio p.value
 0h - 1h    0.012147 0.00318 10   3.822  0.0220
 0h - 4h    0.015018 0.00318 10   4.725  0.0056
 0h - 8h    0.015345 0.00318 10   4.828  0.0049
 0h - non   0.006717 0.00318 10   2.113  0.2860
 1h - 4h    0.002871 0.00318 10   0.903  0.8893
 1h - 8h    0.003198 0.00318 10   1.006  0.8469
 1h - non  -0.005430 0.00318 10  -1.708  0.4706
 4h - 8h    0.000327 0.00318 10   0.103  1.0000
 4h - non  -0.008301 0.00318 10  -2.612  0.1412
 8h - non  -0.008628 0.00318 10  -2.715  0.1211

P value adjustment: tukey method for comparing a family of 5
estimate
```

#### Fig. 6

#### Time to lysis (**Table S3**)

Not determined – rate already negative from start

#### Maximum decline rates & times at max decline rates (**Table S3**)

- break point 1 for non-infected samples was chosen to avoid artefacts in OD traces immediately following infection which would artificially inflate lysis rates
- second break-points for all non-infected samples capped at 1400 to avoid artefacts introduced due to condensation and clumping at late times during stationary phase
- **only** second period lysis rates were analysed for non-infected samples
- second break points for infected samples were chosen to separate lysis periods from periods of new growth

**Breakpoints for lysis rate calculation**

| infection status | supplementation time (h) | infection time (h) | bp 1 (h) | bp 2 (h) | replicate |
|---|---|---|---|---|---|
| non-infected | non | 11.0 | 13.5 | 23.3 | 1 |
| non-infected | non | 11.0 | 12.7 | 23.3 | 2 |
| non-infected | non | 11.0 | 14.8 | 23.3 | 3 |
| non-infected | 0 | 11.0 | 13.3 | 23.3 | 1 |
| non-infected | 0 | 11.0 | 11.9 | 23.3 | 2 |
| non-infected | 0 | 11.0 | 14.0 | 23.3 | 3 |
| non-infected | 1 | 11.0 | 13.7 | 23.3 | 1 |
| non-infected | 1 | 11.0 | 14.0 | 23.3 | 2 |
| non-infected | 1 | 11.0 | 14.4 | 23.3 | 3 |
| non-infected | 4 | 11.0 | 13.0 | 23.3 | 1 |
| non-infected | 4 | 11.0 | 14.0 | 23.3 | 2 |
| non-infected | 4 | 11.0 | 13.5 | 23.3 | 3 |
| non-infected | 8 | 11.0 | 13.8 | 23.3 | 1 |
| non-infected | 8 | 11.0 | 14.8 | 23.3 | 2 |
| non-infected | 8 | 11.0 | 14.2 | 23.3 | 3 |
| infected | non | 11.0 | 21.0 | 27.8 | 1 |
| infected | non | 11.0 | 19.9 | 27.8 | 2 |
| infected | non | 11.0 | 19.6 | 27.5 | 3 |
| infected | 0 | 11.0 | 18.6 | 24.7 | 1 |
| infected | 0 | 11.0 | 17.1 | 29.1 | 2 |
| infected | 0 | 11.0 | 18.3 | 24.5 | 3 |
| infected | 1 | 11.0 | 17.7 | 23.8 | 1 |
| infected | 1 | 11.0 | 17.8 | 25.7 | 2 |
| infected | 1 | 11.0 | 17.5 | 24.1 | 3 |
| infected | 4 | 11.0 | 19.1 | 24.8 | 1 |
| infected | 4 | 11.0 | 19.3 | 24.4 | 2 |
| infected | 4 | 11.0 | 19.2 | 27.9 | 3 |
| infected | 8 | 11.0 | 20.0 | 24.5 | 1 |
| infected | 8 | 11.0 | 18.1 | 26.0 | 2 |
| infected | 8 | 11.0 | 17.6 | 26.3 | 3 |

* periods noted in Table S3 (1-2) correspond to time periods between the above listed breakpoints

#### Fig. 6

**Model (Max decline rate)**
**NOTE:** time of supplementation and period of lysis were combined here into a single variable to compare directly between the different recorded rates

```
Call:
glm(formula = (mxdecrt * -1) ~ ISSP, family = gaussian(link =
"inverse"))
Deviance Residuals:
      Min         1Q     Median         3Q        Max
-0.036805  -0.001693   0.000083   0.004270   0.027204

Coefficients:
             Estimate Std. Error t value Pr(>|t|)
(Intercept)   13.7011     1.4412   9.506 3.01e-09 ***
ISSPinf0h2   -12.2110     1.4413  -8.472 2.25e-08 ***
ISSPinf1h1     2.4042     2.4583   0.978 0.338702
ISSPinf1h2   -12.0521     1.4414  -8.361 2.81e-08 ***
ISSPinf4h1    -5.9381     1.5137  -3.923 0.000728 ***
ISSPinf4h2   -12.0124     1.4414  -8.334 2.98e-08 ***
ISSPinf8h1    -0.2475     2.0021  -0.124 0.902740
ISSPinf8h2   -12.2566     1.4413  -8.504 2.11e-08 ***
ISSPinfnon1   -5.8925     1.5154  -3.888 0.000791 ***
ISSPinfnon2  -10.8683     1.4426  -7.534 1.58e-07 ***
ISSPnonnon2    1.3244     2.2543   0.588 0.562843
---
Signif. codes:  0 ‘***’ 0.001 ‘**’ 0.01 ‘*’ 0.05 ‘.’ 0.1 ‘ ’ 1
(Dispersion parameter for gaussian family taken to be
0.0001768404)
    Null deviance: 2.2362313  on 32  degrees of freedom
Residual deviance: 0.0038905  on 22  degrees of freedom
AIC: -180.86
Number of Fisher Scoring iterations: 5
```

**Analysis of Deviance Table**

```
Model: gaussian, link: inverse
Response: (mxdecrt * -1)
Terms added sequentially (first to last)

      Df Deviance Resid. Df Resid. Dev  Pr(>Chi)
NULL                     32    2.23623
ISSP  10   2.2323        22    0.00389 < 2.2e-16 ***
---
Signif. codes:  0 ‘***’ 0.001 ‘**’ 0.01 ‘*’ 0.05 ‘.’ 0.1 ‘ ’ 1
```

**Simultaneous Tests for General Linear Hypotheses**

```
 contrast             estimate     SE df t.ratio p.value
 inf0h1 - inf0h2     -0.598107 0.0109 22 -55.085  <.0001
 inf0h1 - inf1h1      0.010896 0.0109 22   1.003  0.9933
 inf0h1 - inf1h2     -0.533456 0.0109 22 -49.131  <.0001
 inf0h1 - inf4h1     -0.055830 0.0109 22  -5.142  0.0015
 inf0h1 - inf4h2     -0.519192 0.0109 22 -47.817  <.0001
 inf0h1 - inf8h1     -0.001343 0.0109 22  -0.124  1.0000
 inf0h1 - inf8h2     -0.619287 0.0109 22 -57.036  <.0001
 inf0h1 - infnon1    -0.055077 0.0109 22  -5.073  0.0017
 inf0h1 - infnon2    -0.280017 0.0109 22 -25.789  <.0001
 inf0h1 - nonnon2     0.006433 0.0109 22   0.593  0.9999
 inf0h2 - inf1h1      0.609003 0.0109 22  56.089  <.0001
 inf0h2 - inf1h2      0.064651 0.0109 22   5.954  0.0002
 inf0h2 - inf4h1      0.542277 0.0109 22  49.943  <.0001
 inf0h2 - inf4h2      0.078915 0.0109 22   7.268  <.0001
 inf0h2 - inf8h1      0.596764 0.0109 22  54.961  <.0001
 inf0h2 - inf8h2     -0.021180 0.0109 22  -1.951  0.6810
 inf0h2 - infnon1     0.543030 0.0109 22  50.012  <.0001
 inf0h2 - infnon2     0.318090 0.0109 22  29.296  <.0001
 inf0h2 - nonnon2     0.604540 0.0109 22  55.678  <.0001
 inf1h1 - inf1h2     -0.544351 0.0109 22 -50.134  <.0001
 inf1h1 - inf4h1     -0.066725 0.0109 22  -6.145  0.0001
 inf1h1 - inf4h2     -0.530088 0.0109 22 -48.821  <.0001
 inf1h1 - inf8h1     -0.012238 0.0109 22  -1.127  0.9843
 inf1h1 - inf8h2     -0.630183 0.0109 22 -58.039  <.0001
 inf1h1 - infnon1    -0.065973 0.0109 22  -6.076  0.0002
 inf1h1 - infnon2    -0.290912 0.0109 22 -26.793  <.0001
 inf1h1 - nonnon2    -0.004462 0.0109 22  -0.411  1.0000
 inf1h2 - inf4h1      0.477626 0.0109 22  43.989  <.0001
 inf1h2 - inf4h2      0.014264 0.0109 22   1.314  0.9564
 inf1h2 - inf8h1      0.532113 0.0109 22  49.007  <.0001
 inf1h2 - inf8h2     -0.085831 0.0109 22  -7.905  <.0001
 inf1h2 - infnon1     0.478379 0.0109 22  44.058  <.0001
 inf1h2 - infnon2     0.253439 0.0109 22  23.341  <.0001
 inf1h2 - nonnon2     0.539889 0.0109 22  49.723  <.0001
 inf4h1 - inf4h2     -0.463362 0.0109 22 -42.675  <.0001
 inf4h1 - inf8h1      0.054487 0.0109 22   5.018  0.0020
 inf4h1 - inf8h2     -0.563457 0.0109 22 -51.894  <.0001
 inf4h1 - infnon1     0.000753 0.0109 22   0.069  1.0000
 inf4h1 - infnon2    -0.224187 0.0109 22 -20.647  <.0001
 inf4h1 - nonnon2     0.062263 0.0109 22   5.734  0.0004
 inf4h2 - inf8h1      0.517849 0.0109 22  47.693  <.0001
 inf4h2 - inf8h2     -0.100095 0.0109 22  -9.219  <.0001
 inf4h2 - infnon1     0.464115 0.0109 22  42.745  <.0001
 inf4h2 - infnon2     0.239175 0.0109 22  22.028  <.0001
 inf4h2 - nonnon2     0.525625 0.0109 22  48.410  <.0001
 inf8h1 - inf8h2     -0.617945 0.0109 22 -56.912  <.0001
 inf8h1 - infnon1    -0.053734 0.0109 22  -4.949  0.0023
 inf8h1 - infnon2    -0.278674 0.0109 22 -25.666  <.0001
 inf8h1 - nonnon2     0.007776 0.0109 22   0.716  0.9996
 inf8h2 - infnon1     0.564210 0.0109 22  51.963  <.0001
 inf8h2 - infnon2     0.339270 0.0109 22  31.246  <.0001
 inf8h2 - nonnon2     0.625721 0.0109 22  57.628  <.0001
 infnon1 - infnon2   -0.224940 0.0109 22 -20.717  <.0001
 infnon1 - nonnon2    0.061511 0.0109 22   5.665  0.0004
 infnon2 - nonnon2    0.286450 0.0109 22  26.382  <.0001
P value adjustment: tukey method for comparing a family of 11
estimates
```

**Model (Time at max decline rate)**
**NOTE:** data were box-cox transformed before analysis

```
Call:
lm(formula = (tmxdec^5.050505 - 1)/5.050505 ~ ISSP)

Residuals:
    Min      1Q  Median      3Q     Max
-292923       0       0    8551  467401

Coefficients:
            Estimate Std. Error t value Pr(>|t|)
(Intercept)   328916      91274   3.604  0.00158 **
ISSPinf0h2   1164147     129080   9.019 7.63e-09 ***
ISSPinf1h1    -47918     129080  -0.371  0.71402
ISSPinf1h2   1092874     129080   8.467 2.27e-08 ***
ISSPinf4h1    257061     129080   1.991  0.05899 .
ISSPinf4h2    863911     129080   6.693 9.97e-07 ***
ISSPinf8h1    -59223     129080  -0.459  0.65088
ISSPinf8h2   1058022     129080   8.197 3.94e-08 ***
ISSPinfnon1   292187     129080   2.264  0.03381 *
ISSPinfnon2  2185304     129080  16.930 4.20e-14 ***
ISSPnonnon2  3026721     129080  23.448  < 2e-16 ***
---
Signif. codes:  0 ‘***’ 0.001 ‘**’ 0.01 ‘*’ 0.05 ‘.’ 0.1 ‘ ’ 1

Residual standard error: 158100 on 22 degrees of freedom
Multiple R-squared:  0.9815,  Adjusted R-squared:  0.9731
F-statistic: 116.7 on 10 and 22 DF,  p-value: < 2.2e-16
```

**Analysis of Variance Table**

```
Response: (tmxdec^5.050505 - 1)/5.050505
          Df     Sum Sq    Mean Sq F value    Pr(>F)
ISSP       10 2.9158e+13 2.9158e+12  116.67 < 2.2e-16 ***
Residuals 22 5.4984e+11 2.4993e+10
---
Signif. codes:  0 ‘***’ 0.001 ‘**’ 0.01 ‘*’ 0.05 ‘.’ 0.1 ‘ ’ 1
```

Fig. 6

```
Simultaneous Tests for General Linear Hypotheses
 contrast           estimate     SE df t.ratio p.value
 inf0h1 - inf0h2    -1164147 129081 22  -9.019  <.0001
 inf0h1 - inf1h1       47918 129081 22   0.371  1.0000
 inf0h1 - inf1h2    -1092874 129081 22  -8.467  <.0001
 inf0h1 - inf4h1     -257061 129081 22  -1.991  0.6561
 inf0h1 - inf4h2     -863911 129081 22  -6.693  <.0001
 inf0h1 - inf8h1       59223 129081 22   0.459  1.0000
 inf0h1 - inf8h2    -1058022 129081 22  -8.197  <.0001
 inf0h1 - infnon1    -292187 129081 22  -2.264  0.4884
 inf0h1 - infnon2   -2185304 129081 22 -16.930  <.0001
 inf0h1 - nonnon2   -3026721 129081 22 -23.448  <.0001
 inf0h2 - inf1h1     1212065 129081 22   9.390  <.0001
 inf0h2 - inf1h2       71273 129081 22   0.552  1.0000
 inf0h2 - inf4h1      907086 129081 22   7.027  <.0001
 inf0h2 - inf4h2      300236 129081 22   2.326  0.4514
 inf0h2 - inf8h1     1223370 129081 22   9.478  <.0001
 inf0h2 - inf8h2      106125 129081 22   0.822  0.9986
 inf0h2 - infnon1     871961 129081 22   6.755  <.0001
 inf0h2 - infnon2   -1021157 129081 22  -7.911  <.0001
 inf0h2 - nonnon2   -1862574 129081 22 -14.430  <.0001
 inf1h1 - inf1h2    -1140792 129081 22  -8.838  <.0001
 inf1h1 - inf4h1     -304978 129081 22  -2.363  0.4302
 inf1h1 - inf4h2     -911828 129081 22  -7.064  <.0001
 inf1h1 - inf8h1       11305 129081 22   0.088  1.0000
 inf1h1 - inf8h2    -1105940 129081 22  -8.568  <.0001
 inf1h1 - infnon1    -340104 129081 22  -2.635  0.2896
 inf1h1 - infnon2   -2233222 129081 22 -17.301  <.0001
 inf1h1 - nonnon2   -3074638 129081 22 -23.820  <.0001
 inf1h2 - inf4h1      835813 129081 22   6.475  0.0001
 inf1h2 - inf4h2      228964 129081 22   1.774  0.7825
 inf1h2 - inf8h1     1152097 129081 22   8.925  <.0001
 inf1h2 - inf8h2       34852 129081 22   0.270  1.0000
 inf1h2 - infnon1     800688 129081 22   6.203  0.0001
 inf1h2 - infnon2   -1092430 129081 22  -8.463  <.0001
 inf1h2 - nonnon2   -1933846 129081 22 -14.982  <.0001
 inf4h1 - inf4h2     -606850 129081 22  -4.701  0.0041
 inf4h1 - inf8h1      316284 129081 22   2.450  0.3815
 inf4h1 - inf8h2     -800961 129081 22  -6.205  0.0001
 inf4h1 - infnon1     -35126 129081 22  -0.272  1.0000
 inf4h1 - infnon2   -1928243 129081 22 -14.938  <.0001
 inf4h1 - nonnon2   -2769660 129081 22 -21.457  <.0001
 inf4h2 - inf8h1      923133 129081 22   7.152  <.0001
 inf4h2 - inf8h2     -194111 129081 22  -1.504  0.9035
 inf4h2 - infnon1     571724 129081 22   4.429  0.0076
 inf4h2 - infnon2   -1321393 129081 22 -10.237  <.0001
 inf4h2 - nonnon2   -2162810 129081 22 -16.756  <.0001
 inf8h1 - inf8h2    -1117245 129081 22  -8.655  <.0001
 inf8h1 - infnon1    -351409 129081 22  -2.722  0.2516
 inf8h1 - infnon2   -2244527 129081 22 -17.389  <.0001
 inf8h1 - nonnon2   -3085943 129081 22 -23.907  <.0001
 inf8h2 - infnon1     765836 129081 22   5.933  0.0002
 inf8h2 - infnon2   -1127282 129081 22  -8.733  <.0001
 inf8h2 - nonnon2   -1968699 129081 22 -15.252  <.0001
 infnon1 - infnon2  -1893118 129081 22 -14.666  <.0001
 infnon1 - nonnon2  -2734534 129081 22 -21.185  <.0001
 infnon2 - nonnon2   -841417 129081 22  -6.519  0.0001

P value adjustment: tukey method for comparing a family of 11
estimates
```

### Duration of infection (**Table S3**)

```
Model
Call:
lm(formula = dur ~ time.sup)

Residuals:
     Min       1Q   Median       3Q      Max
-0.55556 -0.38889 -0.05556  0.30556  0.94444

Coefficients:
              Estimate Std. Error t value Pr(>|t|)
(Intercept)    14.6667     0.3315  44.247 8.36e-13 ***
time.sup1h     -0.2778     0.4688  -0.593    0.567
time.sup4h     -0.7778     0.4688  -1.659    0.128
time.sup8h     -0.2778     0.4688  -0.593    0.567
time.supnon     3.3889     0.4688   7.229 2.83e-05 ***
---
Signif. codes:  0 ‘***’ 0.001 ‘**’ 0.01 ‘*’ 0.05 ‘.’ 0.1 ‘ ’ 1

Residual standard error: 0.5741 on 10 degrees of freedom
Multiple R-squared:  0.9121,  Adjusted R-squared:  0.8769
F-statistic: 25.94 on 4 and 10 DF,  p-value: 2.921e-05
```

```
Analysis of Variance Table
Response: dur
          Df Sum Sq Mean Sq F value    Pr(>F)
time.sup   4 34.196  8.5491  25.935 2.921e-05 ***
Residuals 10  3.296  0.3296
---
Signif. codes:  0 ‘***’ 0.001 ‘**’ 0.01 ‘*’ 0.05 ‘.’ 0.1 ‘ ’ 1
```

```
Simultaneous Tests for General Linear Hypotheses
 contrast  estimate    SE df t.ratio p.value
 0h - 1h      0.278 0.469 10   0.593  0.9731
 0h - 4h      0.778 0.469 10   1.659  0.4967
 0h - 8h      0.278 0.469 10   0.593  0.9731
 0h - non    -3.389 0.469 10  -7.229  0.0002
 1h - 4h      0.500 0.469 10   1.067  0.8191
 1h - 8h      0.000 0.469 10   0.000  1.0000
 1h - non    -3.667 0.469 10  -7.822  0.0001
 4h - 8h     -0.500 0.469 10  -1.067  0.8191
 4h - non    -4.167 0.469 10  -8.888  <.0001
 8h - non    -3.667 0.469 10  -7.822  0.0001

P value adjustment: tukey method for comparing a family of 5
estimates
```

### % decrease OD (**Table S3**)

```
Model
Call:
lm(formula = percdec ~ infstat * time.sup)

Residuals:
    Min      1Q  Median      3Q     Max
-5.2100 -0.8145  0.4392  0.8716  4.0147

Coefficients:
                        Estimate Std. Error t value Pr(>|t|)
(Intercept)                4.460      1.484   3.006  0.00698 **
infstatnon                83.976      2.098  40.022  < 2e-16 ***
time.sup1h                 2.621      2.098   1.249  0.22603
time.sup4h                12.933      2.098   6.164 5.06e-06 ***
time.sup8h                26.067      2.098  12.423 7.35e-11 ***
time.supnon               44.558      2.098  21.236 3.43e-15 ***
infstatinf:time.sup1h     -2.719      2.967  -0.916  0.37041
infstatinf:time.sup4h    -12.253      2.967  -4.129  0.00052 ***
infstatinf:time.sup8h    -25.431      2.967  -8.570 3.96e-08 ***
infstatinf:time.supnon   -43.649      2.967 -14.709 3.45e-12 ***
---
Signif. codes:  0 ‘***’ 0.001 ‘**’ 0.01 ‘*’ 0.05 ‘.’ 0.1 ‘ ’ 1

Residual standard error: 2.57 on 20 degrees of freedom
Multiple R-squared:  0.9965,  Adjusted R-squared:  0.995
F-statistic: 637.6 on 9 and 20 DF,  p-value: < 2.2e-16
```

```
Analysis of Variance Table
Response: percdec
                 Df Sum Sq Mean Sq  F value    Pr(>F)
infstat           1  33834   33834 5123.204 < 2.2e-16 ***
time.sup          4   2119     530   80.201 5.045e-12 ***
infstat:time.sup  4   1945     486   73.621 1.122e-11 ***
Residuals        20    132       7
---
Signif. codes:  0 ‘***’ 0.001 ‘**’ 0.01 ‘*’ 0.05 ‘.’ 0.1 ‘ ’ 1
```

#### Fig. 6

```
Simultaneous Tests for General Linear Hypotheses
time.sup = 0h:
 contrast  estimate  SE df t.ratio p.value
 non - inf    -84.0 2.1 20 -40.022  <.0001

time.sup = 1h:
 contrast  estimate  SE df t.ratio p.value
 non - inf    -81.3 2.1 20 -38.726  <.0001

time.sup = 4h:
 contrast  estimate  SE df t.ratio p.value
 non - inf    -71.7 2.1 20 -34.182  <.0001

time.sup = 8h:
 contrast  estimate  SE df t.ratio p.value
 non - inf    -58.5 2.1 20 -27.901  <.0001

time.sup = non:
 contrast  estimate  SE df t.ratio p.value
 non - inf    -40.3 2.1 20 -19.219  <.0001

------


infstat = non:
 contrast  estimate  SE df t.ratio p.value
 0h - 1h     -2.6211 2.1 20  -1.249  0.7237
 0h - 4h    -12.9333 2.1 20  -6.164  <.0001
 0h - 8h    -26.0667 2.1 20 -12.423  <.0001
 0h - non   -44.5581 2.1 20 -21.236  <.0001
 1h - 4h    -10.3122 2.1 20  -4.915  0.0007
 1h - 8h    -23.4456 2.1 20 -11.174  <.0001
 1h - non   -41.9370 2.1 20 -19.987  <.0001
 4h - 8h    -13.1334 2.1 20  -6.259  <.0001
 4h - non   -31.6248 2.1 20 -15.072  <.0001
 8h - non   -18.4914 2.1 20  -8.813  <.0001

infstat = inf:
 contrast  estimate  SE df t.ratio p.value
 0h - 1h      0.0980 2.1 20   0.047  1.0000
 0h - 4h     -0.6801 2.1 20  -0.324  0.9974
 0h - 8h     -0.6355 2.1 20  -0.303  0.9980
 0h - non    -0.9095 2.1 20  -0.433  0.9921
 1h - 4h     -0.7781 2.1 20  -0.371  0.9956
 1h - 8h     -0.7335 2.1 20  -0.350  0.9965
 1h - non    -1.0076 2.1 20  -0.480  0.9883
 4h - 8h      0.0446 2.1 20   0.021  1.0000
 4h - non    -0.2294 2.1 20  -0.109  1.0000
 8h - non    -0.2740 2.1 20  -0.131  0.9999

P value adjustment: tukey method for comparing a family of 5
estimates
```

#### Time to 50% max OD loss (**Table S3**)

```
Model (Table S3)
Call:
lm(formula = time50 ~ time.sup)

Residuals:
     Min       1Q   Median       3Q      Max
-0.61111 -0.30556  0.05556  0.19444  0.72222

Coefficients:
                Estimate Std. Error t value Pr(>|t|)
(Intercept)      10.8889     0.2699  40.346 2.09e-12 ***
time.sup1h       -0.2222     0.3817  -0.582   0.5733
time.sup4h       -0.7778     0.3817  -2.038   0.0689 .
time.sup8h       -0.3333     0.3817  -0.873   0.4030
time.supnon       0.8889     0.3817   2.329   0.0421 *
---
Signif. codes:  0 ‘***’ 0.001 ‘**’ 0.01 ‘*’ 0.05 ‘.’ 0.1 ‘ ’ 1

Residual standard error: 0.4675 on 10 degrees of freedom
Multiple R-squared:  0.6755,  Adjusted R-squared:  0.5457
F-statistic: 5.203 on 4 and 10 DF,  p-value: 0.01576
```

```
Analysis of Variance Table
Response: time50
              Df Sum Sq Mean Sq F value  Pr(>F)
time.sup       4 4.5481 1.13704  5.2034 0.01576 *
Residuals     10 2.1852 0.21852
---
Signif. codes:  0 ‘***’ 0.001 ‘**’ 0.01 ‘*’ 0.05 ‘.’ 0.1 ‘ ’ 1

Simultaneous Tests for General Linear Hypotheses
 contrast  estimate    SE df t.ratio p.value
 0h - 1h      0.222 0.382 10   0.582  0.9748
 0h - 4h      0.778 0.382 10   2.038  0.3159
 0h - 8h      0.333 0.382 10   0.873  0.9003
 0h - non    -0.889 0.382 10  -2.329  0.2128
 1h - 4h      0.556 0.382 10   1.456  0.6097
 1h - 8h      0.111 0.382 10   0.291  0.9982
 1h - non    -1.111 0.382 10  -2.911  0.0899
 4h - 8h     -0.444 0.382 10  -1.164  0.7704
 4h - non    -1.667 0.382 10  -4.367  0.0096
 8h - non    -1.222 0.382 10  -3.202  0.0574

P value adjustment: tukey method for comparing a family of 5
estimates
```

#### % regrowth (**Table S3**)

```
Model
Call:
lm(formula = percrgw ~ infstat * time.sup)

Residuals:
     Min       1Q   Median       3Q      Max
-24.2420  -2.3050   0.2322   2.6986  13.6348

Coefficients:
                        Estimate Std. Error t value Pr(>|t|)
(Intercept)              14.4177     4.8917   2.947 0.007964 **
infstatnon               26.0419     6.9180   3.764 0.001220 **
time.sup1h                0.9107     6.9180   0.132 0.896582
time.sup4h                8.3295     6.9180   1.204 0.242633
time.sup8h               22.9585     6.9180   3.319 0.003426 **
time.supnon             -11.0817     6.9180  -1.602 0.124862
infstatnon:time.sup1h    -2.9489     9.7835  -0.301 0.766209
infstatnon:time.sup4h    -8.2914     9.7835  -0.847 0.406752
infstatnon:time.sup8h   -39.1679     9.7835  -4.003 0.000698 ***
infstatnon:time.supnon  -27.7553     9.7835  -2.837 0.010188 *
---
Signif. codes:  0 ‘***’ 0.001 ‘**’ 0.01 ‘*’ 0.05 ‘.’ 0.1 ‘ ’ 1

Residual standard error: 8.473 on 20 degrees of freedom
Multiple R-squared:  0.8089,  Adjusted R-squared:  0.7229
F-statistic: 9.408 on 9 and 20 DF,  p-value: 1.823e-05



Analysis of Variance Table
Response: percrgw
                 Df Sum Sq Mean Sq F value    Pr(>F)
infstat           1  812.6  812.63 11.3200  0.003083 **
time.sup          4 3525.8  881.45 12.2786 3.338e-05 ***
infstat:time.sup  4 1740.0  435.01  6.0596  0.002311 **
Residuals        20 1435.7   71.79
---
Signif. codes:  0 ‘***’ 0.001 ‘**’ 0.01 ‘*’ 0.05 ‘.’ 0.1 ‘ ’ 1
```

### Fig. 6

```
Simultaneous Tests for General Linear Hypotheses
time.sup = 0h:
 contrast  estimate   SE df t.ratio p.value
 inf - non   -26.04 6.92 20  -3.764  0.0012

time.sup = 1h:
 contrast  estimate   SE df t.ratio p.value
 inf - non   -23.09 6.92 20  -3.338  0.0033

time.sup = 4h:
 contrast  estimate   SE df t.ratio p.value
 inf - non   -17.75 6.92 20  -2.566  0.0184

time.sup = 8h:
 contrast  estimate   SE df t.ratio p.value
 inf - non    13.13 6.92 20   1.897  0.0723

time.sup = non:
 contrast  estimate   SE df t.ratio p.value
 inf - non     1.71 6.92 20   0.248  0.8069

------


infstat = inf:
 contrast  estimate   SE df t.ratio p.value
 0h - 1h    -0.9107 6.92 20  -0.132  0.9999
 0h - 4h    -8.3295 6.92 20  -1.204  0.7491
 0h - 8h   -22.9585 6.92 20  -3.319  0.0252
 0h - non   11.0817 6.92 20   1.602  0.5131
 1h - 4h    -7.4188 6.92 20  -1.072  0.8183
 1h - 8h   -22.0478 6.92 20  -3.187  0.0334
 1h - non   11.9924 6.92 20   1.734  0.4372
 4h - 8h   -14.6290 6.92 20  -2.115  0.2526
 4h - non   19.4112 6.92 20   2.806  0.0728
 8h - non   34.0402 6.92 20   4.921  0.0007

infstat = non:
 contrast  estimate   SE df t.ratio p.value
 0h - 1h     2.0382 6.92 20   0.295  0.9982
 0h - 4h    -0.0381 6.92 20  -0.006  1.0000
 0h - 8h    16.2094 6.92 20   2.343  0.1726
 0h - non   38.8370 6.92 20   5.614  0.0001
 1h - 4h    -2.0763 6.92 20  -0.300  0.9981
 1h - 8h    14.1712 6.92 20   2.048  0.2802
 1h - non   36.7988 6.92 20   5.319  0.0003
 4h - 8h    16.2475 6.92 20   2.349  0.1709
 4h - non   38.8751 6.92 20   5.619  0.0001
 8h - non   22.6276 6.92 20   3.271  0.0279

P value adjustment: tukey method for comparing a family of 5
estimates
```

---

#### *Fig. 6c*
#### Fold change in PFU (**Table S3**)

```
Model
Call:
glm(formula = fch ~ ppn * time.sup, family = Gamma(link =
"log"))

Deviance Residuals:
     Min        1Q    Median        3Q       Max
-0.40520  -0.14962   0.01014   0.10014   0.36488

Coefficients:
                   Estimate  Std. Error t value Pr(>|t|)
(Intercept)         2.04122     0.13950  14.633  3.8e-12 ***
ppntotal            0.07904     0.19728   0.401  0.69291
time.sup0           0.24794     0.19728   1.257  0.22330
time.sup1           0.22228     0.19728   1.127  0.27319
time.sup4          -0.01747     0.19728  -0.089  0.93032
time.sup8          -0.29783     0.19728  -1.510  0.14675
ppntotal:time.sup0  0.64488     0.27899   2.311  0.03159 *
ppntotal:time.sup1  0.56800     0.27899   2.036  0.05522 .
ppntotal:time.sup4  0.83725     0.27899   3.001  0.00706 **
ppntotal:time.sup8  0.85402     0.27899   3.061  0.00616 **
---
Signif. codes:  0 ‘***’ 0.001 ‘**’ 0.01 ‘*’ 0.05 ‘.’ 0.1 ‘ ’ 1

(Dispersion parameter for Gamma family taken to be 0.05837704)
Null deviance: 6.6355  on 29  degrees of freedom
Residual deviance: 1.1880  on 20  degrees of freedom
AIC: 153.41
Number of Fisher Scoring iterations: 4


Analysis of Deviance Table
Model: Gamma, link: log
Response: fch
Terms added sequentially (first to last)
              Df Deviance Resid. Df Resid. Dev  Pr(>Chi)
NULL          29   6.6355
ppn           1    3.4800        28     3.1555 1.155e-14 ***
time.sup      4    1.2521        24     1.9034  0.000258 ***
ppn:time.sup  4    0.7154        20     1.1880  0.015554 *
---
Signif. codes:  0 ‘***’ 0.001 ‘**’ 0.01 ‘*’ 0.05 ‘.’ 0.1 ‘ ’ 1
```

```
Simultaneous Tests for General Linear Hypotheses
time.sup = non:
 contrast     ratio     SE df null t.ratio p.value
 free / total 1.202 0.3508 20    1   0.630  0.5360

time.sup = 0:
 contrast     ratio     SE df null t.ratio p.value
 free / total 0.341 0.0995 20    1  -3.686  0.0015

time.sup = 1:
 contrast     ratio     SE df null t.ratio p.value
 free / total 0.362 0.1056 20    1  -3.484  0.0023

time.sup = 4:
 contrast     ratio     SE df null t.ratio p.value
 free / total 0.440 0.1285 20    1  -2.811  0.0108

time.sup = 8:
 contrast     ratio     SE df null t.ratio p.value
 free / total 0.438 0.1278 20    1  -2.831  0.0103

Tests are performed on the log scale

------

ppn = free:
 contrast ratio    SE df null t.ratio p.value
 non / 0  1.560 0.455 20    1   1.524  0.5598
 non / 1  1.700 0.496 20    1   1.818  0.3910
 non / 4  1.764 0.515 20    1   1.944  0.3279
 non / 8  1.898 0.554 20    1   2.195  0.2218
 0 / 1    1.090 0.318 20    1   0.295  0.9982
 0 / 4    1.130 0.330 20    1   0.420  0.9930
 0 / 8    1.216 0.355 20    1   0.671  0.9604
 1 / 4    1.037 0.303 20    1   0.125  0.9999
 1 / 8    1.116 0.326 20    1   0.377  0.9954
 4 / 8    1.076 0.314 20    1   0.251  0.9990

ppn = total:
 contrast ratio    SE df null t.ratio p.value
 non / 0  0.443 0.129 20    1  -2.792  0.0748
 non / 1  0.512 0.149 20    1  -2.296  0.1873
 non / 4  0.646 0.189 20    1  -1.498  0.5756
 non / 8  0.691 0.202 20    1  -1.265  0.7145
 0 / 1    1.156 0.337 20    1   0.496  0.9868
 0 / 4    1.459 0.426 20    1   1.294  0.6976
 0 / 8    1.561 0.456 20    1   1.527  0.5581
 1 / 4    1.262 0.368 20    1   0.798  0.9281
 1 / 8    1.351 0.394 20    1   1.030  0.8384
 4 / 8    1.070 0.312 20    1   0.232  0.9993

P value adjustment: tukey method for comparing a family of 5
estimates
Tests are performed on the log scale
```

---

#### *Fig. 6d*

#### Fold change in PFU (**Table S3**)

**Model**

```
Call:
glm(formula = fch ~ time.sup * ppn, family = Gamma(link =
"log"))

Deviance Residuals:
     Min        1Q    Median        3Q       Max
-0.60325  -0.22578  -0.00879   0.24451   0.48321

Coefficients:
                      Estimate Std. Error t value Pr(>|t|)
(Intercept)            1.59601    0.22661   7.043 2.78e-06 ***
time.sup10            -0.72751    0.32048  -2.270  0.03737 *
time.sup6             -0.95416    0.32048  -2.977  0.00889 **
time.supnon            0.04943    0.32048   0.154  0.87934
ppntotal               1.85397    0.32048   5.785 2.79e-05 ***
time.sup10:ppntotal   -0.61428    0.45322  -1.355  0.19413
time.sup6:ppntotal    -0.13970    0.45322  -0.308  0.76188
time.supnon:ppntotal  -1.61996    0.45322  -3.574  0.00253 **
---
Signif. codes:  0 ‘***’ 0.001 ‘**’ 0.01 ‘*’ 0.05 ‘.’ 0.1 ‘ ’ 1

(Dispersion parameter for Gamma family taken to be 0.1540573)
Null deviance: 20.4004  on 23  degrees of freedom
Residual deviance:  2.7209  on 16  degrees of freedom
AIC: 118.87
Number of Fisher Scoring iterations: 4
```

**Analysis of Deviance Table**

```
Model: Gamma, link: log
Response: fch
Terms added sequentially (first to last)
             Df Deviance Resid. Df Resid. Dev       F    Pr(>F)
NULL                            23    20.4004
time.sup      3   6.9192        20    13.4811 14.9711 6.622e-05 ***
ppn           1   8.4183        19     5.0629 54.6436 1.522e-06 ***
time.sup:ppn  3   2.3419        16     2.7209  5.0672   0.01176 *
---
Signif. codes:  0 ‘***’ 0.001 ‘**’ 0.01 ‘*’ 0.05 ‘.’ 0.1 ‘ ’ 1
```

**Simultaneous Tests for General Linear Hypotheses**

```
ppn = free:
 contrast      ratio    SE df null t.ratio p.value
 0 / 10        2.070 0.663 16    1   2.270  0.1469
 0 / 6         2.596 0.832 16    1   2.977  0.0399
 0 / non       0.952 0.305 16    1  -0.154  0.9986
 10 / 6        1.254 0.402 16    1   0.707  0.8928
 10 / non      0.460 0.147 16    1  -2.424  0.1123
 6 / non       0.367 0.117 16    1  -3.132  0.0295

ppn = total:
 contrast      ratio    SE df null t.ratio p.value
 0 / 10        3.826 1.226 16    1   4.187  0.0035
 0 / 6         2.986 0.957 16    1   3.413  0.0168
 0 / non       4.809 1.541 16    1   4.901  0.0008
 10 / 6        0.780 0.250 16    1  -0.774  0.8652
 10 / non      1.257 0.403 16    1   0.714  0.8902
 6 / non       1.611 0.516 16    1   1.487  0.4673

P value adjustment: tukey method for comparing a family of 4
estimates
Tests are performed on the log scale

------

time.sup = 0:
 contrast     ratio     SE df null t.ratio p.value
 free / total 0.157 0.0502 16    1  -5.785  <.0001

time.sup = 10:
 contrast     ratio     SE df null t.ratio p.value
 free / total 0.289 0.0928 16    1  -3.868  0.0014

time.sup = 6:
 contrast     ratio     SE df null t.ratio p.value
 free / total 0.180 0.0577 16    1  -5.349  0.0001

time.sup = non:
 contrast     ratio     SE df null t.ratio p.value
 free / total 0.791 0.2536 16    1  -0.730  0.4758

Tests are performed on the log scale
```

#### Supplementary figure 2 (Fig. S2)

### Supplementary figure S2

#### *Fig. S2a (main)*

##### Growth rates at selected time-points (**Table S3**)

**Model**

```
Call:
lm(formula = r0 ~ time.fac)

Residuals:
      Min        1Q    Median        3Q       Max 
-0.059545 -0.015308 -0.004961  0.012000  0.071028 

Coefficients:
            Estimate Std. Error t value Pr(>|t|)    
(Intercept)  1.14658    0.01951   58.77 4.94e-14 ***
time.fac4   -0.68787    0.02759  -24.93 2.46e-10 ***
time.fac6   -1.07058    0.02759  -38.80 3.08e-12 ***
time.fac10  -1.23256    0.02759  -44.68 7.59e-13 ***
time.fac30  -1.16483    0.02759  -42.22 1.33e-12 ***
---
Signif. codes:  0 '***' 0.001 '**' 0.01 '*' 0.05 '.' 0.1 ' ' 1

Residual standard error: 0.03379 on 10 degrees of freedom
Multiple R-squared:  0.9964,  Adjusted R-squared:  0.9949 
F-statistic: 683.9 on 4 and 10 DF,  p-value: 3.834e-12
```

**Analysis of Variance Table**

```
Response: r0
          Df  Sum Sq Mean Sq F value    Pr(>F)    
time.fac   4 3.12342 0.78086  683.91 3.834e-12 ***
Residuals 10 0.01142 0.00114                      
---
Signif. codes:  0 '***' 0.001 '**' 0.01 '*' 0.05 '.' 0.1 ' ' 1
```

**Simultaneous Tests for General Linear Hypotheses**

```
Multiple Comparisons of Means: Tukey Contrasts
Fit: lm(formula = r0 ~ time.fac)

Linear Hypotheses:
contrast      Estimate Std. Error t value Pr(>|t|)    
4 - 2 == 0    -0.68787    0.02759 -24.932  <0.0001 ***
6 - 2 == 0    -1.07058    0.02759 -38.804  <0.0001 ***
10 - 2 == 0   -1.23256    0.02759 -44.676  <0.0001 ***
30 - 2 == 0   -1.16483    0.02759 -42.221  <0.0001 ***
6 - 4 == 0    -0.38271    0.02759 -13.872  <0.0001 ***
10 - 4 == 0   -0.54470    0.02759 -19.743  <0.0001 ***
30 - 4 == 0   -0.47697    0.02759 -17.288  <0.0001 ***
10 - 6 == 0   -0.16199    0.02759  -5.871   0.0012 ** 
30 - 6 == 0   -0.09426    0.02759  -3.416   0.0412 *  
30 - 10 == 0   0.06773    0.02759   2.455   0.1777    
---
Signif. codes:  0 '***' 0.001 '**' 0.01 '*' 0.05 '.' 0.1 ' ' 1
(Adjusted p values reported -- single-step method)
```

#### *Fig. S2a (inset)*

##### Growth rate~time relationship

**Determining breakpoints (mod_bp)**

```
Call:
lm(formula = r0 ~ time)

Residuals:
    Min      1Q  Median      3Q     Max 
-0.3826 -0.2690 -0.2238  0.1908  0.9173 

Coefficients:
             Estimate Std. Error t value Pr(>|t|)    
(Intercept)  0.514724   0.082286   6.255 1.20e-07 ***
time        -0.025914   0.005979  -4.334 7.88e-05 ***
---
Signif. codes:  0 '***' 0.001 '**' 0.01 '*' 0.05 '.' 0.1 ' ' 1

Residual standard error: 0.3723 on 46 degrees of freedom
Multiple R-squared:    0.29,  Adjusted R-squared:  0.2745 
F-statistic: 18.79 on 1 and 46 DF,  p-value: 7.876e-05
```

*****Regression Model with Segmented Relationship(s)*****

```
Call:
segmented.lm(obj = mod_bp, seg.Z = ~time, psi = c(5))

Estimated Break-Point(s):
              Est. St.Err
psi1.time 5.228   0.15

Meaningful coefficients of the linear terms:
            Estimate Std. Error t value Pr(>|t|)    
(Intercept)  1.75763    0.06330   27.77   <2e-16 ***
time        -0.33354    0.01782  -18.71   <2e-16 ***
U1.time      0.33161    0.01794   18.48       NA    
---
Signif. codes:  0 '***' 0.001 '**' 0.01 '*' 0.05 '.' 0.1 ' ' 1

Residual standard error: 0.1009 on 44 degrees of freedom
Multiple R-Squared: 0.9502,  Adjusted R-squared: 0.9468 

Convergence attained in 2 iter. (rel. change 0)
```

**Model (panel)**

```
Call:
lm(formula = r0 ~ time * (time >= 5.228))

Residuals:
     Min       1Q   Median       3Q      Max 
-0.16364 -0.05825  0.00366  0.03649  0.36657 

Coefficients:
                       Estimate Std. Error t value Pr(>|t|)    
(Intercept)             1.75763    0.06330   27.77   <2e-16 ***
time                   -0.33354    0.01782  -18.71   <2e-16 ***
time >= 5.228TRUE      -1.73373    0.07259  -23.88   <2e-16 ***
time:time >= 5.228TRUE  0.33161    0.01794   18.48   <2e-16 ***
---
Signif. codes:  0 '***' 0.001 '**' 0.01 '*' 0.05 '.' 0.1 ' ' 1

Residual standard error: 0.1009 on 44 degrees of freedom
Multiple R-squared:  0.9502,  Adjusted R-squared:  0.9468 
F-statistic: 279.6 on 3 and 44 DF,  p-value: < 2.2e-16
```

**Analysis of Variance Table**

```
Response: r0
                  Df Sum Sq Mean Sq F value    Pr(>F)    
time               1 2.6046  2.6046  255.98 < 2.2e-16 ***
time >= 5.228      1 2.4541  2.4541  241.19 < 2.2e-16 ***
time:time >= 5.228 1 3.4757  3.4757  341.58 < 2.2e-16 ***
Residuals         44 0.4477  0.0102                      
---
Signif. codes:  0 '***' 0.001 '**' 0.01 '*' 0.05 '.' 0.1 ' ' 1
```

---

#### Supplementary figure 2 (Fig. S2)

##### *Fig. S2b*

##### CFU~time*infection status relationship

**Model (panel)**

```
Call:
glm(formula = CFU ~ time.fac, family = quasipoisson(link =
“log))

Deviance Residuals:
    Min       1Q   Median       3Q      Max
-8943.0  -2830.2   -959.7   1943.6  10364.5

Coefficients:
            Estimate Std. Error t value Pr(>|t|)
(Intercept)  18.9474     0.2737  69.235 9.63e-15 ***
time.fac4     1.3032     0.3086   4.223  0.00176 **
time.fac6     1.0237     0.3191   3.208  0.00936 **
time.fac10    0.9473     0.3224   2.938  0.01483 *
time.fac30    1.0680     0.3172   3.367  0.00716 **
---
Signif. codes:  0 ‘***’ 0.001 ‘**’ 0.01 ‘*’ 0.05 ‘.’ 0.1 ‘ ’ 1

(Dispersion parameter for quasipoisson family taken to be
38046245)
Null deviance: 1253186279  on 14  degrees of freedom
Residual deviance:  371947846  on 10  degrees of freedom
AIC: NA
Number of Fisher Scoring iterations: 4
```

**Analysis of Deviance Table**

```
Model: quasipoisson, link: log
Response: CFU
Terms added sequentially (first to last)
         Df  Deviance Resid. Df Resid. Dev      F Pr(>F)
NULL                         14 1253186279
time.fac  4 881238433        10  371947846 5.7906 0.0112 *
---
Signif. codes:  0 ‘***’ 0.001 ‘**’ 0.01 ‘*’ 0.05 ‘.’ 0.1 ‘ ’ 1
```

**Simultaneous Tests for General Linear Hypotheses**

```
 contrast  ratio     SE  df null z.ratio p.value
 2.25 / 4  0.272 0.0838 Inf    1  -4.223  0.0002
 2.25 / 6  0.359 0.1146 Inf    1  -3.208  0.0117
 2.25 / 10 0.388 0.1250 Inf    1  -2.938  0.0273
 2.25 / 30 0.344 0.1090 Inf    1  -3.367  0.0068
 4 / 6     1.322 0.2875 Inf    1   1.286  0.7000
 4 / 10    1.427 0.3172 Inf    1   1.602  0.4964
 4 / 30    1.265 0.2716 Inf    1   1.096  0.8088
 6 / 10    1.079 0.2553 Inf    1   0.323  0.9977
 6 / 30    0.957 0.2195 Inf    1  -0.193  0.9997
 10 / 30   0.886 0.2075 Inf    1  -0.516  0.9858

P value adjustment: tukey method for comparing a family of 5
estimates
Tests are performed on the log scale
```

---

##### *Fig. S2d*

##### Length~time relationship

**Model (panel, top)**

```
Call:
lm(formula = log(len) ~ time.fac)

Residuals:
     Min       1Q   Median       3Q      Max
-1.06943 -0.19627 -0.00677  0.19274  1.27935

Coefficients:
            Estimate Std. Error t value Pr(>|t|)
(Intercept)  1.69976    0.01641 103.588   <2e-16 ***
time.fac4   -0.22651    0.02361  -9.594   <2e-16 ***
time.fac6   -0.35080    0.02218 -15.818   <2e-16 ***
time.fac10  -0.54576    0.02184 -24.985   <2e-16 ***
time.fac30  -0.66469    0.02034 -32.675   <2e-16 ***
---
Signif. codes:  0 ‘***’ 0.001 ‘**’ 0.01 ‘*’ 0.05 ‘.’ 0.1 ‘ ’ 1
Residual standard error: 0.2866 on 1917 degrees of freedom
Multiple R-squared:  0.403,   Adjusted R-squared:  0.4018
F-statistic: 323.6 on 4 and 1917 DF,  p-value: < 2.2e-16
```

**Anova Table (Type II tests)**

```
Response: log(len)
          Sum Sq   Df F value    Pr(>F)
time.fac  106.28    4  323.56 < 2.2e-16 ***
Residuals 157.43 1917
---
Signif. codes:  0 ‘***’ 0.001 ‘**’ 0.01 ‘*’ 0.05 ‘.’ 0.1 ‘ ’ 1
```

**Simultaneous Tests for General Linear Hypotheses**

```
 contrast  ratio     SE   df null t.ratio p.value
 2.25 / 4   1.25 0.0296 1917    1   9.594  <.0001
 2.25 / 6   1.42 0.0315 1917    1  15.818  <.0001
 2.25 / 10  1.73 0.0377 1917    1  24.985  <.0001
 2.25 / 30  1.94 0.0395 1917    1  32.675  <.0001
 4 / 6      1.13 0.0256 1917    1   5.500  <.0001
 4 / 10     1.38 0.0306 1917    1  14.334  <.0001
 4 / 30     1.55 0.0322 1917    1  21.065  <.0001
 6 / 10     1.22 0.0252 1917    1   9.397  <.0001
 6 / 30     1.37 0.0262 1917    1  16.382  <.0001
 10 / 30    1.13 0.0211 1917    1   6.335  <.0001

P value adjustment: tukey method for comparing a family of 5
estimates
Tests are performed on the log scale
```

##### Width~time relationship

**Model (panel, 2nd from top)**

```
Call:
lm(formula = wid ~ time.fac)

Residuals:
      Min        1Q    Median        3Q       Max
-0.249401 -0.040435  0.003216  0.044287  0.212720

Coefficients:
             Estimate Std. Error t value Pr(>|t|)
(Intercept)  0.820566   0.003638 225.558  < 2e-16 ***
time.fac4   -0.030603   0.005234  -5.847 5.88e-09 ***
time.fac6   -0.038977   0.004917  -7.928 3.76e-15 ***
time.fac10  -0.069149   0.004843 -14.278  < 2e-16 ***
time.fac30  -0.025495   0.004510  -5.653 1.82e-08 ***
---
Signif. codes:  0 ‘***’ 0.001 ‘**’ 0.01 ‘*’ 0.05 ‘.’ 0.1 ‘ ’ 1

Residual standard error: 0.06353 on 1917 degrees of freedom
Multiple R-squared:  0.1036,  Adjusted R-squared:  0.1017 F-
statistic: 55.37 on 4 and 1917 DF,  p-value: < 2.2e-16
```

**Anova Table (Type II tests)**

```
Response: wid
          Sum Sq   Df F value    Pr(>F)
time.fac  0.8940    4  55.368 < 2.2e-16 ***
Residuals 7.7381 1917
---
Signif. codes:  0 ‘***’ 0.001 ‘**’ 0.01 ‘*’ 0.05 ‘.’ 0.1 ‘ ’ 1
```

**Simultaneous Tests for General Linear Hypotheses**

```
Multiple Comparisons of Means: Tukey Contrasts
Fit: lm(formula = wid ~ time.fac)

Linear Hypotheses:
contrast         Estimate Std. Error t value Pr(>|t|)
4 - 2.25 == 0   -0.030603   0.005234  -5.847   <0.001 ***
6 - 2.25 == 0   -0.038977   0.004917  -7.928   <0.001 ***
10 - 2.25 == 0  -0.069149   0.004843 -14.278   <0.001 ***
30 - 2.25 == 0  -0.025495   0.004510  -5.653   <0.001 ***
6 - 4 == 0      -0.008374   0.005010  -1.671   0.4497
10 - 4 == 0     -0.038546   0.004938  -7.806   <0.001 ***
30 - 4 == 0      0.005108   0.004612   1.108   0.8012
10 - 6 == 0     -0.030172   0.004600  -6.559   <0.001 ***
30 - 6 == 0      0.013482   0.004248   3.174   0.0131 *
30 - 10 == 0     0.043654   0.004162  10.488   <0.001 ***
---
Signif. codes:  0 ‘***’ 0.001 ‘**’ 0.01 ‘*’ 0.05 ‘.’ 0.1 ‘ ’ 1
(Adjusted p values reported -- single-step method)
```

#### Supplementary figure 2 (Fig. S2)

### SA~time relationship

**Model (panel, 3rd from top)**

```
Call:
lm(formula = log(vol) ~ time.fac)

Residuals:
     Min       1Q   Median       3Q      Max 
-1.15622 -0.23615  0.00186  0.22687  1.27594 

Coefficients:
            Estimate Std. Error t value Pr(>|t|)    
(Intercept)  1.00461    0.01922   52.28   <2e-16 ***
time.fac4   -0.31448    0.02765  -11.37   <2e-16 ***
time.fac6   -0.46877    0.02597  -18.05   <2e-16 ***
time.fac10  -0.75820    0.02558  -29.64   <2e-16 ***
time.fac30  -0.78251    0.02382  -32.85   <2e-16 ***
---
Signif. codes:  0 ‘***’ 0.001 ‘**’ 0.01 ‘*’ 0.05 ‘.’ 0.1 ‘ ’ 1
Residual standard error: 0.3356 on 1917 degrees of freedom
Multiple R-squared:  0.421,   Adjusted R-squared:  0.4198 
F-statistic: 348.5 on 4 and 1917 DF,  p-value: < 2.2e-16
```

**Anova Table (Type II tests)**

```
Response: log(vol)
          Sum Sq   Df F value    Pr(>F)    
time.fac   157.0    4  348.52 < 2.2e-16 ***
Residuals  215.9 1917                      
---
Signif. codes:  0 ‘***’ 0.001 ‘**’ 0.01 ‘*’ 0.05 ‘.’ 0.1 ‘ ’ 1
```

**Simultaneous Tests for General Linear Hypotheses**

```
 contrast         ratio     SE   df null t.ratio p.value
 2.25 / 4    1.37 0.0379 1917    1  11.374  <.0001
 2.25 / 6    1.60 0.0415 1917    1  18.050  <.0001
 2.25 / 10   2.13 0.0546 1917    1  29.639  <.0001
 2.25 / 30   2.19 0.0521 1917    1  32.847  <.0001
 4 / 6       1.17 0.0309 1917    1   5.830  <.0001
 4 / 10      1.56 0.0406 1917    1  17.012  <.0001
 4 / 30      1.60 0.0389 1917    1  19.212  <.0001
 6 / 10      1.34 0.0325 1917    1  11.912  <.0001
 6 / 30      1.37 0.0307 1917    1  13.982  <.0001
 10 / 30     1.02 0.0225 1917    1   1.106  0.8035

P value adjustment: tukey method for comparing a family of 5
estimates 
Tests are performed on the log scale
```

### Volume~time relationship

**Model (panel, bottom)**

```
Call:
lm(formula = log(surf) ~ time.fac)

Residuals:
     Min       1Q   Median       3Q      Max 
-1.04660 -0.20675 -0.00717  0.19434  1.24416 

Coefficients:
            Estimate Std. Error t value Pr(>|t|)    
(Intercept)  2.64419    0.01668  158.48   <2e-16 ***
time.fac4   -0.26431    0.02401  -11.01   <2e-16 ***
time.fac6   -0.39965    0.02255  -17.72   <2e-16 ***
time.fac10  -0.63430    0.02221  -28.56   <2e-16 ***
time.fac30  -0.69862    0.02068  -33.77   <2e-16 ***
---
Signif. codes:  0 ‘***’ 0.001 ‘**’ 0.01 ‘*’ 0.05 ‘.’ 0.1 ‘ ’ 1
Residual standard error: 0.2914 on 1917 degrees of freedom
Multiple R-squared:  0.4265,  Adjusted R-squared:  0.4253 
F-statistic: 356.4 on 4 and 1917 DF,  p-value: < 2.2e-16
```

**Anova Table (Type II tests)**

```
Response: log(surf)
          Sum Sq   Df F value    Pr(>F)    
time.fac  121.04    4  356.39 < 2.2e-16 ***
Residuals 162.76 1917                      
---
Signif. codes:  0 ‘***’ 0.001 ‘**’ 0.01 ‘*’ 0.05 ‘.’ 0.1 ‘ ’ 1
```

**Simultaneous Tests for General Linear Hypotheses**

```
 contrast  ratio     SE   df null t.ratio p.value
 2.25 / 4    1.30 0.0313 1917    1  11.010  <.0001
 2.25 / 6    1.49 0.0336 1917    1  17.723  <.0001
 2.25 / 10   1.89 0.0419 1917    1  28.558  <.0001
 2.25 / 30   2.01 0.0416 1917    1  33.775  <.0001
 4 / 6       1.14 0.0263 1917    1   5.890  <.0001
 4 / 10      1.45 0.0328 1917    1  16.338  <.0001
 4 / 30      1.54 0.0327 1917    1  20.533  <.0001
 6 / 10      1.26 0.0267 1917    1  11.123  <.0001
 6 / 30      1.35 0.0263 1917    1  15.346  <.0001
 10 / 30     1.07 0.0204 1917    1   3.369  0.0069

P value adjustment: tukey method for comparing a family of 5
estimates 
Tests are performed on the log scale
```

### % at 10h within morphological range at 30h

| length | | width | | surface area | | volume | |
|---|---|---|---|---|---|---|---|
| **mean % within range** | **SD % within range** | **mean % within range** | **SD % within range** | **mean % within range** | **SD % within range** | **mean % within range** | **SD % within range** |
| 97.1 | 1.73 | 100 | 0 | 97.5 | 0.708 | 98.0 | 0.152 |

==================================================================

Supplementary figure 5 (Fig. S5)

#### Supplementary figure S5

##### *Fig. S5a (main)*

#### Maximum growth rate (**Table S3**)

```
Welch Two Sample t-test growth rate (Maximum growth rate)
t = -0.75892, df = 3.037, p-value = 0.5025
alternative hypothesis: true difference in means is not equal to 0
95 percent confidence interval:
 -0.6030645  0.3695288
sample estimates:
mean of x mean of y
 1.426566  1.543334
```

#### Maximum decline rate & time at max decline rate (**Table S3**)

```
Welch Two Sample t-test (Max decline rate)
t = 0.31946, df = 2.0727, p-value = 0.7787
alternative hypothesis: true difference in means is not equal to
0
95 percent confidence interval:
 -0.2069002  0.2413089
sample estimates:
  mean of x   mean of y
-0.05956082 -0.07676515
```

```
Welch Two Sample t-test (Time at max decline rate)
t = -1.8579, df = 2.0248, p-value = 0.2027
alternative hypothesis: true difference in means is not equal to
0
95 percent confidence interval:
 -44.94860  17.61526
sample estimates:
mean of x mean of y
  9.00000  22.66667
```

#### % decrease OD (**Table S3**)

```
Welch Two Sample t-test (Table S3)
t = -0.55629, df = 3.9189, p-value = 0.6082
alternative hypothesis: true difference in means is not equal to 0
95 percent confidence interval:
 -12.483163   8.344168
sample estimates:
mean of x mean of y
 57.08299  59.15249
```

#### Growth rates at selected time-points (**Table S3**)

```
Model
Call:
lm(formula = log(r0 + 1) ~ time.fac + strain)

Residuals:
      Min        1Q    Median        3Q       Max
-0.044551 -0.009031  0.000949  0.012265  0.046462

Coefficients:
                Estimate Std. Error t value Pr(>|t|)
(Intercept)     0.693577   0.010072  68.864   <2e-16 ***
time.fac4      -0.359924   0.012740 -28.252   <2e-16 ***
time.fac6      -0.649235   0.012740 -50.961   <2e-16 ***
time.fac10     -0.739482   0.012740 -58.045   <2e-16 ***
strain2778      0.004661   0.009008   0.517    0.611
---
Signif. codes:  0 ‘***’ 0.001 ‘**’ 0.01 ‘*’ 0.05 ‘.’ 0.1 ‘ ’ 1

Residual standard error: 0.02207 on 19 degrees of freedom
Multiple R-squared:  0.9954,  Adjusted R-squared:  0.9944
F-statistic:  1027 on 4 and 19 DF,  p-value: < 2.2e-16
```

```
Analysis of Variance Table
Response: log(r0 + 1)
           Df  Sum Sq Mean Sq   F value Pr(>F)
time.fac    3 2.00069 0.66690 1369.6708 <2e-16 ***
strain      1 0.00013 0.00013    0.2677 0.6108
Residuals  19 0.00925 0.00049
---
Signif. codes:  0 ‘***’ 0.001 ‘**’ 0.01 ‘*’ 0.05 ‘.’ 0.1 ‘ ’ 1

Simultaneous Tests for General Linear Hypotheses
Multiple Comparisons of Means: Tukey Contrasts
Fit: lm(formula = log(r0 + 1) ~ time.fac + strain)

Linear Hypotheses:
contrast       Estimate Std. Error t value Pr(>|t|)
4 - 2.25 == 0  -0.35992    0.01274 -28.252   <1e-05 ***
6 - 2.25 == 0  -0.64924    0.01274 -50.961   <1e-05 ***
10 - 2.25 == 0 -0.73948    0.01274 -58.045   <1e-05 ***
6 - 4 == 0     -0.28931    0.01274 -22.709   <1e-05 ***
10 - 4 == 0    -0.37956    0.01274 -29.793   <1e-05 ***
10 - 6 == 0    -0.09025    0.01274  -7.084   <1e-05 ***
---
Signif. codes:  0 ‘***’ 0.001 ‘**’ 0.01 ‘*’ 0.05 ‘.’ 0.1 ‘ ’ 1
(Adjusted p values reported -- single-step method)
```

### Supplementary figure 5 (Fig. S5)

#### *Fig. S5a (inset)*

#### Growth rate~time relationship

**Determining breakpoints (mod_bp)**

```
Call:
lm(formula = log(r0 + 1) ~ time * strain)

Residuals:
     Min       1Q   Median       3Q      Max 
-0.30024 -0.24116  0.01836  0.17116  0.61765 

Coefficients:
                   Estimate Std. Error t value Pr(>|t|)    
(Intercept)            0.378468   0.062247   6.080 4.82e-08 ***
time                  -0.017212   0.004283  -4.019 0.000139 ***
strainpsyB             0.020109   0.088031   0.228 0.819938    
time:strainpsyB       -0.001036   0.006057  -0.171 0.864646    
---
Signif. codes:  0 ‘***’ 0.001 ‘**’ 0.01 ‘*’ 0.05 ‘.’ 0.1 ‘ ’ 1

Residual standard error: 0.2627 on 74 degrees of freedom
Multiple R-squared:  0.3169,  Adjusted R-squared:  0.2892 
F-statistic: 11.44 on 3 and 74 DF,  p-value: 3.015e-06
```

*****Regression Model with Segmented Relationship(s)*****

```
Call: 
segmented.lm(obj = mod_bp, seg.Z = ~time, psi = c(5))

Estimated Break-Point(s):
              Est. St.Err
psi1.time 5.691  0.115

Meaningful coefficients of the linear terms:
                   Estimate Std. Error t value Pr(>|t|)    
(Intercept)       1.111464   0.024119  46.082   <2e-16 ***
time             -0.196395   0.006762 -29.045   <2e-16 ***
strainpsyB        0.020109   0.017084   1.177    0.243    
U1.time           0.196527   0.006778  28.996       NA    
time:strainpsyB  -0.001036   0.001175  -0.881    0.381    
---
Signif. codes:  0 ‘***’ 0.001 ‘**’ 0.01 ‘*’ 0.05 ‘.’ 0.1 ‘ ’ 1

Residual standard error: 0.05099 on 72 degrees of freedom
Multiple R-Squared: 0.975,  Adjusted R-squared: 0.9732 
Convergence attained in 2 iter. (rel. change 0)
```

**Model (panel)**

```
Call:
lm(formula = log(r0 + 1) ~ time * (time >= 5.691) + strain)

Residuals:
      Min        1Q    Median        3Q       Max 
-0.191478 -0.031843  0.004164  0.024822  0.105172 

Coefficients:
                         Estimate Std. Error t value Pr(>|t|)    
(Intercept)              1.117013   0.023247  48.049   <2e-16 ***
time                    -0.196913   0.006726 -29.277   <2e-16 ***
time >= 5.691TRUE       -1.118383   0.026389 -42.381   <2e-16 ***
strainpsyB               0.009011   0.011529   0.782    0.437    
time:time >= 5.691TRUE   0.196527   0.006767  29.040   <2e-16 ***
---
Signif. codes:  0 ‘***’ 0.001 ‘**’ 0.01 ‘*’ 0.05 ‘.’ 0.1 ‘ ’ 1

Residual standard error: 0.05091 on 73 degrees of freedom
Multiple R-squared:  0.9747,  Adjusted R-squared:  0.9733 
F-statistic:   703 on 4 and 73 DF,  p-value: < 2.2e-16
```

**Analysis of Variance Table**

```
Response: log(r0 + 1)
                   Df  Sum Sq Mean Sq   F value Pr(>F)    
time                1 2.36620 2.36620  912.8728 <2e-16 ***
time >= 5.691       1 2.73537 2.73537 1055.2991 <2e-16 ***
strain              1 0.00158 0.00158    0.6109  0.437    
time:time >= 5.691  1 2.18591 2.18591  843.3178 <2e-16 ***
Residuals          73 0.18922 0.00259                     
---
Signif. codes:  0 ‘***’ 0.001 ‘**’ 0.01 ‘*’ 0.05 ‘.’ 0.1 ‘ ’ 1
```

---

#### *Fig. S5c*

#### GFP foci~growth phase*strain relationship

**Model (panel)**

```
Call:
glm(formula = maxima ~ time.fac * strain, family = 
quasipoisson(link = log))

Deviance Residuals: 
    Min       1Q   Median       3Q      Max  
-5.4321  -1.1509  -0.1663   0.5274   8.9609  

Coefficients:
                     Estimate Std. Error t value Pr(>|t|)    
(Intercept)           1.32914    0.03277   40.56   <2e-16 ***
time.fac4            -1.16737    0.06218  -18.77   <2e-16 ***
time.fac6            -1.64661    0.07208  -22.84   <2e-16 ***
time.fac10           -1.74126    0.07394  -23.55   <2e-16 ***
strainpsyB            1.36237    0.03744   36.39   <2e-16 ***
time.fac4:strainpsyB  0.94925    0.06768   14.03   <2e-16 ***
time.fac6:strainpsyB  1.37011    0.07650   17.91   <2e-16 ***
time.fac10:strainpsyB 1.25585    0.07870   15.96   <2e-16 ***
---
Signif. codes:  0 ‘***’ 0.001 ‘**’ 0.01 ‘*’ 0.05 ‘.’ 0.1 ‘ ’ 1

(Dispersion parameter for quasipoisson family taken to be 
1.533424)
Null deviance: 22398.3  on 3374  degrees of freedom
Residual deviance:  5005.8  on 3367  degrees of freedom
AIC: NA

Number of Fisher Scoring iterations: 5
```

**Analysis of Deviance Table**

```
Model: quasipoisson, link: log
Response: maxima
Terms added sequentially (first to last)
                Df Deviance Resid. Df Resid. Dev  Pr(>Chi)    
NULL                             3374    22398.3              
time.fac         3   1286.0      3371    21112.2 < 2.2e-16 ***
strain           1  15302.5      3370     5809.7 < 2.2e-16 ***
time.fac:strain  3    803.9      3367     5005.8 < 2.2e-16 ***
---
Signif. codes:  0 ‘***’ 0.001 ‘**’ 0.01 ‘*’ 0.05 ‘.’ 0.1 ‘ ’ 1
```

**Simultaneous Tests for General Linear Hypotheses**

```
time.fac = 2.25:
 contrast     ratio      SE  df null z.ratio p.value
 WT / psyB   0.2561 0.00959 Inf    1 -36.389  <.0001

time.fac = 4:
 contrast     ratio      SE  df null z.ratio p.value
 WT / psyB   0.0991 0.00559 Inf    1 -40.999  <.0001

time.fac = 6:
 contrast     ratio      SE  df null z.ratio p.value
 WT / psyB   0.0651 0.00434 Inf    1 -40.956  <.0001

time.fac = 10:
 contrast     ratio      SE  df null z.ratio p.value
 WT / psyB   0.0729 0.00505 Inf    1 -37.822  <.0001

Tests are performed on the log scale

------

strain = WT:
contrast  ratio     SE df null t.ratio p.value
 2.25 / 4    3.21 0.1998 Inf    1  18.773  <.0001
 2.25 / 6    5.19 0.3741 Inf    1  22.844  <.0001
 2.25 / 10   5.70 0.4218 Inf    1  23.549  <.0001
 4 / 6       1.61 0.1343 Inf    1   5.763  <.0001
 4 / 10      1.78 0.1505 Inf    1   6.770  <.0001
 6 / 10      1.10 0.1014 Inf    1   1.026  0.7344

strain = psyB:
contrast  ratio     SE df null t.ratio p.value
 2.25 / 4    1.24 0.0332 Inf    1   8.164  <.0001
 2.25 / 6    1.32 0.0338 Inf    1  10.785  <.0001
 2.25 / 10   1.62 0.0438 Inf    1  18.014  <.0001
 4 / 6       1.06 0.0284 Inf    1   2.183  0.1279
 4 / 10      1.31 0.0366 Inf    1   9.546  <.0001
 6 / 10      1.23 0.0332 Inf    1   7.745  <.0001

P value adjustment: tukey method for comparing a family of 4 
estimates
Tests are performed on the log scale
```

### Supplementary figure 5 (Fig. S5)

#### % no foci~growth phase*strain relationship

**Model (panel)**

```
Call:
glm(formula = (percNF) ~ time.fac + strain, family =
quasipoisson(link = "inverse"))

Deviance Residuals:
    Min       1Q   Median       3Q      Max
-5.0157  -0.8849  -0.8711   1.1087   3.0849

Coefficients:
            Estimate Std. Error t value Pr(>|t|)
(Intercept)  0.17101    0.08075   2.118   0.0476 *
time.fac4   -0.13795    0.08104  -1.702   0.1050
time.fac6   -0.15214    0.08080  -1.883   0.0751 .
time.fac10  -0.14948    0.08083  -1.849   0.0800 .
strainpsyB   2.53522    2.38468   1.063   0.3011
---
Signif. codes:  0 ‘***’ 0.001 ‘**’ 0.01 ‘*’ 0.05 ‘.’ 0.1 ‘ ’ 1

(Dispersion parameter for quasipoisson family taken to be
3.91228)
Null deviance: 731.251  on 23  degrees of freedom
Residual deviance:  65.024  on 19  degrees of freedom
AIC: NA
Number of Fisher Scoring iterations: 9
```

**Analysis of Deviance Table**

```
Model: quasipoisson, link: inverse
Response: (percNF)
Terms added sequentially (first to last)

         Df Deviance Resid. Df Resid. Dev  Pr(>Chi)
NULL                        23     731.25
time.fac  3   136.32        20     594.93 1.314e-07 ***
strain    1   529.90        19      65.02 < 2.2e-16 ***
---
Signif. codes:  0 ‘***’ 0.001 ‘**’ 0.01 ‘*’ 0.05 ‘.’ 0.1 ‘ ’ 1
```

**Simultaneous Tests for General Linear Hypotheses**

```
time.fac = 2.25:
 contrast    estimate   SE  df z.ratio p.value
 WT - psyB       5.48 2.77 Inf   1.976  0.0481

time.fac = 4:
 contrast    estimate   SE  df z.ratio p.value
 WT - psyB      29.86 6.29 Inf   4.747  <.0001

time.fac = 6:
 contrast    estimate   SE  df z.ratio p.value
 WT - psyB      52.59 8.32 Inf   6.321  <.0001

time.fac = 10:
 contrast    estimate   SE  df z.ratio p.value
 WT - psyB      46.06 7.79 Inf   5.912  <.0001

------

strain = WT:
 contrast   estimate       SE  df z.ratio p.value
 2.25 - 4  -24.39714  6.86047 Inf  -3.556  0.0021
 2.25 - 6  -47.13544  8.75895 Inf  -5.381  <.0001
 2.25 - 10 -40.60001  8.25810 Inf  -4.916  <.0001
 4 - 6     -22.73831 10.41810 Inf  -2.183  0.1280
 4 - 10    -16.20287 10.00070 Inf  -1.620  0.3671
 6 - 10      6.53543 11.38714 Inf   0.574  0.9399

strain = psyB:
 contrast   estimate       SE  df z.ratio p.value
 2.25 - 4   -0.01985  0.03766 Inf  -0.527  0.9526
 2.25 - 6   -0.02201  0.04152 Inf  -0.530  0.9518
 2.25 - 10  -0.02160  0.04079 Inf  -0.530  0.9519
 4 - 6      -0.00216  0.00419 Inf  -0.517  0.9551
 4 - 10     -0.00176  0.00348 Inf  -0.505  0.9578
 6 - 10      0.00041  0.00104 Inf   0.390  0.9799

P value adjustment: tukey method for comparing a family of 4
estimates
```

==================================================================

Supplementary figure 6 (Fig. S6)

### Figure S6

#### *Fig. S6a*

##### OD~time*infection status relationship

```
Model (panel):
Call:
glm(formula = od ~ time + I(time^2) * infstat, family =
Gamma(link = "log"))

Deviance Residuals:
     Min        1Q    Median        3Q       Max
-0.58650  -0.11840  -0.01304   0.10230   0.31420

Coefficients:
                        Estimate Std. Error t value Pr(>|t|)
(Intercept)            -2.517666   0.269928  -9.327 2.39e-13 ***
time                    1.216943   0.138168   8.808 1.82e-12 ***
I(time^2)              -0.098540   0.017017  -5.791 2.62e-07 ***
infstatinf              0.842309   0.082880  10.163 9.57e-15 ***
I(time^2):infstatinf   -0.117825   0.004886 -24.117  < 2e-16 ***
---
Signif. codes:  0 '***' 0.001 '**' 0.01 '*' 0.05 '.' 0.1 ' ' 1

(Dispersion parameter for Gamma family taken to be 0.03020798)
Null deviance: 29.3928  on 65  degrees of freedom
Residual deviance:  1.9906  on 61  degrees of freedom
AIC: -24.348
Number of Fisher Scoring iterations: 5
```

```
Analysis of Deviance Table
Model: Gamma, link: log
Response: od
Terms added sequentially (first to last)

                  Df Deviance Resid. Df Resid. Dev  Pr(>Chi)
NULL                                65    29.3928
time               1   2.7653        64    26.6275 < 2.2e-16 ***
I(time^2)          1   0.4847        63    26.1428 6.181e-05 ***
infstat            1   8.4513        62    17.6915 < 2.2e-16 ***
I(time^2):infstat  1  15.7008        61     1.9906 < 2.2e-16 ***
---
Signif. codes:  0 '***' 0.001 '**' 0.01 '*' 0.05 '.' 0.1 ' ' 1
```

##### Maximum growth rates (**Table S3**)

```
Welch Two Sample t-test
t = -1.1597, df = 3.833, p-value = 0.3133
alternative hypothesis: true difference in means is not equal to 0
95 percent confidence interval:
 -0.17136979  0.07161507
sample estimates:
mean of x mean of y
 1.547743  1.597621
```

##### Maximum decline rate & times at max decline rate (**Table S3**)

```
Welch Two Sample t-test 2h vs 6h max lysis rate
t = -0.88417, df = 9.9911, p-value = 0.3974
alternative hypothesis: true difference in means is not equal to 0
95 percent confidence interval:
 -1.2577900  0.5432068
sample estimates:
 mean of x  mean of y
-0.5430675 -0.1857758
```

##### % decrease OD (**Table S3**)

```
Wilcoxon rank sum test 2h vs 6h percentage decrease in OD
W = 0, p-value = 0.1
alternative hypothesis: true location shift is not equal to 0
```

---

#### *Fig. S6b*

##### CFU at infection

```
Welch Two Sample t-test (panel)
t = 1.1225, df = 3.6483, p-value= 0.33
alternative hypothesis: true difference in means is not equal to 0
95 percent confidence interval:
 -26171778  59505111
sample estimates:
mean of x mean of y
 97666667  81000000
```

---

#### *Fig. S6c*

##### PFU at end of 2h infection period

```
Wilcoxon rank sum test (panel)
W = 9, p-value = 0.1
alternative hypothesis: true location shift is not equal to 0
```

---

#### Supplementary figure 6 (Fig. S6)

##### *Fig. S6d*

##### OD~time*infection status relationship

**Model 0-10h (panel)**

```
Call:
glm(formula = od ~ time + I(time^2) * infstat, family =
gaussian(link = "1/mu^2"))

Deviance Residuals:
     Min        1Q    Median        3Q       Max
-0.97049  -0.36169   0.03407   0.41096   0.68225

Coefficients:
                       Estimate Std. Error t value Pr(>|t|)
(Intercept)           0.0674220  0.0061107  11.033 3.62e-16 ***
time                  0.0002725  0.0029526   0.092  0.92678
I(time^2)             0.0003315  0.0003220   1.030  0.30723
infstatinf           -0.0023511  0.0064301  -0.366  0.71590
I(time^2):infstatinf  0.0010557  0.0002687   3.929  0.00022 ***
---
Signif. codes:  0 ‘***’ 0.001 ‘**’ 0.01 ‘*’ 0.05 ‘.’ 0.1 ‘ ’ 1

(Dispersion parameter for gaussian family taken to be 0.1887571)
Null deviance: 28.701  on 65  degrees of freedom
Residual deviance: 11.514  on 61  degrees of freedom
AIC: 84.056
Number of Fisher Scoring iterations: 5
```

**Analysis of Deviance Table**

```
Model: gaussian, link: 1/mu^2
Response: od
Terms added sequentially (first to last)

                  Df Deviance Resid. Df Resid. Dev  Pr(>Chi)
NULL                                65     28.701
time               1  11.1840       64     17.517 1.388e-14 ***
I(time^2)          1   0.9000       63     16.617  0.028996 *
infstat            1   1.4030       62     15.214  0.006405 **
I(time^2):infstat  1   3.7006       61     11.514 9.520e-06 ***
---
Signif. codes:  0 ‘***’ 0.001 ‘**’ 0.01 ‘*’ 0.05 ‘.’ 0.1 ‘ ’ 1
```

**Model 26-30h (panel)**

```
Call:
lm(formula = od ~ time + infstat)

Residuals:
     Min       1Q   Median       3Q      Max
-0.31950 -0.08804  0.00375  0.06499  0.29450

Coefficients:
            Estimate Std. Error t value Pr(>|t|)
(Intercept)  2.02522    0.73254   2.765   0.0145 *
time        -0.02008    0.02607  -0.770   0.4531
infstatinf  -0.85356    0.08516 -10.023 4.85e-08 ***
---
Signif. codes:  0 ‘***’ 0.001 ‘**’ 0.01 ‘*’ 0.05 ‘.’ 0.1 ‘ ’ 1

Residual standard error: 0.1806 on 15 degrees of freedom
Multiple R-squared:  0.8708,  Adjusted R-squared:  0.8535
F-statistic: 50.53 on 2 and 15 DF,  p-value: 2.165e-07
```

**Analysis of Variance Table**

```
Response: od
          Df Sum Sq Mean Sq  F value    Pr(>F)
time       1 0.0194  0.0194   0.5933    0.4531
infstat    1 3.2785  3.2785 100.4690 4.845e-08 ***
Residuals 15 0.4895  0.0326
---
Signif. codes:  0 ‘***’ 0.001 ‘**’ 0.01 ‘*’ 0.05 ‘.’ 0.1 ‘ ’ 1
```

##### Maximum decline rates & times at max decline rates (**Table S3**)

**Model max decline rate**

```
Call:
lm(formula = maxdecrt ~ infstat * tpd)

Residuals:
      Min        1Q    Median        3Q       Max
-0.093836 -0.007693  0.005792  0.016177  0.066387

Coefficients:
                  Estimate Std. Error t value Pr(>|t|)
(Intercept)       -0.22321    0.02664  -8.379 3.13e-05 ***
infstatnon         0.18565    0.03768   4.928 0.001153 **
tpd30              0.22100    0.03768   5.866 0.000376 ***
infstatnon:tpd30 -0.21642    0.05328  -4.062 0.003625 **
---
Signif. codes:  0 ‘***’ 0.001 ‘**’ 0.01 ‘*’ 0.05 ‘.’ 0.1 ‘ ’ 1

Residual standard error: 0.04614 on 8 degrees of freedom
Multiple R-squared:  0.8427,  Adjusted R-squared:  0.7838
F-statistic: 14.29 on 3 and 8 DF,  p-value: 0.001407
```

**Analysis of Variance Table**

```
Response: maxdecrt
            Df   Sum Sq  Mean Sq F value   Pr(>F)
infstat      1 0.017993 0.017993  8.4506 0.019680 *
tpd          1 0.038164 0.038164 17.9246 0.002862 **
infstat:tpd  1 0.035127 0.035127 16.4980 0.003625 **
Residuals    8 0.017033 0.002129
---
Signif. codes:  0 ‘***’ 0.001 ‘**’ 0.01 ‘*’ 0.05 ‘.’ 0.1 ‘ ’ 1
```

**Simultaneous Tests for General Linear Hypotheses**

```
infstat = inf:
 contrast     estimate     SE df t.ratio p.value
 0-10 - 26-30 -0.22100 0.0377  8  -5.866  0.0004

infstat = non:
 contrast     estimate     SE df t.ratio p.value
 0-10 - 26-30 -0.00458 0.0377  8  -0.122  0.9062

------

tpd = 0-10:
 contrast  estimate     SE df t.ratio p.value
 inf - non  -0.1857 0.0377  8  -4.928  0.0012

tpd = 26-30:
 contrast  estimate     SE df t.ratio p.value
 inf - non   0.0308 0.0377  8   0.817  0.4378
```

**Model time at max decline rate**

```
Call:
lm(formula = maxdecrt_t ~ infstat + tpd)

Residuals:
    Min      1Q  Median      3Q     Max
-5.7500 -0.4583  0.4167  0.7917  2.2500

Coefficients:
              Estimate Std. Error t value Pr(>|t|)
(Intercept)      9.250      1.156   8.004 2.21e-05 ***
infstatnon      -1.500      1.334  -1.124     0.29
tpd30           17.833      1.334  13.363 3.06e-07 ***
---
Signif. codes:  0 ‘***’ 0.001 ‘**’ 0.01 ‘*’ 0.05 ‘.’ 0.1 ‘ ’ 1

Residual standard error: 2.311 on 9 degrees of freedom
Multiple R-squared:  0.9523,  Adjusted R-squared:  0.9418
F-statistic: 89.92 on 2 and 9 DF,  p-value: 1.126e-06
```

**Analysis of Variance Table**

```
Response: maxdecrt_t
          Df Sum Sq Mean Sq  F value    Pr(>F)
infstat    1   6.75    6.75   1.2634    0.2901
tpd        1 954.08  954.08 178.5806 3.064e-07 ***
Residuals  9  48.08    5.34
---
Signif. codes:  0 ‘***’ 0.001 ‘**’ 0.01 ‘*’ 0.05 ‘.’ 0.1 ‘ ’ 1
```

**Simultaneous Tests for General Linear Hypotheses**

```
Multiple Comparisons of Means: Tukey Contrasts
Fit: lm(formula = maxdecrt_t ~ infstat + tpd)

Linear Hypotheses:
contrast           Estimate Std. Error t value Pr(>|t|)
26-30 - 0-10 == 0    17.833      1.334   13.36 3.06e-07 ***
---
Signif. codes:  0 ‘***’ 0.001 ‘**’ 0.01 ‘*’ 0.05 ‘.’ 0.1 ‘ ’ 1
(Adjusted p values reported -- single-step method)
```

#### Supplementary figure 6 (Fig. S6)

##### Time to lysis (**Table S3**)

Not determined – rate already negative from start

**Model**

```
Call:
lm(formula = percdec ~ infstat + tpd)

Residuals:
    Min      1Q  Median      3Q     Max 
-9.0889 -3.3803 -0.6539  1.2317 16.0094 

Coefficients:
              Estimate Std. Error t value Pr(>|t|)    
(Intercept)     49.146      3.537  13.894 2.19e-07 ***
infstatnon     -24.857      4.084  -6.086 0.000182 ***
tpd30           38.314      4.084   9.380 6.08e-06 ***
---
Signif. codes:  0 '***' 0.001 '**' 0.01 '*' 0.05 '.' 0.1 ' ' 1

Residual standard error: 7.074 on 9 degrees of freedom
Multiple R-squared:  0.9329,	Adjusted R-squared:  0.9179 
F-statistic: 62.51 on 2 and 9 DF,  p-value: 5.269e-06
```

**Analysis of Variance Table**

```
Response: percdec
          Df Sum Sq Mean Sq F value    Pr(>F)    
infstat    1 1853.6  1853.6  37.037 0.0001824 ***
tpd        1 4403.9  4403.9  87.993 6.078e-06 ***
Residuals  9  450.4    50.0                      
---
Signif. codes:  0 '***' 0.001 '**' 0.01 '*' 0.05 '.' 0.1 ' ' 1
```

##### % decrease OD (**Table S3**)

**Simultaneous Tests for General Linear Hypotheses**

```
tpd = 10:
 contrast  estimate   SE df t.ratio p.value
 inf - non     24.9 4.08  9   6.086  0.0002

tpd = 30:
 contrast  estimate   SE df t.ratio p.value
 inf - non     24.9 4.08  9   6.086  0.0002

------

infstat = inf:
 contrast estimate   SE df t.ratio p.value
 10 - 30     -38.3 4.08  9  -9.380  <.0001

infstat = non:
 contrast estimate   SE df t.ratio p.value
 10 - 30     -38.3 4.08  9  -9.380  <.0001
```

---

##### *Fig. S6e*

##### CFU~time*infection status relationship

**Model (panel)**

```
Call:
glm(formula = CFU ~ time.fac * infstat, family = Gamma(link = 
"log"))

Deviance Residuals: 
     Min        1Q    Median        3Q       Max  
-1.56321  -0.38076   0.00027   0.23188   1.17787  

Coefficients:
                      Estimate Std. Error t value Pr(>|t|)    
(Intercept)            20.8659     0.3639  57.341  < 2e-16 ***
time.fac6              -0.7106     0.5146  -1.381 0.186291    
time.fac10             -0.9866     0.5146  -1.917 0.073245 .  
time.fac30             -0.9781     0.5146  -1.901 0.075511 .  
infstatinf             -0.3452     0.5146  -0.671 0.511945    
time.fac6:infstatinf   -0.1955     0.7278  -0.269 0.791652    
time.fac10:infstatinf  -1.0061     0.7278  -1.382 0.185844    
time.fac30:infstatinf  -3.6080     0.7278  -4.958 0.000143 ***
---
Signif. codes:  0 '***' 0.001 '**' 0.01 '*' 0.05 '.' 0.1 ' ' 1

(Dispersion parameter for Gamma family taken to be 0.3972455)
Null deviance: 35.2323  on 23  degrees of freedom
Residual deviance:  7.9388  on 16  degrees of freedom
AIC: 985.26
Number of Fisher Scoring iterations: 6
```

**Analysis of Deviance Table**

```
Model: Gamma, link: log
Response: CFU
Terms added sequentially (first to last)
                 Df Deviance Resid. Df Resid. Dev  Pr(>Chi)    
NULL                                23     35.232              
time.fac          3   8.5000        20     26.732 8.706e-05 ***
infstat           1   8.2238        19     18.508 5.366e-06 ***
time.fac:infstat  3  10.5697        16      7.939 7.115e-06 ***
---
Signif. codes:  0 '***' 0.001 '**' 0.01 '*' 0.05 '.' 0.1 ' ' 1
```

**Simultaneous Tests for General Linear Hypotheses**

```
infstat = non:
 contrast ratio     SE df null t.ratio p.value
 1 / 6    2.035  1.047 16    1   1.381  0.5283
 1 / 10   2.682  1.380 16    1   1.917  0.2600
 1 / 30   2.659  1.369 16    1   1.901  0.2666
 6 / 10   1.318  0.678 16    1   0.536  0.9489
 6 / 30   1.307  0.672 16    1   0.520  0.9531
 10 / 30  0.992  0.510 16    1  -0.016  1.0000

infstat = inf:
 contrast ratio     SE df null t.ratio p.value
 1 / 6    2.475  1.274 16    1   1.761  0.3269
 1 / 10   7.335  3.775 16    1   3.872  0.0066
 1 / 30  98.118 50.493 16    1   8.912  <.0001
 6 / 10   2.964  1.525 16    1   2.111  0.1915
 6 / 30  39.648 20.403 16    1   7.151  <.0001
 10 / 30 13.376  6.884 16    1   5.040  0.0006

P value adjustment: tukey method for comparing a family of 4 
estimates 
Tests are performed on the log scale 

------

time.fac = 1:
 contrast  ratio     SE df null t.ratio p.value
 non / inf  1.41  0.727 16    1   0.671  0.5119

time.fac = 6:
 contrast  ratio     SE df null t.ratio p.value
 non / inf  1.72  0.884 16    1   1.051  0.3090

time.fac = 10:
 contrast  ratio     SE df null t.ratio p.value
 non / inf  3.86  1.988 16    1   2.626  0.0184

time.fac = 30:
 contrast  ratio     SE df null t.ratio p.value
 non / inf 52.10 26.813 16    1   7.682  <.0001

Tests are performed on the log scale 
```

---

### Supplementary figure 6 (Fig. S6)

#### *Fig. S6f*

#### PFU~time*population relationship

```
Model (panel)
Call:
glm(formula = pfu_mL ~ time + I(time^2) + ppn, family =
inverse.gaussian(link = "log"))

Deviance Residuals:
       Min          1Q      Median          3Q         Max
-2.634e-05  -8.498e-06  -4.364e-06   1.848e-06   2.098e-05

Coefficients:
               Estimate Std. Error t value Pr(>|t|)
(Intercept)   19.915019   0.116387 171.110  < 2e-16 ***
time           0.007909   0.103461   0.076    0.940
I(time^2)      0.019655   0.012235   1.607    0.124
ppntotal       1.050220   0.190842   5.503 2.19e-05 ***
---
Signif. codes:  0 ‘***’ 0.001 ‘**’ 0.01 ‘*’ 0.05 ‘.’ 0.1 ‘ ’ 1

(Dispersion parameter for inverse.gaussian family taken to be
1.470364e-10)
Null deviance: 1.7284e-08  on 23  degrees of freedom
Residual deviance: 3.4084e-09  on 20  degrees of freedom
AIC: 1045.2
Number of Fisher Scoring iterations: 3
```

```
Analysis of Deviance Table
Model: inverse.gaussian, link: log
Response: pfu_mL
Terms added sequentially (first to last)

            Df   Deviance Resid. Df Resid. Dev  Pr(>Chi)
NULL                             23 1.7284e-08
time         1 8.1778e-09        22 9.1060e-09 8.803e-14 ***
I(time^2)    1 1.3310e-10        21 8.9729e-09    0.3414
ppn          1 5.5644e-09        20 3.4084e-09 7.664e-10 ***
---
Signif. codes:  0 ‘***’ 0.001 ‘**’ 0.01 ‘*’ 0.05 ‘.’ 0.1 ‘ ’ 1


Simultaneous Tests for General Linear Hypotheses
Multiple Comparisons of Means: Tukey Contrasts
Fit: glm(formula = pfu_mL ~ time + I(time^2) + ppn, family =
inverse.gaussian(link = "log"))

Linear Hypotheses:
contrast          Estimate Std. Error z value Pr(>|z|)
total - free == 0   1.0502     0.1908   5.503 3.73e-08 ***
---
Signif. codes:  0 ‘***’ 0.001 ‘**’ 0.01 ‘*’ 0.05 ‘.’ 0.1 ‘ ’ 1
(Adjusted p values reported -- single-step method)
```

#### Maximum PFU rate & time at max PFU rate (**Table S3**)

```
Welch Two Sample t-test (Max PFU production rate)
t = 0.74342, df = 3.9797, p-value = 0.4987
alternative hypothesis: true difference in means is not equal to 0
95 percent confidence interval:
 -0.2030296  0.3511067
sample estimates:
mean of x mean of y
0.3088986 0.2348600

Wilcoxon rank sum test with continuity correction (Time at max PFU production rate)
W = 3, p-value = 0.6193
alternative hypothesis: true location shift is not equal to 0
```

#### Fold change in PFU (**Table S3**)

```
Model (Table S3)
Call:
glm(formula = fch ~ ppn + tpd, family = Gamma(link = log))

Deviance Residuals:
    Min       1Q   Median       3Q      Max
-1.2296  -0.7454  -0.4358   0.4026   1.1852

Coefficients:
              Estimate Std. Error t value Pr(>|t|)
(Intercept)     0.6625     0.4235   1.565    0.152
ppntotal        0.6034     0.4890   1.234    0.248
tpd30           0.3490     0.4890   0.714    0.494

(Dispersion parameter for Gamma family taken to be 0.7172901)
Null deviance: 7.7261  on 11  degrees of freedom
Residual deviance: 6.6501  on  9  degrees of freedom
AIC: 56.747
Number of Fisher Scoring iterations: 9
```

```
Analysis of Deviance Table
Model: Gamma, link: log
Response: fch
Terms added sequentially (first to last)
         Df Deviance Resid. Df Resid. Dev Pr(>Chi)
NULL                        11     7.7261
ppn       1  0.74417        10     6.9819   0.3084
tpd       1  0.33179         9     6.6501   0.4964
```

#### Average burst sizes (**Table S3**)

ABS; average burst size

```
Wilcoxon rank sum test (Table S3)
W = 1, p-value = 0.2
alternative hypothesis: true location shift is not equal to 0
```

==================================================================
