## Supplementary material for "Persistent lytic bacteriophage infection as a novel strategy for exploitation of nutrient-limited host bacteria": Table_S3

**Table S3. Supporting information on bacterial growth, phage induced lysis, and PFU production for main Figs. 1 to 4 and 6 and to Supplementary figures 2, 5, and 6 (abbreviated S2, S5 and S6)**

| Figure | Panel | Treatment / condition | Growth rate at infection / measurement ($h^{-1}$) [a] | Maximum growth rate ($h^{-1}$) | Time at max growth rate (h) [b] | Time to initiation of lysis (h) [c] | Maximum decline rate ($h^{-1}$) [d] | Time at max decline rate (h) [b] | Time to reach 50% maximum $OD_{600}$ (h) [e] | Duration of infection (h) [f] | % decrease in $OD_{600}$ [g] | % culture regrowth [h] | Maximum rate of PFU production ($h^{-1}$) | Time at max PFU production rate (h) [b] | Fold-change in PFU [i] | Average burst size (PFU $CFU^{-1}$) [j] |
|---|---|---|---|---|---|---|---|---|---|---|---|---|---|---|---|---|
| 1 | b | EP | 1.25 ± 0.129 A | NA | NA | 0.833 ± 0.144 | -1.32 ± 0.293 A | 1.00 ± 0.00 A | ND | NA | 50.6 ± 14.9 A | NA | NA | NA | NA | NA |
| | | eTP | 0.392 ± 0.0855 B | NA | NA | 0.667 ± 0.144 | -1.34 ± 0.493 A | 1.00 ± 0.00 A | ND | NA | 54.2 ± 6.88 A | NA | NA | NA | NA | NA |
| | | lTP | 0.0651 ± 0.00555 C | NA | NA | ND | -0.104 ± 0.147 B | 1.42 ± 0.577 A | ND | NA | 5.30 ± 2.32 B | NA | NA | NA | NA | NA |
| | | SP | -0.0364 ± 0.0785 C | NA | NA | ND | -0.175 ± 0.206 B | 1.25 ± 0.867 A | ND | NA | 20.1 ± 9.80 B | NA | NA | NA | NA | NA |
| 1 | c | EP | NA | NA | NA | NA | NA | NA | NA | NA | NA | NA | NA | NA | 24.0 (20.1, 31.3) A | NA |
| | | eTP | NA | NA | NA | NA | NA | NA | NA | NA | NA | NA | NA | NA | 7.20 (4.65, 7.23) B | NA |
| | | lTP | NA | NA | NA | NA | NA | NA | NA | NA | NA | NA | NA | NA | 0.845 (0.490, 0.975) C | NA |
| | | SP | NA | NA | NA | NA | NA | NA | NA | NA | NA | NA | NA | NA | 0.650 (0.390, 0.655) C | NA |
| 1 | d | Non-infected | ND | NA | NA | ND | -0.0922 ± 0.0212 A | 10.0 (10.0, 10.1) A | ND | ND | 50.8 (50.5, 51.6) A | NA | NA | NA | NA | NA |
| | | Infected | ND | NA | NA | ND | -0.212 ± 0.0377 B | 28.0 (28.0, 28.0) B | ND | ND | 90.0 (89.8, 90.7) B | NA | NA | NA | NA | NA |
| 1 | e | Infected (2h p.i.) | NA | NA | NA | NA | NA | NA | NA | NA | NA | NA | NA | NA | 0.325 ± 0.0926 A | NA |
| | | Infected (18h p.i.) | NA | NA | NA | NA | NA | NA | NA | NA | NA | NA | NA | NA | 5.90 ± 2.65 B | NA |
| 2 | a | Non-infected | NA | 1.04 ± 0.0424 | 2.33 ± 0.00 | NA | -0.0458 (-0.0517, -0.0430) A | 20.1 (20.0, 20.5) A | ND | NA | 32.8 (32.7, 33.8) A | NA | NA | NA | NA | NA |
| 2 | a,b | Infected (EP) | 0.709 (0.700, 0.722) A | NA | NA | 0.528 ± 0.0482 A | -1.184 (-1.244, -1.178) B | 4.58 (4.54, 4.71) B | 0.250 (0.250, 0.250) A | 6.92 (6.79, 7.63) A | 84.5 (83.2, 84.6) B | NA | NA | NA | NA | NA |
| | | Infected (eTP, p1) | 0.122 (0.120, 0.123) B | NA | NA | 1.08 ± 0.221 B | -0.105 (-0.131, -0.0979) C | 7.67 (7.63, 8.58) C | 8.33 (6.83, 10.9) B | 17.7 (17.2, 18.8) B | 77.4 (77.0, 79.8) C | NA | NA | NA | NA | NA |
| | | Infected (eTP, p2) | NA | NA | NA | NA | -0.173 (-0.182, -0.172) CD | 10.9 (10.9, 10.9) D | NA | NA | NA | NA | NA | NA | NA | NA |
| | | Infected (eTP, p3) | NA | NA | NA | NA | -0.0892 (-0.113, -0.0824) CD | 13.3 (13.2, 13.3) E | NA | NA | NA | NA | NA | NA | NA | NA |
| | | Infected (eTP, p4) | NA | NA | NA | NA | -0.142 (-0.179, -0.122) CD | 20.1 (19.6, 20.1) A | NA | NA | NA | NA | NA | NA | NA | NA |
| | | Infected (lTP, p1) | 0.0922 (0.0861, 0.0957) B | NA | NA | 1.78 ± 0.555 B | -0.131 (-0.147, -0.0938) C | 13.2 (12.9, 14.1) E | 11.2 (10.9, 12.5) B | 18.0 (17.8, 18.8) B | 83.9 (83.4, 84.5) B | NA | NA | NA | NA | NA |
| | | Infected (lTP, p2) | NA | NA | NA | NA | -0.203 (-0.219, -0.194) CD | 21.4 (21.1, 21.7) A | NA | NA | NA | NA | NA | NA | NA | NA |
| | | Infected (SP, p1) | 0.0337 (0.0323, 0.0418) C | NA | NA | 1.86 ± 0.394 B | -0.147 (-0.154, -0.127) CD | 21.3 (21.2, 21.6) A | 11.7 (11.5, 12.3) B | 17.42 (16.9, 17.7) B | 83.2 (83.2, 83.3) B | NA | NA | NA | NA | NA |
| | | Infected (SP, p2) | NA | NA | NA | NA | -0.277 (-0.297, -0.274) D | 25.4 (24.8, 25.5) F | NA | NA | NA | NA | NA | NA | NA | NA |
| 3 | a | Non-infected (0-10h) | NA | NA | NA | NA | -0.0909 ± 0.0243 A | 2.00 ± 2.00 A | ND | NA | 62.4 ± 7.17 A | NA | NA | NA | NA | NA |
| | | Infected (0-10h) | NA | NA | NA | ND | -0.132 ± 0.0289 A | 5.25 ± 3.86 A | ND | ND | 40.3 ± 2.88 B | NA | NA | NA | NA | NA |
| | | Non-infected (26-30h) | NA | NA | NA | NA | 0.01075 ± 0.0243 B | 28.0 ± 0.00 B | ND | NA | 86.1 ± 6.98 C | NA | NA | NA | NA | NA |
| | | Infected (26-30h) | NA | NA | NA | ND | 0.00604 ± 0.029 B | 28.0 ± 0.00 B | ND | ND | 47.5 ± 9.83 B | NA | NA | NA | NA | NA |
| 3 | c | Infected (0-10h, free phages) | NA | NA | NA | NA | NA | NA | NA | NA | NA | NA | 0.318 (0.276, 0.545) A | 4.5 ± 1.91 A | 1.17 ± 0.414 A | 5.80 ± 3.85 A |
| | | Infected (0-10h, total phages) | NA | NA | NA | NA | NA | NA | NA | NA | NA | NA | 0.23 (0.169, 0.281) A | 6.5 ± 2.52 A | 2.51 ± 0.933 B | NA |
| | | Infected (26-30h, free phages) | NA | NA | NA | NA | NA | NA | NA | NA | NA | NA | NA | NA | 3.26 ± 0.804 C | 50.9 ± 24.7 B |
| | | Infected (26-30h, total phages) | NA | NA | NA | NA | NA | NA | NA | NA | NA | NA | NA | NA | 3.40 ± 0.747 C | NA |
| 4 | b | Non-infected ($Bs^{wt}$) | -0.0705 ± 0.000906 A | 0.821 ± 0.00715 A | 1.94 ± 0.0962 A | NA | -0.0809 (-0.0839, -0.0808) A | 19.3 (19.3, 19.3) A | 34.7 (17.3, 36.7) A | NA | 43.5 (43.5, 45.7) A | 0.228 (-1.80, 1.50) A | NA | NA | NA | NA |
| | | Non-infected ($Bs^{Pspac::yueB}$) | -0.052 ± 0.0111 B | 0.781 ± 0.0205 A | 2.11 ± 0.0962 A | NA | -0.0319 (-0.0391, -0.0317) A | 27.5 (22.5, 27.5) B | ND | NA | 43.5 (41.5,45.1) A | 0.00 (0.00, 3.92) A | NA | NA | NA | NA |
| | | Infected ($Bs^{wt}$, p1) | -0.0741 ± 0.00137 A | NA | NA | ND | -0.0837 (-0.0855, -0.0631) A | 17.2 (15.7, 18.5) A | 23 (22.8, 23.2) B | 18 (18, 18) A | 89.3 (89.0, 89.4) B | 0.644 (0.540, 1.02) A | NA | NA | NA | NA |
| | | Infected ($Bs^{wt}$, p2) | NA | NA | NA | ND | -0.326 (-0.326, -0.326) B | 26.2 (26.2, 26.2) B | NA | NA | NA | NA | NA | NA | NA | NA |
| | | Infected ($Bs^{Pspac::yueB}$, p1) | -0.0495 ± 0.000803 B | NA | NA | ND | -0.117 (-0.165, -0.103) A | 15.8 (15.7, 16.3) A | 17.8 (17.4, 18.5) C | 10.5 (10.5, 10.5) B | 88.5 (87.8, 88.8) B | 28.0 (25.1, 31.0) B | NA | NA | NA | NA |
| | | Infected ($Bs^{Pspac::yueB}$, p2) | NA | NA | NA | ND | -0.46 (-0.460, -0.459) C | 19.0 (19.0, 19.0) A | NA | NA | NA | NA | NA | NA | NA | NA |
| 4 | c | Non-infected (0-10h, $Bs^{wt}$) | NA | NA | NA | NA | -0.0487 (-0.0491, -0.0479) A | 1.00 (1.00, 4.00) A | ND | NA | 39 ± 2.35 A | NA | NA | NA | NA | NA |
| | | Non-infected (0-10h, $Bs^{Pspac::yueB}$) | NA | NA | NA | NA | -0.0696 (-0.0751, -0.0601) A | 1.00 (1.00, 5.00) A | ND | NA | 36.7 ± 4.41 A | NA | NA | NA | NA | NA |
| | | Infected (0-10h, $Bs^{wt}$) | NA | NA | NA | ND | -0.104 (-0.106, -0.100) B | 10.0 (9.00, 10.0) B | ND | ND | 57.3 ± 7.08 B | NA | NA | NA | NA | NA |
| | | Infected (0-10h, $Bs^{Pspac::yueB}$) | NA | NA | NA | ND | -0.422 (-0.430, -0.270) C | 10.0 (9.00, 10.0) B | ND | ND | 77.8 ± 18.08 C | NA | NA | NA | NA | NA |
| | | Non-infected (26-30h, $Bs^{wt}$) | NA | NA | NA | NA | -0.0233 (-0.0236, -0.0171) A | 28.0 (28.0, 28.0) A | ND | NA | 65.0 (65.0, 67.0) A | NA | NA | NA | NA | NA |
| | | Non-infected (26-30h, $Bs^{Pspac::yueB}$) | NA | NA | NA | NA | -0.0106 (-0.0199, 0.000189) B | 28.0 (28.0, 29.0) A | ND | NA | 65.0 (55.0, 67.0) A | NA | NA | NA | NA | NA |
| | | Infected (26-30h, $Bs^{wt}$) | NA | NA | NA | ND | -0.0202 (-0.0322, -0.0110) A | 28.0 (28.0, 29.0) A | ND | ND | 90 (90.0, 91.0) B | NA | NA | NA | NA | NA |
| | | Infected (26-30h, $Bs^{Pspac::yueB}$) | NA | NA | NA | ND | 0.0904 (0.0847, 0.140) C | 28.0 (28.0, 28.0) A | ND | ND | 93.0 (93.0, 93.0) B | NA | NA | NA | NA | NA |
| 4 | d | Infected (0-10h, $Bs^{wt}$, total phages) | NA | NA | NA | NA | NA | NA | NA | NA | NA | NA | 0.249 ± 0.086 A | 4.00 (2.50, 5.00) A | 2.47 ± 1.13 A | ND |
| | | Infected (0-10h, $Bs^{Pspac::yueB}$, total phages) | NA | NA | NA | NA | NA | NA | NA | NA | NA | NA | 1.236 ± 0.789 B | 1.00 (1.00, 1.00) A | 6.00 ± 1.78 A | ND |
| | | Infected (26-30h, $Bs^{wt}$, total phages) | NA | NA | NA | NA | NA | NA | NA | NA | NA | NA | NA | NA | 5.70 ± 1.54 A | ND |
| | | Infected (26-30h, $Bs^{Pspac::yueB}$, total phages) | NA | NA | NA | NA | NA | NA | NA | NA | NA | NA | NA | NA | 3.48 ± 1.38 A | ND |
| 4 | e | Infected (0-10h, $Bs^{wt}$, free phages) | NA | NA | NA | NA | NA | NA | NA | NA | NA | NA | 0.614 ± 0.338 C | 2.00 (1.50, 6.00) A | 2.89 ± 2.41 A | ND |
| | | Infected (0-10h, $Bs^{Pspac::yueB}$, free phages) | NA | NA | NA | NA | NA | NA | NA | NA | NA | NA | 2.265 ± 0.584 D | 1.00 (1.00, 2.50) A | 3.17 ± 3.27 A | ND |
| | | Infected (26-30h, $Bs^{wt}$, free phages) | NA | NA | NA | NA | NA | NA | NA | NA | NA | NA | NA | NA | 4.72 ± 1.06 B | ND |
| | | Infected (26-30h, $Bs^{Pspac::yueB}$, free phages) | NA | NA | NA | NA | NA | NA | NA | NA | NA | NA | NA | NA | 2.78 ± 1.28 B | ND |
| 6 | b | Non-infected (none) | NA | 0.803 ± 0.0184 | 1.89 ± 0.0962 | NA | -0.0666 (-0.0672, -0.0660) A | 27.0 (27.0, 27.0) A | NA | NA | 49.0 ± 3.53 A | 3.34 ± 2.29 A | NA | NA | NA | NA |
| | | Non-infected (0h supplemented) | NA | NA | NA | NA | ND | ND | NA | NA | 4.46 ± 2.74 B | 14.4 ± 4.99 BC | NA | NA | NA | NA |
| | | Non-infected (1h supplemented) | NA | NA | NA | NA | ND | ND | NA | NA | 7.08 ± 3.85 B | 15.3 ± 7.00 BC | NA | NA | NA | NA |
| | | Non-infected (4h supplemented) | NA | NA | NA | NA | ND | ND | NA | NA | 17.4 ± 2.28 C | 22.8 ± 2.38 C | NA | NA | NA | NA |
| | | Non-infected (8h supplemented) | NA | NA | NA | NA | ND | ND | NA | NA | 30.5 ± 4.73 D | 37.4 ± 0.918 CD | NA | NA | NA | NA |
| | | Infected (none, p1) | -0.00921 ± 0.00447 A | NA | NA | ND | -0.1264 (-0.133, -0.122) B | 19.3 (19.3, 19.3) B | 11.8 ± 0.585 A | 18.1 ± 0.347 A | 89.4 ± 0.810 E | 1.62 ± 0.970 A | NA | NA | NA | NA |
| | | Infected (none, p2) | NA | NA | NA | NA | -0.353 (-0.353, -0.353) C | 25.5 (25.5, 25.5) C | NA | NA | NA | NA | NA | NA | NA | NA |
| | | Infected (0h supplemented, p1) | -0.00249 ± 0.00456 AB | NA | NA | ND | -0.0885 (-0.0914, -0.0623) A | 18.2 (14.6, 18.3) B | 10.9 ± 0.192 AB | 14.7 ± 0.442 B | 88.4 ± 0.220 E | 40.5 ± 7.75 D | NA | NA | NA | NA |
| | | Infected (0h supplemented, p2) | NA | NA | NA | NA | -0.676 (-0.676, -0.669) D | 23.0 (23.0, 23.0) C | NA | NA | NA | NA | NA | NA | NA | NA |
| | | Infected (1h supplemented, p1) | -0.01464 ± 0.0029 B | NA | NA | ND | -0.0613 (-0.0675, -0.0563) A | 17.7 (14.3, 17.8) B | 10.7 ± 0.442 AB | 14.4 ± 0.822 B | 88.3 ± 1.15 E | 38.4 ± 9.21 D | NA | NA | NA | NA |
| | | Infected (1h supplemented, p2) | NA | NA | NA | NA | -0.6067 (-0.607, -0.605) E | 22.8 (22.8, 22.8) C | NA | NA | NA | NA | NA | NA | NA | NA |
| | | Infected (4h supplemented, p1) | -0.01751 ± 0.00401 B | NA | NA | ND | -0.1331 (-0.133, -0.127) B | 19.2 (19.1, 19.2) B | 10.1 ± 0.632 B | 13.9 ± 0.693 B | 89.1 ± 0.460 E | 40.5 ± 6.57 D | NA | NA | NA | NA |
| | | Infected (4h supplemented, p2) | NA | NA | NA | NA | -0.592 (-0.592, -0.592) E | 22.0 (22.0, 22.0) C | NA | NA | NA | NA | NA | NA | NA | NA |
| | | Infected (8h supplemented, p1) | -0.01784 ± 0.00323 B | NA | NA | ND | -0.0607 (-0.0811, -0.0607) A | 11.0 (11.0, 15.5) B | 10.6 ± 0.347 AB | 14.4 ± 0.42 B | 89.1 ± 1.21 E | 24.3 ± 21.0 C | NA | NA | NA | NA |
| | | Infected (8h supplemented, p2) | NA | NA | NA | NA | -0.6925 (-0.693, -0.692) D | 22.7 (22.7, 22.7) C | NA | NA | NA | NA | NA | NA | NA | NA |
| 6 | c | Infected (none, free phages) | NA | NA | NA | NA | NA | NA | NA | NA | NA | NA | NA | NA | 15.8 ± 5.82 A | NA |
| | | Infected (none, total phages) | NA | NA | NA | NA | NA | NA | NA | NA | NA | NA | NA | NA | 13.1 ± 3.48 A | NA |
| | | Infected (0h supplemented, free phages) | NA | NA | NA | NA | NA | NA | NA | NA | NA | NA | NA | NA | 10.1 ± 1.76 A | NA |
| | | Infected (0h supplemented, total phages) | NA | NA | NA | NA | NA | NA | NA | NA | NA | NA | NA | NA | 29.7 ± 8.75 B | NA |
| | | Infected (1h supplemented, free phages) | NA | NA | NA | NA | NA | NA | NA | NA | NA | NA | NA | NA | 9.28 ± 2.72 A | NA |
| | | Infected (1h supplemented, total phages) | NA | NA | NA | NA | NA | NA | NA | NA | NA | NA | NA | NA | 25.7 ± 8.81 B | NA |
| | | Infected (4h supplemented, free phages) | NA | NA | NA | NA | NA | NA | NA | NA | NA | NA | NA | NA | 8.95 ± 2.22 A | NA |
| | | Infected (4h supplemented, total phages) | NA | NA | NA | NA | NA | NA | NA | NA | NA | NA | NA | NA | 20.3 ± 4.80 B | NA |
| | | Infected (8h supplemented, free phages) | NA | NA | NA | NA | NA | NA | NA | NA | NA | NA | NA | NA | 8.32 ± 6.28 A | NA |
| | | Infected (8h supplemented, total phages) | NA | NA | NA | NA | NA | NA | NA | NA | NA | NA | NA | NA | 19.0 ± 4.77 B | NA |
| 6 | d | Infected (none, free phages) | NA | NA | NA | NA | NA | NA | NA | NA | NA | NA | NA | NA | 5.05 (4.53, 5.40) AB | NA |
| | | Infected (none, total phages) | NA | NA | NA | NA | NA | NA | NA | NA | NA | NA | NA | NA | 35.5 (27.3, 37.8) AC | NA |
| | | Infected (0h supplemented, free phages) | NA | NA | NA | NA | NA | NA | NA | NA | NA | NA | NA | NA | 1.75 (1.65, 2.08) AB | NA |
| | | Infected (0h supplemented, total phages) | NA | NA | NA | NA | NA | NA | NA | NA | NA | NA | NA | NA | 9.60 (7.58, 13.0) D | NA |
| | | Infected (6h supplemented, free phages) | NA | NA | NA | NA | NA | NA | NA | NA | NA | NA | NA | NA | 2.45 (1.90, 2.90) BE | NA |
| | | Infected (6h supplemented, total phages) | NA | NA | NA | NA | NA | NA | NA | NA | NA | NA | NA | NA | 7.90 (6.08, 10.2) C | NA |
| | | Infected (10h supplemented, free phages) | NA | NA | NA | NA | NA | NA | NA | NA | NA | NA | NA | NA | 4.40 (4.10, 5.88) B | NA |
| | | Infected (10h supplemented, total phages) | NA | NA | NA | NA | NA | NA | NA | NA | NA | NA | NA | NA | 7.75 (5.55, 8.15) C | NA |
| S2 | a | Non-infected | NA | 1.44 ± 0.146 | 1.00 ± 0.00 | NA | -0.0891 ± 0.023 | 11.0 ± 1.00 | NA | NA | 64.5 ± 4.98 | NA | NA | NA | NA | NA |
| S2 | b-d | Non-infected (2h) | 1.15 ± 0.0660 A | NA | NA | NA | NA | NA | NA | NA | NA | NA | NA | NA | NA | NA |
| | | Non-infected (4h) | 0.459 ± 0.0136 B | NA | NA | NA | NA | NA | NA | NA | NA | NA | NA | NA | NA | NA |
| | | Non-infected (6h) | 0.0760 ± 0.0199 C | NA | NA | NA | NA | NA | NA | NA | NA | NA | NA | NA | NA | NA |
| | | Non-infected (10h) | -0.0860 ± 0.0230 D | NA | NA | NA | NA | NA | NA | NA | NA | NA | NA | NA | NA | NA |
| | | Non-infected (30h) | -0.0182 ± 0.0155 D | NA | NA | NA | NA | NA | NA | NA | NA | NA | NA | NA | NA | NA |
| S5 | a | $Bs^{wt}$ | NA | 1.43 ± 0.236 A | 1.00 ± 0.00 A | NA | -0.0596 ± 0.0125 A | 9.00 ± 1.00 A | ND | NA | 57.1 ± 4.22 A | NA | NA | NA | NA | NA |
| | | $Bs^{Pspac::yueB}$ | NA | 1.54 ± 0.125 A | 1.00 ± 0.00 A | NA | -0.0768 ± 0.0924 A | 22.7 ± 12.7 A | ND | NA | 59.2 ± 4.87 A | NA | NA | NA | NA | NA |
| S5 | a, b | EP ($Bs^{wt}$) | 0.992 ± 0.072 A | NA | NA | NA | NA | NA | NA | NA | NA | NA | NA | NA | NA | NA |
| | | EP ($Bs^{Pspac::yueB}$) | 1.02 ± 0.0172 A | NA | NA | NA | NA | NA | NA | NA | NA | NA | NA | NA | NA | NA |
| | | eTP ($Bs^{wt}$) | 0.409 ± 0.0102 B | NA | NA | NA | NA | NA | NA | NA | NA | NA | NA | NA | NA | NA |
| | | eTP ($Bs^{Pspac::yueB}$) | 0.390 ± 0.0365 B | NA | NA | NA | NA | NA | NA | NA | NA | NA | NA | NA | NA | NA |
| | | lTP ($Bs^{wt}$) | 0.0483 ± 0.0109 C | NA | NA | NA | NA | NA | NA | NA | NA | NA | NA | NA | NA | NA |
| | | lTP ($Bs^{Pspac::yueB}$) | 0.0473 ± 0.0169 C | NA | NA | NA | NA | NA | NA | NA | NA | NA | NA | NA | NA | NA |
| | | SP ($Bs^{wt}$) | -0.0519 ± 0.0199 D | NA | NA | NA | NA | NA | NA | NA | NA | NA | NA | NA | NA | NA |
| | | SP ($Bs^{Pspac::yueB}$) | -0.0328 ± 0.0331 D | NA | NA | NA | NA | NA | NA | NA | NA | NA | NA | NA | NA | NA |
| S6 | a | Non-infected | NA | 1.60 ± 0.0579 A | 1.00 ± 0.00 A | NA | NA | NA | ND | ND | NA | NA | NA | NA | NA | NA |
| | | Infected (up to 2h post-inoculation) | NA | 1.55 ± 0.0469 A | 1.00 ± 0.00 A | 2.91 ± 0.144 | -0.81 ± 0.257 A | 4.25 ± 0.00 | ND | ND | 54.1 ± 17.9 A | NA | NA | NA | NA | NA |
| | | Infected (up to 6h post-inoculation) | NA | NA | NA | NA | -1.116 ± 0.449 A | 4.58 ± 0.72 | ND | ND | 97.3 ± 0.0263 A | NA | NA | NA | NA | NA |
| S6 | c | Infected (free phages) | NA | NA | NA | NA | NA | NA | NA | NA | NA | NA | NA | NA | 14.1 ± 1.30 A | ND |
| | | Infected (total phages) | NA | NA | NA | NA | NA | NA | NA | NA | NA | NA | NA | NA | 30.8 ± 9.75 A | ND |
| S6 | d | Non-infected (0-10h) | NA | NA | NA | ND | -0.0376 ± 0.00777 A | 7.33 ± 4.62 A | ND | NA | 22.1 ± 1.47 A | NA | NA | NA | NA | NA |
| | | Infected (0-10h) | NA | NA | NA | ND | -0.223 ± 0.0836 B | 9.67 ± 0.577 A | ND | NA | 51.3 ± 12.1 B | NA | NA | NA | NA | NA |
| | | Non-infected (26-30h) | NA | NA | NA | NA | -0.0330 ± 0.0102 AC | 26.0 ± 0.00 B | ND | NA | 64.8 ± 3.10 C | NA | NA | NA | NA | NA |
| | | Infected (26-30h) | NA | NA | NA | NA | -0.00222 ± 0.0370 C | 26.67 ± 1.16 B | ND | NA | 85.3 ± 6.28 D | NA | NA | NA | NA | NA |
| S6 | f | Infected (1-10h, free phages) | NA | NA | NA | NA | NA | NA | NA | NA | NA | NA | 0.235 ± 0.118 A | 10.0 (8.00, 10.0) A | 0.825 (0.620, 1.84) A | 5.90 ± 8.80 A |
| | | Infected (1-10h, total phages) | NA | NA | NA | NA | NA | NA | NA | NA | NA | NA | 0.309 ± 0.126 A | 6.00 (6.00, 8.00) A | 2.45 (2.13, 6.00) A | NA |
| | | Infected (26-30h, free phages) | NA | NA | NA | NA | NA | NA | NA | NA | NA | NA | NA | NA | 3.57 ± 1.84 A | 49.6 ± 35.1 A |
| | | Infected (26-30h, total phages) | NA | NA | NA | NA | NA | NA | NA | NA | NA | NA | NA | NA | 3.53 ± 2.89 A | NA |

[a] growth rate of the culture measured at the point of infection with SPP1 or at the time-point at which samples were taken or when measurements were made
[b] time at which the given maximum rates were recorded. For 'Time at max growth rate' this is the time post-inoculation. For 'Time at max decline rate' and 'Time at max PFU production rate' this is the time post-infection
[c] time from the point of infection until a switch from a positive (growth) rate to a negative (decline) rate was recorded
[d] maximum rate of decrease in $OD_{600}$. For non-infected samples, within the time period examined, these values represent a decrease in cell size with minimal cell death (see **Supplementary figure 2**) whereas for infected samples this represents phage-induced cell lysis
[e] time until 50% of the maximum $OD_{600}$ value recorded during the experiment (before $OD_{600}$ values began to decline) was reached. For non-infected samples this is time post-inoculation, for infected samples, this is time post-infection
[f] the population average of the duration of infection, calculated as the time between the point of infection and the point at which the rate of $OD_{600}$ decline dropped to below the mean $OD_{600}$ decline rate of non-infected samples
[g] the difference between the minimum and maximum $OD_{600}$ value (before $OD_{600}$ values began to decline) obtained during a given experiment, expressed as a percentage of the maximum $OD_{600}$ value
[h] the difference between the minimum $OD_{600}$ value reached during the experiment (after $OD_{600}$ values began to decline) and the final $OD_{600}$ value recorded at the end of the experiment, expressed as a percentage of the maximum $OD_{600}$ value recorded during the experiment (before $OD_{600}$ values began to decline)
[i] A description of how fold-change in PFU was calculated can be found in **Supplementary Materials and Methods**.
[j] the number of PFU released per CFU lost during the given time-period. This is a population average, and was not determined at the single cell level. A description of how average burst size was calculated can be found in **Supplementary Materials and Methods**.

Results are presented either as mean and standard deviation **[x ± x]** or median and 25% and 75% quantiles **[x (x, x)]**
Bold letters next to presented values represent Tukey's honest significant difference contrasts. Values that are not significantly different to one another are marked with the same letter. Where different colours are present this indicates separate statistical tests.
